# TDP-43 amyloid conversion in cells tracks the net charge of a polyampholyte motif, not condensate solidification

**DOI:** 10.64898/2026.10.15.759291

**Authors:** Yoon Jeong Choi, Yemuk Kang, Soojin Cho, SoYoon Jeong, Minkyo Jung, Changi Pack, Youngtae Jeong, Kiwon Song

## Abstract

Liquid-liquid phase separation of TDP-43 is implicated in ALS and FTLD, yet how its condensates convert into pathological amyloid in living cells remains poorly defined. Here we identify a polyampholyte motif (PAM) in the linker between RRM2 and the amyloid-forming low-complexity domain as a charge-tunable determinant of fibrillization that operates independently of condensate solidification. In budding yeast expressing GFP-TDP-43ΔNLS, charge reversal of three arginines (R268E/R272E/R275E; 3R-E) converted liquid puncta into ProteoStat-positive fibrils. Unexpectedly, fluorescence recovery after photobleaching showed 3R-E condensates to be more mobile than those of wild type, whereas the least mobile condensates, in yeast and in human U2OS cells, remained ProteoStat-negative. Across 24 PAM charge variants, the proportion of fibril-bearing cells scaled linearly with net charge regardless of residue identity, with charge neutralization delaying rather than abolishing conversion. Phosphomimetic substitution of the PAM serines alone induced fibrils, implicating phosphorylation as a potential physiological modulator of PAM charge.

## INTRODUCTION

Cytoplasmic inclusions of TDP-43 are a pathological hallmark of amyotrophic lateral sclerosis (ALS) and frontotemporal lobar degeneration (FTLD), where the protein is redistributed from the nucleus and deposited as hyperphosphorylated, insoluble aggregates (Neumann et al., 2006). In cells, TDP-43 undergoes liquid-liquid phase separation (LLPS) to form biomolecular condensates (Alberti and Hyman, 2021), a property shared with other proteins implicated in neurodegeneration. Phase separation of TDP-43 is driven largely by its intrinsically disordered, prion-like low-complexity domain (LCD) (Chien et al., 2021; Conicella et al., 2020), which forms a transient α-helical structure that mediates self-association and phase separation in vitro; ALS-associated mutations in this region disrupt the helix and alter phase-separation behavior (Conicella et al., 2016).

These condensates have been thought to age and convert into amyloid fibrils over time: as prion-like domains reside within a condensate, the weak, reversible, kinked β-sheet contacts (LARKS) that initially drive phase separation are gradually reinforced or replaced by more stable, cross-β-like interactions (Hughes et al., 2018). In this view, disease-associated mutations in hnRNPA1/hnRNPA2B1 accelerate maturation in vitro by strengthening a steric-zipper-forming segment within the prion-like domain (Kim et al., 2013). Consistently, condensates formed by FUS and related prion-like proteins become progressively less dynamic and acquire a fibrous morphology over time in vitro, and disease-associated mutations accelerate this transition (Molliex et al., 2015; Murakami et al., 2015; Patel et al., 2015). Solidification of the condensate was thus taken to be both the signature and the driving event of amyloid conversion, with the concentrated interior of the condensate assumed to be the site at which fibrils nucleate.

Recent studies in reconstituted systems suggest the opposite: the condensate interior is not where fibrils begin. The interfaces of condensates formed by the hnRNPA1 LCD promote fibril formation more effectively than their interiors (Linsenmeier et al., 2023). Spatially resolved rheology of FUS condensates further showed that liquid-to-solid conversion does not proceed uniformly: gelation initiates at the interface and propagates inward (Shen et al., 2023). A complementary thermodynamic analysis of the hnRNPA1 prion-like domain showed that condensates are metastable with respect to fibrils and function as kinetic sinks: their interiors suppress fibril formation, fibrils nucleate at the surface but grow from protein that has exited into the dilute phase, and ALS-associated mutations act by weakening this metastability rather than by uniformly hardening the assembly (Das et al., 2025). These conclusions rest on reconstituted systems; how a condensate becomes amyloid inside a cell is not known.

Both views nonetheless place the condensate on the path to the fibril, which makes the properties of the condensate, and the sequence features that set them, candidate determinants of conversion (Ahmad et al., 2022; Visser et al., 2024). Phase separation is described in terms of stickers and spacers, whose identity, valence and patterning determine not only whether a sequence demixes, and at what concentration (Das and Pappu, 2013; Martin et al., 2020), but also what network forms within the condensate, and hence its material state (Harmon et al., 2017; Alshareedah et al., 2021). Charged residues are prominent among such motifs. Charge influences condensation whichever sign it carries: blocks of like charge can drive demixing on their own (Nott et al., 2015; Pak et al., 2016), the acidic tracts of nucleophosmin are required for it to phase separate with arginine-rich partners (Mitrea et al., 2016), and arginine, which contributes with tyrosine to the phase separation of prion-like RNA-binding proteins (Wang et al., 2018b), determines condensate behavior in cells through its proportion and patterning (Chen et al., 2021; Miyagi et al., 2022). Charge also sets the properties of the condensate that forms: polyarginine condensates are roughly two orders of magnitude more viscous than polylysine condensates, with correspondingly slower internal exchange (Fisher and Elbaum-Garfinkle, 2020). In earlier work, we also showed that the polyanionic and polyampholyte domains of the yeast P-body protein Nst1 govern its association with P-body components and the material properties of the condensates it forms (Choi et al., 2022).

Charged clusters therefore govern condensation at several levels: whether a condensate forms, how it behaves thereafter, and the material state that results all depend on them. Charge is moreover a variable the cell itself controls, since phosphorylation alters it directly, and TDP-43 is hyperphosphorylated in disease. Beyond its LCD, TDP-43 carries several discrete clusters of charged residues — an acidic cluster in its N-terminal domain (PA1), a second acidic cluster within RRM2 (PA2), and a polyampholyte motif (PAM) in the linker between RRM2 and the LCD.

We expressed GFP-TDP-43ΔNLS in budding yeast, a system free of endogenous TDP-43 (Johnson et al., 2008), and found that charge variants of PAM converted liquid puncta into ProteoStat-positive fibrils. Loss of exchange was a consequence of this conversion rather than a precondition for it: the variants that converted formed puncta as mobile as those that did not, and the least mobile condensates we examined never converted. The same was observed in human U2OS cells. The proportion of fibril-bearing cells scaled linearly with PAM net charge irrespective of the identity of the substituted residues, and phosphomimetic substitution of the PAM serines alone was sufficient to trigger conversion.

## RESULTS

### Charged clusters outside the LCD set the material state of TDP-43 condensates without producing amyloid

Overexpression of TDP-43 in yeast cells induces the formation of puncta (Johnson et al., 2008); the same system has been used to reconstitute the aggregation of other neurodegeneration-associated proteins, including FUS (Fushimi et al., 2011; Sun et al., 2011) and α-synuclein (Outeiro and Lindquist, 2003). Because TDP-43 is otherwise retained in the nucleus, all constructs used in this study were expressed in an NLS-deficient background (K82A/R83A/K84A; hereafter ΔNLS), which confines the protein to the cytoplasm and reproduces the mislocalization seen in patients (Neumann et al., 2006; Winton et al., 2008). A sliding-window scan of the TDP-43 sequence identified three discrete charged clusters outside the C-terminal low-complexity domain (LCD): PA1 (residues 2–29, SEYIRVTEDENDEPIEIPSEDDGTVLLS) and PA2 (residues 197–220, RCTEDMTEDELREFFSQYGDVMDV), both of which are dominated by acidic (Glu/Asp) residues and are thus polyanionic, and PAM (residues 255–276, VHISNAEPKHNSNRQLERSGRF), which contains an interspersed mixture of acidic (Glu261, Glu271) and basic (Lys263, Arg268, Arg272, Arg275) residues and is therefore a polyampholyte, following the nomenclature we previously applied to the polyelectrolyte domains of the yeast P-body protein Nst1 (Choi et al., 2022). We then generated GFP-tagged TDP-43ΔNLS constructs in which each cluster was deleted individually (ΔPA1, ΔPA2 and ΔPAM), together with the two polyanionic clusters in combination (ΔPA1ΔPA2) and PAM combined with the entire LCD (ΔPAMΔLCD), and asked how each cluster contributes to puncta formation, assessing the extent and morphology of the puncta, their material properties, and the dynamics of molecules within them. Throughout this study we compared constructs by the size, number and shape of the assemblies they formed, describing the resulting GFP distributions as diffuse, microphasic assemblies (numerous and small, against a granular background), mature puncta or filamentous structures. The assemblies were characterized by three assays: sensitivity to 1,6-hexanediol, ProteoStat staining, and FRAP (Figure 1A). Cells were examined 4 and 12 h after galactose induction, within the range used for imaging and aggregation readouts in yeast TDP-43 models (Johnson et al., 2009; Park et al., 2024). Expression of every construct was confirmed by western blot after 4 h of induction (Figure S5), so all constructs were already expressed at the earlier of the two time points.

**Figure 1.**
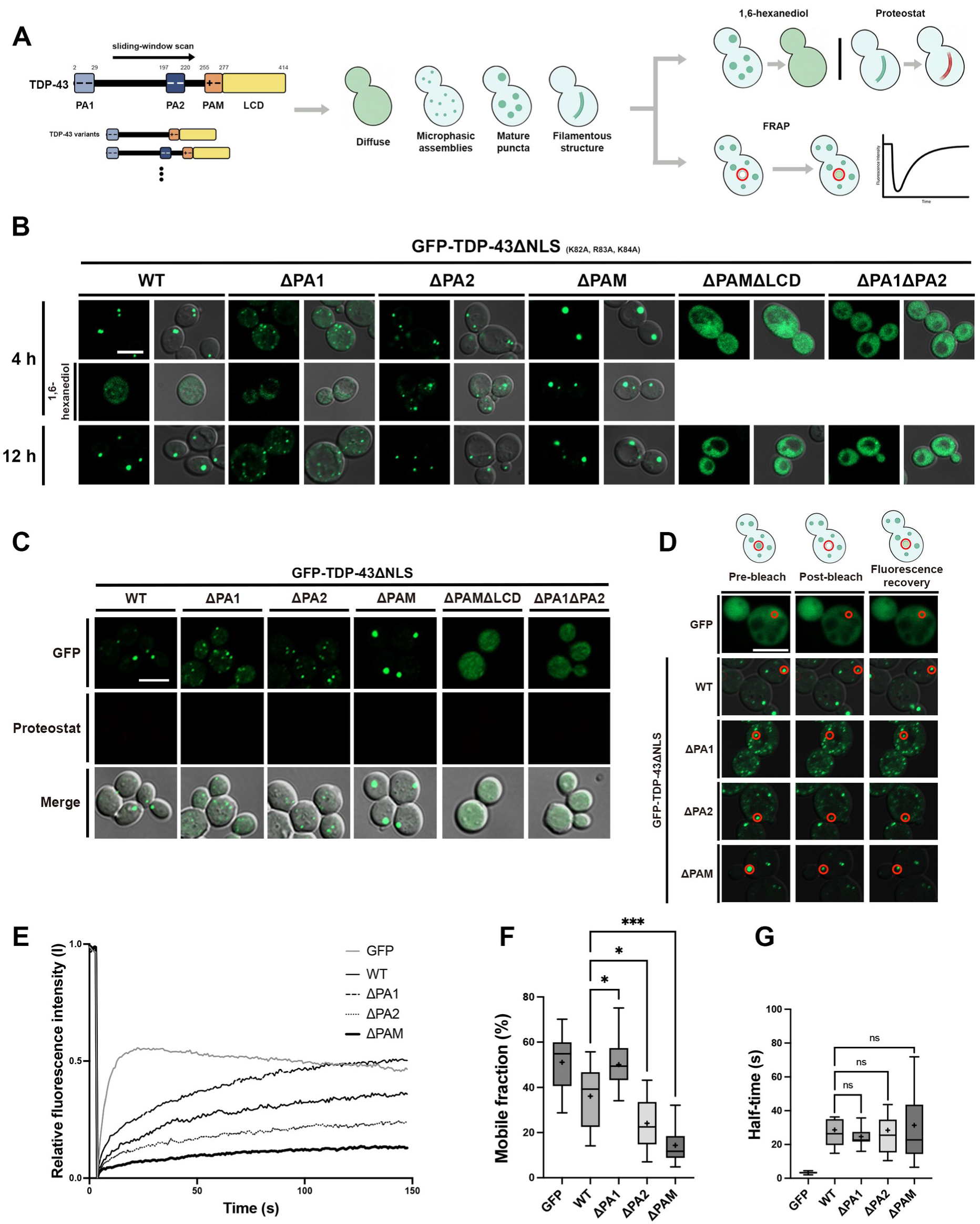
Charged clusters outside the LCD set the material state of TDP-43 condensates without producing amyloid. (A) Schematic of the experimental workflow. PA1 (polyanion 1; residues 2–29) and PA2 (polyanion 2; residues 197–220) are negatively charged clusters, whereas PAM (polyampholyte; residues 255–276) is a mixed-charge cluster; residues 277–414 are the low-complexity domain (LCD). Clusters were identified by a sliding-window scan of net charge along the sequence (arrow), and the series below the map indicates the charge-cluster variants generated on this background. NLS-deficient TDP-43 and its cluster deletion mutants (ΔPA1, ΔPA2, ΔPAM, ΔPAMΔLCD and ΔPA1ΔPA2) were overexpressed and compared by the size, number and shape of the assemblies formed — diffuse, microphasic assemblies, mature puncta or filamentous structures — then analyzed by 1,6-hexanediol treatment, ProteoStat staining and fluorescence recovery after photobleaching (FRAP). (B) Confocal images of W303a yeast cells after 4 and 12 h of overexpression of NLS-deficient GFP-TDP-43 (GFP-TDP-43ΔNLS) and its domain deletion mutants; each construct is shown as a GFP image paired with its brightfield merge. The middle row shows the same constructs at 4 h after treatment with 1,6-hexanediol; ΔPAMΔLCD and ΔPA1ΔPA2, which remain diffuse, were not treated. ΔNLS denotes replacement of three NLS residues (K82A, R83A, K84A), which confines the protein to the cytoplasm. Scale bar: 5 μm. (C) Representative confocal images of GFP-TDP-43ΔNLS and its domain deletion mutants after 4 and 12 h of galactose induction. Cells were fixed and stained with ProteoStat; GFP, ProteoStat and merged channels are shown for each of the six constructs. None gave detectable ProteoStat signal. Scale bar: 5 μm. (D) Representative confocal images of the pre-bleach, post-bleach and recovery states of condensates formed by GFP-TDP-43ΔNLS and its cluster deletion mutants ΔPA1, ΔPA2 and ΔPAM after 4 h of galactose induction. Cells expressing GFP alone were measured as a freely diffusing control. The bleached region of interest (ROI) is circled in red. Scale bar: 5 μm. (E) Average FRAP curves of GFP-TDP-43ΔNLS and its domain deletion mutants, with the GFP-only control. (F) Mobile fraction of the structures in (E), taken as the fitted plateau of a one-phase association fit and expressed as a percentage of the pre-bleach intensity. Brown-Forsythe and Welch ANOVA with Dunnett’s T3, each deletion mutant versus wild type; the GFP control is plotted for reference and was not included in the comparisons. n = 14 (WT) and 18 (GFP, ΔPA1, ΔPA2, ΔPAM) structures. *P < 0.05; ***P < 0.001. (G) Recovery half-time of the structures in (F), from the rate constant of the same fits, in seconds. n as in (F). Brown-Forsythe and Welch ANOVA with Dunnett’s T3, each deletion mutant versus wild type. ns, not significant.

Wild-type GFP-TDP-43ΔNLS formed a few discrete cytoplasmic puncta by 4 h of overexpression and retained them at 12 h (Figure 1B), as described for TDP-43 overexpressed in this system (Johnson et al., 2008). These puncta dissolved in 1,6-hexanediol and remained ProteoStat-negative through 12 h (Figure 1B, 1C), and FRAP showed that a substantial proportion of the protein within them exchanged with the cytoplasm (Figure 1D–1F). Wild type therefore reproduced the behavior reported for TDP-43 condensates in yeast, which dissolve in 1,6-hexanediol and have been described as liquid-like rather than amyloid-like (Park et al., 2017; Park et al., 2024).

Deletion of either polyanionic cluster (ΔPA1 or ΔPA2) yielded puncta that were smaller and more numerous than those of WT at both 4 and 12 h (Figure 1B). The shift was more pronounced for ΔPA1, which remained microphasic against a granular background at both time points. Deleting both clusters (ΔPA1ΔPA2) left the protein diffuse.

The two deletions nonetheless differed in the material state of the assemblies they permitted. ΔPA1 puncta dissolved in 1,6-hexanediol (Kroschwald et al., 2017), as did those of WT, whereas ΔPA2 puncta did not (Figure 1B, middle row), a behavior not previously reported for TDP-43 condensates in this system (Park et al., 2024). FRAP (Kawai-Noma et al., 2006; Pack et al., 2014) ordered them the same way: the mobile fraction of ΔPA1 was higher than that of WT (50.0% versus 36.1%, Padj = 0.0140) and that of ΔPA2 lower (24.1%, Padj = 0.0398) (Figure 1D–1F). Both deletions therefore left puncta smaller than those of WT, yet one was more dynamic than WT and the other less, so the morphology of a condensate is not a proxy for its material state.

To ask whether these phenotypes reflect the charge of the clusters rather than their physical presence, we substituted their acidic residues with asparagine or glutamine, neutralizing the charge while leaving the sequence in place. Residues were neutralized either across a whole cluster, giving the fully neutralized variants PA1 and PA2, or within individual acidic patches — groups of acidic residues that are locally concentrated but not necessarily contiguous. PA1 contains three such patches and PA2 two. For PA1 this yielded PA1-1 (E3/E9/D10/E11), PA1-3 (D13/E14/E17) and PA1-4 (E21/D22/D23), together with PA1-2, which combines PA1-3 and PA1-4; for PA2, PA2-1 (E200/D201/E204/D205/E206/E209) and PA2-2 (D216/D219) (Figure S1A).

For PA1, full neutralization left the protein completely diffuse rather than reproducing ΔPA1, and within the cluster the requirement mapped to the patches containing D13/E14/E17: PA1-2 and PA1-3 were diffuse, whereas PA1-1 and PA1-4 formed puncta (Figure S1B). This is consistent with the negatively charged N-terminal surface implicated in TDP-43 oligomerization and condensate formation (Afroz et al., 2017), and narrows the requirement within that surface, since our variants place E21 in a separate, dispensable patch. PA1 charge is thus required for puncta formation, although neutralizing it within the intact sequence is not equivalent to deleting the cluster.

For PA2, every neutralization variant formed puncta that remained hexanediol-resistant, as ΔPA2 did (Figure S1C–S1G). The two patches differ in the number of acidic residues they contain, and their effect on the mobile fraction followed that difference: PA2-1, which neutralizes six acidic residues, lowered the mobile fraction to the ΔPA2 level, whereas PA2-2, which neutralizes two, left it close to that of WT. When the constructs were ordered by the number of acidic residues neutralized — WT, PA2-2, PA2-1 and PA2 — the mobile fraction declined monotonically (*P*_trend_ < 0.0001, Jonckheere–Terpstra test; Figure S1F). Within PA2 the effect therefore tracks the amount of charge removed rather than which residues carry it, in contrast to PA1, where PA1-3 and PA1-4 neutralize three residues each yet differ in outcome (Figure S1B). The puncta themselves followed the same ordering: variants in which more acidic residues were neutralized formed smaller puncta, although every variant still formed puncta (Figure S1C). The acidic residues of PA2 therefore influence condensate maturation rather than assembly.

Deleting PAM had the opposite effect. Where loss of either polyanionic cluster left puncta that failed to mature, ΔPAM formed a small number of large, bright puncta, comparable to or larger than those of WT (Figure 1B). As in ΔPA2, these puncta did not dissolve in 1,6-hexanediol (Figure 1B, middle row), and their mobile fraction was the lowest of the three deletions — 14.3% (Padj = 0.0001), below that of ΔPA2 (Figure 1E, 1F) — so they exchanged least with the cytoplasm and retained the largest immobile fraction. The recovery half-time was indistinguishable among ΔPA1, ΔPA2, ΔPAM and wild type (Figure 1G), so each deletion altered the proportion of exchangeable molecules rather than the rate at which they exchanged. Assembly still required the LCD, the region that forms the amyloid core in patients (Arseni et al., 2022, 2023): deleting it together with PAM (ΔPAMΔLCD) abolished puncta formation at both time points. PAM is therefore dispensable for assembly, but its presence keeps the condensates that the LCD forms smaller and more dynamic. ΔPAM condensates were the least dynamic assemblies in the series, yet none of them was amyloid: they were ProteoStat-negative at both 4 and 12 h, as were those of WT, ΔPA1, ΔPA2, the neutralized variants of both polyanionic clusters, and the constructs that remained diffuse (Figure 1C; Figure S1B, C). PAM may therefore influence the material state of LCD-driven condensates without determining whether they convert.

### Charge substitution across PAM identifies 3R-E as a TDP-43 variant that forms fibril-like structures in living cells

Because deleting PAM altered the material state of the condensates the LCD forms without making them amyloid (Figure 1), we next asked how the charge carried by PAM itself (residues 255–276) governs condensate behavior. PAM is a polyampholyte cluster containing four basic residues (K263, R268, R272, R275) interspersed with two acidic residues (E261, E271). To probe this, we generated a panel of charge-substitution mutants — substituting the basic residues with neutral alanine or with negatively charged glutamate, and, conversely, substituting the acidic residues with positively charged lysine — and overexpressed each in budding yeast: K263E, which has also been reported in atypical FTLD (Kovacs et al., 2009), K263A, the triple substitution 3R-E (R268E/R272E/R275E), the corresponding neutralization 3R-A (R268A/R272A/R275A), the combined mutants K263E/3R-E and K263A/3R-A, and the reverse-charge control 2E-K (E261K/E271K) (Figure 2A).

**Figure 2.**
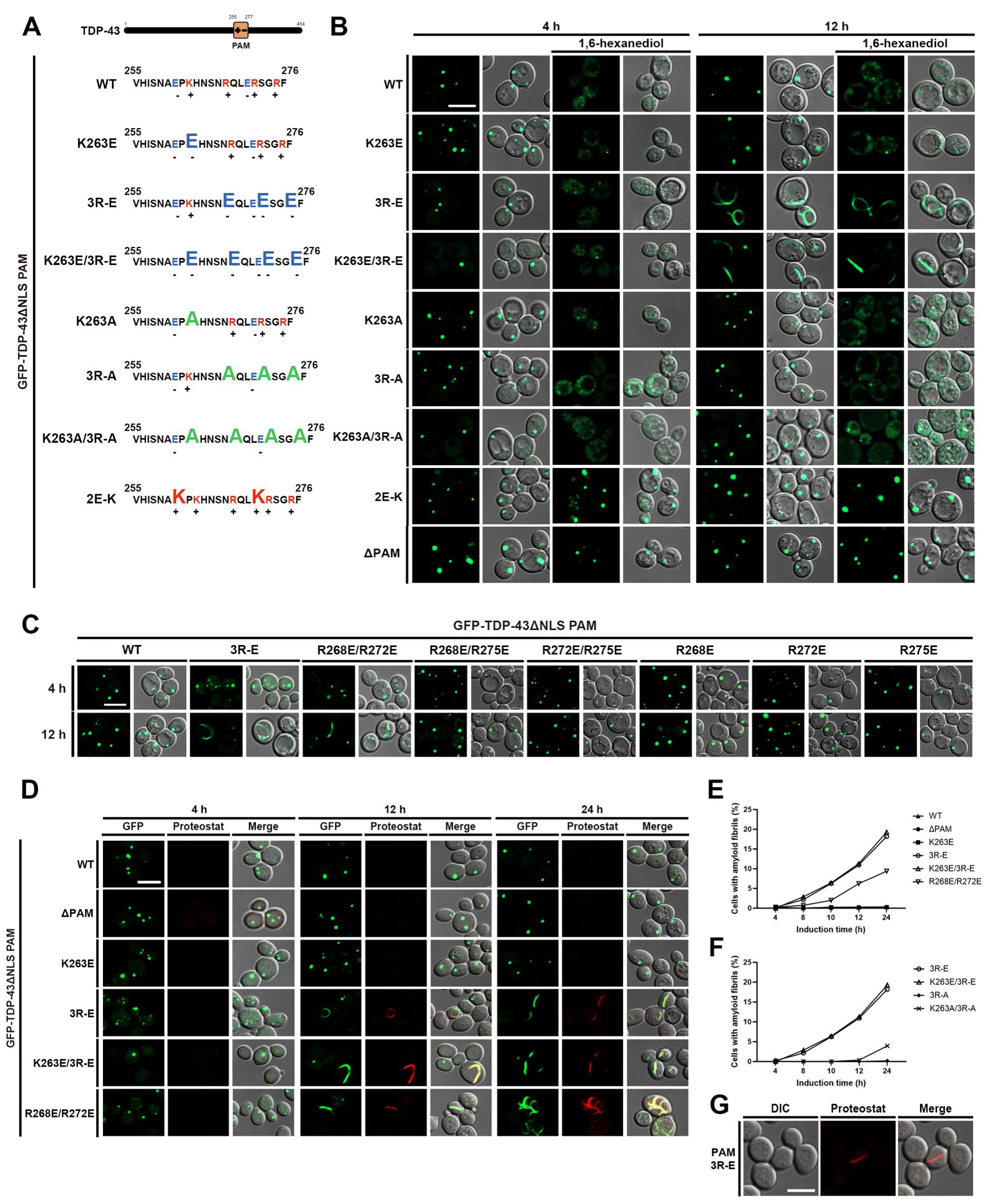
Charge substitution across PAM identifies 3R-E as a TDP-43 variant that forms fibril-like structures in living cells. (A) Schematic of the PAM cluster of TDP-43. Sequences of WT PAM and the charge-substituted mutants (K263E, 3R-E, K263E/3R-E, K263A, 3R-A, K263A/3R-A and 2E-K) are shown with substituted residues in capital letters. (B) Representative confocal images of cells expressing GFP-TDP-43ΔNLS and its PAM mutants (K263E, PAM 3R-E, K263E/3R-E, K263A, PAM 3R-A, K263A/3R-A, 2E-K and ΔPAM) after 4 or 12 h of galactose induction, each shown before and after treatment with 1,6-hexanediol at both time points. For each condition a GFP image is paired with its brightfield merge. Scale bar: 5 μm. (C) Representative confocal images of cells expressing GFP-TDP-43ΔNLS and its PAM mutants (PAM 3R-E, R268E/R272E, R268E/R275E, R272E/R275E, R268E, R272E and R275E) after 4 or 12 h of galactose induction. Scale bar: 5 μm. (D) Representative confocal images of cells expressing GFP-TDP-43ΔNLS and its PAM mutants (ΔPAM, K263E, PAM 3R-E, K263E/3R-E and R268E/R272E) after 4, 12 or 24 h of galactose induction, fixed and stained with ProteoStat. Scale bar: 5 μm. (E) Percentage of cells containing amyloid fibrils after 4, 8, 10, 12 or 24 h of galactose induction of GFP-TDP-43ΔNLS (WT) and its PAM mutants ΔPAM, K263E, PAM 3R-E, K263E/3R-E and R268E/R272E. (F) The same quantification for PAM 3R-E, K263E/3R-E, PAM 3R-A and K263A/3R-A. The PAM 3R-E and K263E/3R-E values are the scoring shown in (E), replotted for comparison with the charge-neutralized mutants. In (E) and (F), more than 300 cells per condition were scored; each point is a single pooled proportion, and no replicate-based statistics were applied. (G) Representative confocal images of cells expressing untagged, NLS-deficient TDP-43 PAM 3R-E after 12 h of galactose induction, fixed and stained with ProteoStat. DIC, ProteoStat and merged images are shown; no GFP channel is present because the construct is untagged. Scale bar: 5 μm.

All constructs formed discrete puncta by 4 h of overexpression, indistinguishable from one another and from those of WT. Their response to 1,6-hexanediol, however, already differed at this stage: the puncta of WT, K263E, 3R-E, K263E/3R-E, K263A, 3R-A and K263A/3R-A all dissolved readily, whereas those of 2E-K remained intact, resembling the hexanediol-resistant behavior of ΔPAM (Figure 2B). By 12 h of overexpression, however, the 3R-E and K263E/3R-E mutants underwent a further transition: their puncta converted into curved, fibril-like structures, and these structures now resisted 1,6-hexanediol, unlike the puncta the same mutants had formed at 4 h (Figure 2B). In contrast, K263E alone, the charge-neutralized mutants K263A, 3R-A and K263A/3R-A, and the reverse-charge control 2E-K all remained as puncta through 12 h of overexpression, with no evidence of fibril formation (Figure 2B). These results indicate that, within this time window, fibril formation by TDP-43 is driven by charge reversal of the basic residues within this cluster, and is not reproduced over 12 h by removing their charge or by reversing the charge of the acidic residues instead.

To ask how the three arginines within 3R-E (R268, R272, R275) contribute to the fibril phenotype, we constructed the three single substitutions (R268E, R272E, R275E) and the three paired substitutions (R268E/R272E, R268E/R275E, R272E/R275E) in the GFP-TDP-43ΔNLS background and compared them with 3R-E after overexpression. Among the three pairs, R268E/R272E gave the most fibrils by 12 h of overexpression (Figure 2C), and we therefore carried it forward for quantification alongside 3R-E.

We next asked whether these fibrillar structures carry amyloid character, using ProteoStat staining. WT, ΔPAM, and K263E remained ProteoStat-negative through 24 h of overexpression, whereas 3R-E, K263E/3R-E and R268E/R272E each accumulated ProteoStat-positive fibril signal over time (Figure 2D). Scoring the proportion of cells carrying amyloid fibrils over a 4–24 h time course showed that, although R268E/R272E also formed fibrils, conversion was substantially greater in 3R-E and K263E/3R-E than in R268E/R272E at every time point from 8 h onward, indicating that charge reversal of all three arginines maximizes amyloid conversion (approximately 18–19% of cells at 24 h for 3R-E and K263E/3R-E, versus approximately 9–10% for R268E/R272E; Figure 2E). Based on this, we selected 3R-E for all subsequent experiments: adding K263E increased conversion only marginally (19.3% versus 18.2% of cells at 24 h), making 3R-E the simplest substitution that achieves maximal conversion. The charge-neutralized mutants converted as well, but later and to a much smaller extent: neither 3R-A nor K263A/3R-A had formed fibrils by 12 h (Figure 2B), yet by 24 h K263A/3R-A reached 4.0% of cells and 3R-A 0.2% (Figure 2F). Removing the positive charge therefore slows conversion rather than preventing it, whereas reversing it drives conversion more strongly. Finally, overexpression of untagged PAM 3R-E, lacking the GFP tag, also produced the same fibrillar morphology and ProteoStat-positive signal, confirming that fibril formation is not an artifact of GFP tagging (Figure 2G).

### The fibril-like structures formed by PAM 3R-E coexist with puncta in single cells and are filamentous at the ultrastructural level

To characterize the fibril-like structures more closely, we acquired confocal z-stacks of cells expressing GFP-TDP-43ΔNLS PAM 3R-E at 24 h of overexpression and reconstructed them in three dimensions. The fibril-like structures were elongated and markedly curved, frequently bending back on themselves into open hooks or closed loops, and in many cells a single structure extended across much of the cell body. Rotating the reconstructions confirmed that these were continuous, thread-like objects of roughly uniform thickness along their length rather than projections of a ring-shaped punctum (Figure 3A; Videos S1–S4). Both morphologies were present within the same cell: discrete round puncta were visible alongside the filaments in these reconstructions (Figure 3A).

**Figure 3.**
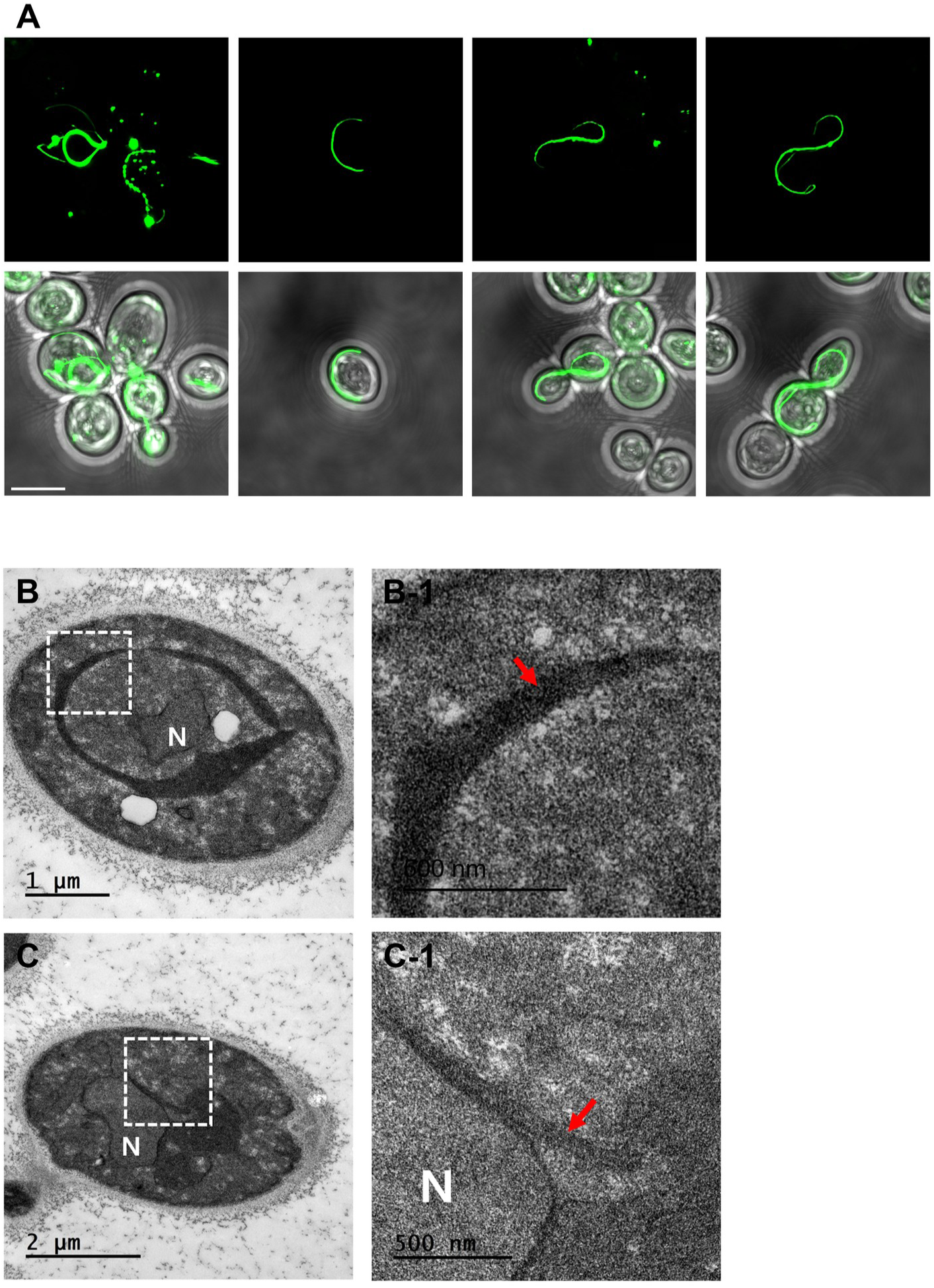
The fibril-like structures formed by PAM 3R-E coexist with puncta in single cells and are filamentous at the ultrastructural level. (A) Three-dimensional reconstructions of confocal z-stacks of W303a yeast cells expressing GFP-TDP-43ΔNLS PAM 3R-E after 24 h of galactose induction. Four representative cells are shown: upper row, the GFP channel alone; lower row, the corresponding GFP/brightfield merges. Elongated curved filaments and discrete round puncta are both present within the same cell, most clearly in the leftmost column. Rotating views of each cell are provided as Videos S1–S4. Scale bar: 5 μm. (B, C) Transmission electron micrographs of two representative yeast cells expressing GFP-TDP-43ΔNLS PAM 3R-E after 24 h of galactose induction, prepared by permanganate fixation (see Methods). Low-magnification views are shown in (B) and (C), with the nucleus marked (N) and the boxed regions enlarged in (B-1) and (C-1), respectively. Red arrows indicate filamentous structures. Scale bars: 1 μm (B), 2 μm (C) and 500 nm (B-1, C-1).

To establish whether these structures are filamentous at the ultrastructural level, we examined cells by transmission electron microscopy after 24 h of galactose induction, using permanganate fixation. At low magnification the cells retained normal overall ultrastructure, with an intact nucleus (N) and vacuole (Figure 3B, 3C). Within the cytoplasm of cells expressing PAM 3R-E, electron-dense fibrillar material was present, most often in the cytoplasmic space adjacent to the nuclear envelope (boxed regions, enlarged in Figure 3B-1 and 3C-1). At higher magnification this material appeared as unbranched, electron-dense fibrillar structures (Figure 3B-1, 3C-1, arrows). The structures formed by PAM 3R-E are therefore filaments rather than optically unresolved clusters of puncta, and their unbranched fibrillar morphology, together with their ProteoStat reactivity (Figure 2D), is consistent with an amyloid-like assembly.

### Amyloid conversion is not preceded by a loss of condensate exchange, and low exchange is a property of the fibril itself

Of the PAM substitutions examined in Figure 2, 3R-E, K263E/3R-E and R268E/R272E produced fibrils within 24 h, and K263A/3R-A did so in a small proportion of cells (Figure 2E, 2F). To determine whether the mutants that convert differ from those that do not in the dynamics of the puncta they form beforehand, we performed FRAP on puncta at 4 h of overexpression across these constructs: the charge-reversed mutants K263E, 3R-E, K263E/3R-E and R268E/R272E, the charge-neutralized mutants K263A, 3R-A and K263A/3R-A, and the reverse-charge control 2E-K, each compared with WT and ΔPAM puncta (Figure 4A, B).

**Figure 4.**
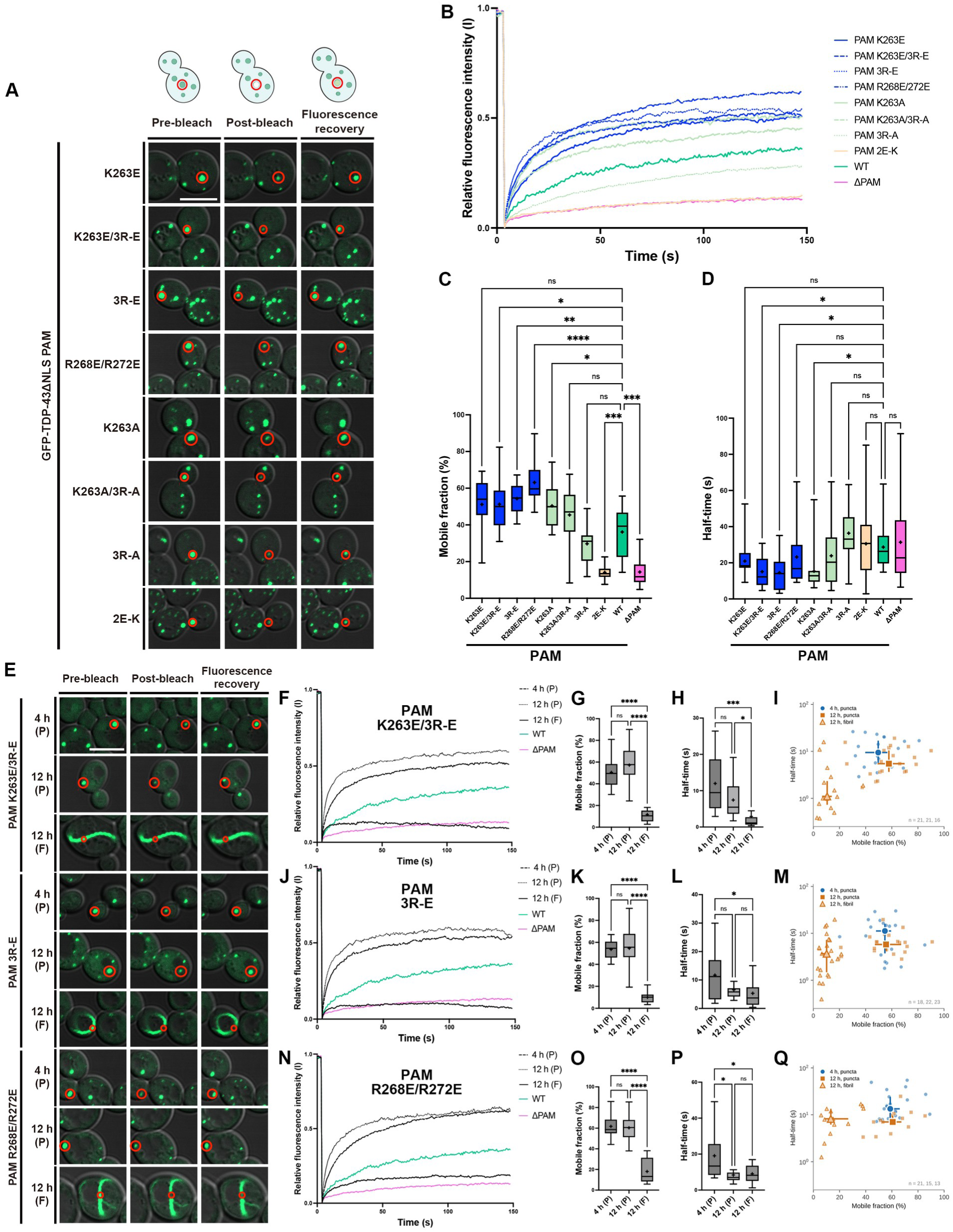
Amyloid conversion is not preceded by a loss of condensate exchange, and low exchange is a property of the fibril itself. (A) Representative confocal images of the pre-bleach, post-bleach and recovery states of puncta formed by GFP-TDP-43ΔNLS and its PAM variants (K263E, K263E/3R-E, PAM 3R-E, R268E/R272E, K263A, K263A/3R-A, PAM 3R-A and 2E-K) after 4 h of galactose induction. The bleached region of interest (ROI) is circled in red. Scale bar: 5 μm. (B) Average FRAP recovery curves of the puncta in (A): K263E, K263E/3R-E, PAM 3R-E and R268E/R272E (blue), K263A, K263A/3R-A and PAM 3R-A (green), 2E-K (orange), with WT and ΔPAM as controls. The WT and ΔPAM measurements are those shown in Figure 1E–1G. (C) Mobile fraction of the puncta in (B), taken as the fitted plateau of a one-phase association fit and expressed as a percentage of the pre-bleach intensity. Brown-Forsythe and Welch ANOVA with Dunnett’s T3, each variant versus wild type. n = 20 (K263E), 21 (K263E/3R-E), 18 (PAM 3R-E), 21 (R268E/R272E), 17 (K263A), 23 (K263A/3R-A), 20 (PAM 3R-A), 23 (2E-K), 14 (WT) and 18 (ΔPAM) structures. ns, not significant; *P < 0.05; **P < 0.01; ***P < 0.001; ****P < 0.0001. (D) Recovery half-time of the structures in (C), from the rate constant of the same fits, in seconds. Brown-Forsythe and Welch ANOVA with Dunnett’s T3, each variant versus wild type; n as in (C). ns, not significant; *P < 0.05. (E) Representative confocal images of the pre-bleach, post-bleach and recovery states of condensates of differing shape formed by GFP-TDP-43ΔNLS PAM 3R-E, K263E/3R-E and R268E/R272E after 4 and 12 h of induction. P, puncta; F, fibril; at 12 h these denote isolated structures — puncta with no attached fibril and fibrils with no attached puncta — and connected structures were excluded. The bleached ROI is circled in red. Scale bar: 5 μm. (F, J, N) Average FRAP recovery curves of condensates formed by the PAM variants under three conditions: puncta at 4 h, puncta at 12 h and fibrils at 12 h, for K263E/3R-E (F), PAM 3R-E (J) and R268E/R272E (N). Each variant is plotted with WT and ΔPAM puncta at 4 h. (G, K, O) Mobile fraction of the structures in (F, J, N), obtained as in (C), for K263E/3R-E (G), PAM 3R-E (K) and R268E/R272E (O). Brown-Forsythe and Welch ANOVA with Dunnett’s T3, comparing 4 h puncta, 12 h puncta and 12 h fibrils of the same variant; n as stated in (I), (M) and (Q). ns, not significant; ****P < 0.0001. (H, L, P) Recovery half-time of the structures in (G, K, O), obtained as in (D), for K263E/3R-E (H), PAM 3R-E (L) and R268E/R272E (P). Brown-Forsythe and Welch ANOVA with Dunnett’s T3, comparing 4 h puncta, 12 h puncta and 12 h fibrils of the same variant; n as in (G, K, O). ns, not significant; *P < 0.05. (I, M, Q) Mobile fraction against recovery half-time (logarithmic ordinate) for the individual structures in (F, J, N), for K263E/3R-E (I), PAM 3R-E (M) and R268E/R272E (Q). Small symbols, individual structures: blue circles, puncta at 4 h; filled orange squares, puncta at 12 h; open orange triangles, fibrils at 12 h. Large symbols with bars, median and interquartile range of each group. n is stated in each figure part in the order 4 h puncta, 12 h puncta, 12 h fibrils.

At 4 h no fibrils had yet formed in any construct (Figure 2E), so these measurements describe the punctate state that precedes conversion. Two constructs had the smallest mobile fractions: ΔPAM (14.3%) and the reverse-charge control 2E-K (14.1%), the only ones whose mobile fraction fell significantly below that of wild type (36.1%; Padj = 0.0004 for each; Figure 4C). By two criteria conventionally taken to indicate a matured condensate — a collapsed exchangeable pool and resistance to 1,6-hexanediol (Kroschwald et al., 2017; Alberti and Hyman, 2021) — these were the least dynamic condensates in the series, yet neither produced fibrils through 12 h (Figure 2B), and ΔPAM remained ProteoStat-negative through 24 h (Figure 2D). Every substitution that removed or reversed positive charge within PAM kept the mobile fraction above this level, from 29.7% (3R-A) to 63.1% (R268E/R272E), and all four constructs that produced fibrils within 24 h lay in this upper range (3R-E, 54.3%; K263E/3R-E, 51.2%; R268E/R272E, 63.1%; K263A/3R-A, 45.4%). Relative to wild type, the mobile fractions of R268E/R272E, 3R-E, K263E/3R-E and K263A (50.4%) were significantly higher, with K263E (51.1%) falling just short (Padj = 0.050). Half-times followed the same ordering, shortening significantly in 3R-E, K263E/3R-E and K263A (Figure 4D). The least dynamic condensates were therefore not the closest to amyloid but the furthest from it.

The fluidity of the puncta, however, did not predict how much amyloid the same mutant subsequently formed. K263E produced puncta as fluid as those of 3R-E (51.1% versus 54.3%), yet remained ProteoStat-negative throughout the 24 h time course (Figure 2D). R268E/R272E formed the most fluid puncta of any construct tested (63.1%) but converted less than either 3R-E or K263E/3R-E at every time point from 8 h onward (Figure 2E). Neither the mobile fraction (Figure 4C) nor the half-time (Figure 4D) of the puncta therefore ordered the variants as their conversion did. The mobile fractions in Figure 4C were themselves related to the net charge of PAM, across the nine constructs for which that charge is defined (R² = 0.52, P = 0.029; ΔPAM carries no PAM segment and was excluded).

We next examined the structures present once conversion has begun. In the mutants that form fibrils (K263E/3R-E, 3R-E, R268E/R272E) we compared puncta at 4 h of overexpression [4 h (P)], puncta at 12 h [12 h (P)], and fibrils at 12 h [12 h (F)] (Figure 4E, F, J, N). At 12 h both categories were scored as isolated structures: puncta with no fibril attached, and fibrils with no punctate material attached. Structures in which punctate and fibrillar material were physically connected were excluded from this comparison. For all three mutants the mobile fraction was indistinguishable between 4 h (P) and 12 h (P) and collapsed in 12 h (F) (Figure 4G, K, O). Half-times were less uniform: they were unchanged between the two punctate time points in 3R-E and K263E/3R-E but shortened in R268E/R272E (Figure 4H, L, P). Plotting mobile fraction against half-time for every individual structure resolved the two populations: puncta measured at 4 h and at 12 h occupied the same region of the plane, whereas fibrils formed a distinct population displaced along the mobile-fraction axis alone (Figure 4I, M, Q). Fibril half-times were not longer than those of puncta: shorter in K263E/3R-E, and indistinguishable from 12 h puncta in 3R-E and R268E/R272E (Figure 4H, L, P). What distinguishes a fibril is therefore the size of the exchangeable pool, not the speed at which that pool turns over.

At 12 h, therefore, two distinct states coexisted in the same population: isolated puncta whose exchangeable pool was as large as that of puncta at 4 h, and fibrils that retained almost none of theirs. These are separate structures measured at one time point, and the puncta scored at 12 h are by definition those that had not converted, so the comparison does not establish how, or whether, an individual structure passes between the two states.

### Molecular exchange declines across the puncta-to-fibril morphological series of PAM 3R-E, with the sharpest separation between the punctate and fibrillar portions of connected structures

To understand how the puncta formed by 3R-E relate to the ProteoStat-positive fibrils, we examined the structures present in cells expressing 3R-E, the variant with the strongest fibril phenotype, after 12 h of induction. Cells displayed a range of morphologies spanning puncta and fibrils, which we classified into four phenotypes: (1) puncta only, in which discrete, separate puncta are present independently within the cell; (2) connected puncta, in which individual puncta are adjoined in a chain-like arrangement but each retains its rounded, punctate shape; (3) connected fibril and puncta, in which punctate and fibrillar material coexist within one continuous structure; and (4) fibril only, consisting solely of an elongated, curved fibril with no punctate component (Figure 5A).

**Figure 5.**
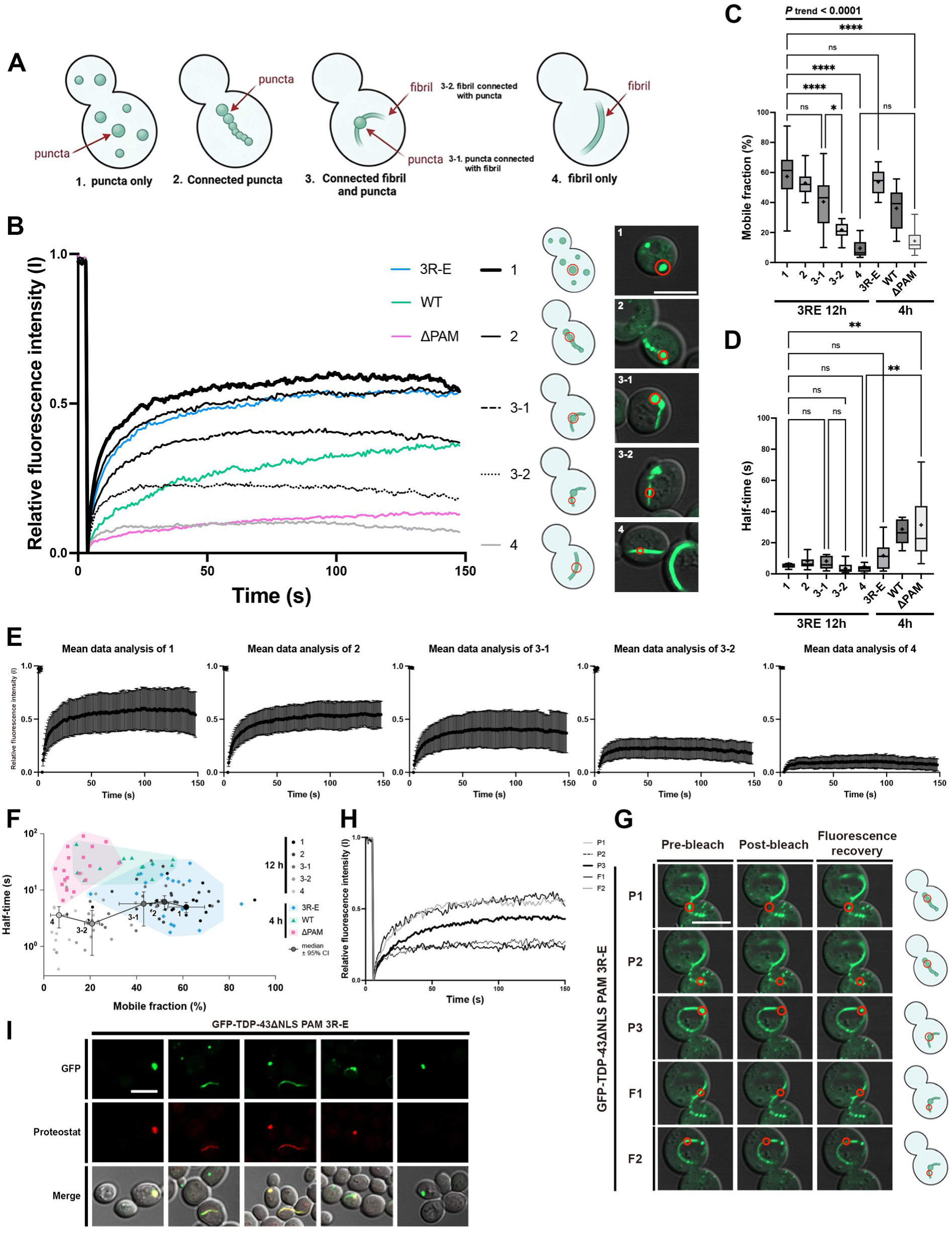
Molecular exchange declines across the puncta-to-fibril morphological series of PAM 3R-E, with the sharpest separation between the punctate and fibrillar portions of connected structures. (A) Schematic of the condensate phenotypes formed after 12 h of induction of GFP-TDP-43ΔNLS PAM 3R-E: (1) puncta only; (2) linearly connected puncta; (3) a single connected structure containing both punctate and fibrillar material, whose punctate (3-1) and fibrillar (3-2) regions were bleached and scored separately; and (4) fibril only. (B) Mean FRAP recovery curves of the phenotypes defined in (A) at 12 h, plotted with PAM 3R-E, WT and ΔPAM puncta at 4 h. The 4 h measurements, here and in (C), (D) and (F), are from the experiment shown in Figure 4B–4D. Representative merged images of a bleached structure of each phenotype, with the bleached region of interest circled in red, and the corresponding schematics are shown at right. Scale bar: 5 μm. (C) Mobile fraction of the phenotypes in (B) at 12 h, with PAM 3R-E, WT and ΔPAM puncta at 4 h. Box plots are Tukey plots. n = 15, 16, 14, 15 and 15 structures for phenotypes 1, 2, 3-1, 3-2 and 4, and 18, 14 and 18 for PAM 3R-E, WT and ΔPAM. Ptrend, Jonckheere–Terpstra test for monotone trend across the ordered phenotypes 1, 2, 3-1, 3-2 and 4; the bracket spans the categories tested. Asterisks, pairwise comparisons planned a priori. ns, not significant; *P < 0.05; ****P < 0.0001. (D) Recovery half-time of the structures in (C), in seconds. Box plots and n as in (C). Brown-Forsythe and Welch ANOVA with Dunnett’s T3, restricted to the comparisons planned a priori. ns, not significant; **P < 0.01. (E) Mean recovery curve of each phenotype at 12 h, plotted separately for phenotypes 1, 2, 3-1, 3-2 and 4. Symbols, mean; error bars, standard deviation; same structures as in (C) and (D). (F) Mobile fraction against recovery half-time (logarithmic ordinate) for the individual structures in (B), both from one-phase association fits. Grey symbols, phenotypes 1–4 defined in (A); coloured symbols, PAM 3R-E (blue), WT (green) and ΔPAM (pink) puncta at 4 h. n as in (C). Large grey symbols, phenotype medians, with error bars showing the 95% confidence interval of the median in both dimensions; the connecting line indicates phenotype order and is not a fitted trajectory. Shaded areas are convex hulls drawn as a visual guide, not confidence regions. Statistical comparisons are in (C) and (D). (G) Representative confocal images of the pre-bleach, post-bleach and recovery states of a single connected structure of a fibril with attached puncta, formed after 12 h of induction. P1–P3 denote the puncta connected to the fibril and F1–F2 positions along the fibril; each row shows the bleaching of one region, indicated in the schematic at right. Scale bar: 5 μm. (H) FRAP recovery curves of the individual regions of the structure in (G). (I) Representative confocal images of cells expressing GFP-TDP-43ΔNLS PAM 3R-E after 12 h of induction and stained with ProteoStat. Rows show the GFP channel, the ProteoStat channel and the merge with the corresponding differential interference contrast image. Each column is an independent field: a single punctum (column 1), fibrils (columns 2 and 3), a chain of linearly connected puncta (column 4) and a single punctum (column 5). Scale bar: 5 μm.

FRAP of each phenotype showed that the mobile fraction fell across this morphological series (Figure 5B, 5C). The mean recovery curve of each phenotype is also plotted separately in Figure 5E, and the individual curves that contribute to each mean in Figure S2. The two parts of phenotype 3 could be bleached independently, so the punctate region (3-1) and the fibrillar region (3-2) were recorded as separate measurements. Both regions were bleached in some structures but only one of the two in others, so 3-1 and 3-2 were analyzed as independent samples and not as paired measurements. The mean mobile fraction was 57.4 ± 18.6% for phenotype 1, 53.0 ± 11.7% for phenotype 2, 40.5 ± 16.9% for 3-1, 21.8 ± 9.1% for 3-2 and 9.5 ± 5.7% for phenotype 4 (mean ± SD; n = 15, 16, 14, 15 and 15) (Figure 5C). The same plots include WT, ΔPAM and 3R-E puncta measured at 4 h of induction, from the same experiment as Figure 4C and shown here for reference: phenotype 1 lay at the level of 3R-E puncta before conversion (53.7%), whereas phenotype 4 fell well below wild type (36.1%). Because these phenotypes form an ordered series, we tested the decline directly with the Jonckheere–Terpstra test for monotone trend, which confirmed the decline (*P*_trend_ < 0.0001). The decline was not evenly distributed along the series: successive phenotypes differed by 4.4, 12.5, 18.7 and 12.3 percentage points, the largest single step falling between the punctate and the fibrillar region of connected structures (Figure 5C). Pairwise comparisons were consistent with this distribution. Phenotype 1 did not differ from 3-1, whereas 3-1 differed from 3-2, and 3-2 and phenotype 4 both differed from phenotype 1 (Figure 5C). Punctate material thus retained most of its exchange even when it was part of a structure that already contained a fibril, while the fibrillar part of the same class of structure did not.

The spread of values narrowed along the same series. Mobile fractions ranged from 19.3% to 90.9% among phenotype 1 puncta but from 3.4% to 21.3% among phenotype 4 fibrils, and the standard deviation fell from 18.6% to 5.7% (Figure 5C). Free puncta were therefore heterogeneous in their exchange, whereas fibrils were uniformly low.

Recovery half-times did not follow the mobile fraction. Across the series the half-time remained short (6.4, 8.2, 8.1, 3.6 and 4.9 s for phenotypes 1, 2, 3-1, 3-2 and 4) and phenotype 4 did not differ from phenotype 1 (Figure 5D). ΔPAM puncta at 4 h reached a comparably small mobile fraction (14.3 ± 7.2%, not different from phenotype 4) but recovered with a half-time of 31.4 s, significantly longer than that of phenotype 4. The fibrils of 3R-E and the condensates of ΔPAM therefore arrive at a similarly small exchangeable pool in different ways: in the fibril the remaining mobile pool still exchanges rapidly, whereas in ΔPAM condensates exchange itself is slow. Plotting the two parameters against each other for individual structures separated them accordingly, with the 3R-E phenotypes distributed along the mobile-fraction axis and ΔPAM puncta displaced along the half-time axis (Figure 5F).

To ask whether the difference between punctate and fibrillar material is local rather than a difference between cells, we bleached punctate regions (P1–P3) and fibrillar regions (F1–F2) separately within one continuous structure. The punctate regions recovered to a higher level than the fibrillar regions of the same structure (Figure 5G, 5H), showing that the two kinds of material can differ in exchange while physically connected. This measurement was made on a single structure; the quantitative comparison is the population analysis in Figure 5C.

ProteoStat staining in these cells showed that amyloid-like character was likewise not uniform across these structures. Fibrils were ProteoStat-positive along their length, whereas puncta were heterogeneous, with ProteoStat-positive and ProteoStat-negative puncta present in the same cell population. The same heterogeneity was present within a single structure: in a chain of connected puncta, only the punctum at one end was labeled while the remaining puncta of that chain were not (Figure 5I). ProteoStat reactivity therefore differs between individual puncta, including between puncta of one connected structure.

Taken together, molecular exchange declines monotonically across the puncta-to-fibril series, but not uniformly along it. Punctate material retains most of its exchange even when attached to a fibril, and the steepest fall occurs between the punctate and fibrillar portions of connected structures.

### PAM charge reversal drives amyloid-like conversion of TDP-43 condensates in human U2OS cells, independently of condensate exchange

To ask whether the behavior established in yeast holds in human cells, we expressed the same NLS-deficient GFP-TDP-43 constructs in U2OS cells from a CMV promoter and examined them 48 h after transfection, without applying any stress. In mammalian cells cytoplasmic TDP-43 is ordinarily driven into visible assemblies by an acute insult: oxidative stress such as sodium arsenite recruits it into stress granules (Colombrita et al., 2009; Liu-Yesucevitz et al., 2010), although cytoplasmic TDP-43 can also de-mix independently of them once it accumulates outside the nucleus (Gasset-Rosa et al., 2019). We deliberately omitted such a trigger, because the question here is what the charged clusters contribute to assembly on their own, not how TDP-43 partitions into a stress-induced compartment. Whatever formed under these conditions therefore reflects what the sequence does in the absence of an acute trigger.

Under these conditions wild-type GFP-TDP-43ΔNLS did not form the large cytoplasmic puncta it forms in yeast. Instead the protein was distributed as fine cytoplasmic puncta, present in about 69% of cells (Figure 6A; Figure S3A, S3B). Deleting either polyanionic cluster removed even these: ΔPA1 and ΔPA2 cells were diffuse, with puncta in fewer than 20% of cells, as were ΔPAMΔLCD cells (Figure 6A; Figure S3A, S3B). The charge-neutralization variants behaved the same way, with one exception: PA1-1 retained puncta in 65% of cells, as it had in yeast, whereas PA1, PA1-2, PA2, PA2-1 and PA2-2 all fell below 20% (Figure S3A, S3B).

**Figure 6.**
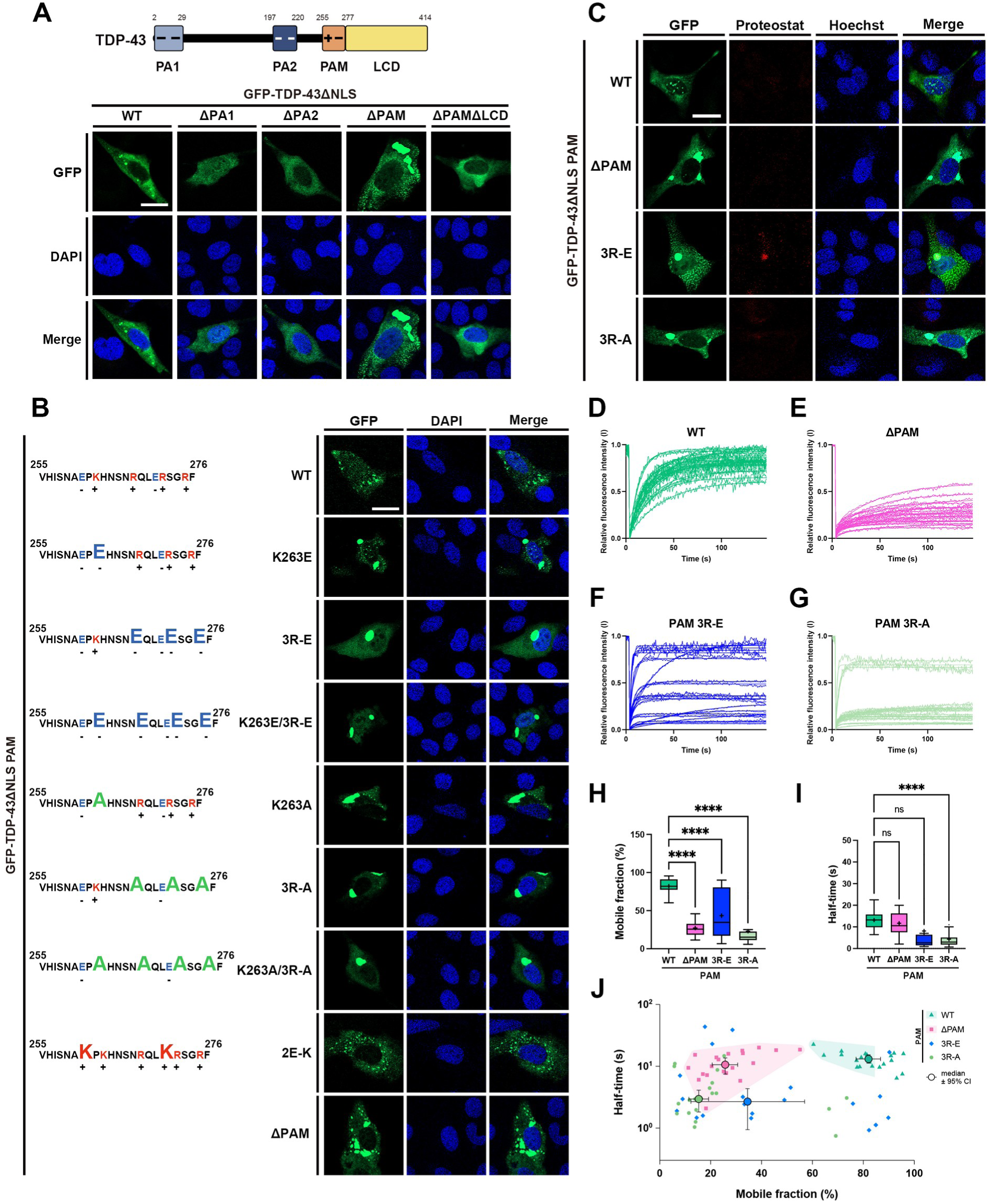
PAM charge reversal drives amyloid-like conversion of TDP-43 condensates in human U2OS cells, independently of condensate exchange. (A) Schematic of the charged cluster architecture in TDP-43 (top) and representative confocal images of U2OS cells expressing NLS-deficient GFP-tagged TDP-43 WT and its variants (ΔPA1, ΔPA2, ΔPAM and ΔPAMΔLCD) 48 h after transfection (bottom). GFP, DAPI and merged images are shown. Scale bar: 20 μm. (B) Representative confocal images of U2OS cells expressing NLS-deficient GFP-tagged TDP-43 WT and its PAM mutants (K263E, PAM 3R-E, K263E/3R-E, K263A, PAM 3R-A, K263A/3R-A, 2E-K and ΔPAM) 48 h after transfection (right), with the corresponding PAM sequences at left and substituted residues in capital letters. Cells were counterstained with DAPI. Scale bar: 20 μm. (C) Representative confocal images of U2OS cells expressing NLS-deficient GFP-tagged TDP-43 WT, ΔPAM, PAM 3R-E and PAM 3R-A 48 h after transfection, fixed and stained with ProteoStat and Hoechst. Scale bar: 20 μm. (D–G) Individual FRAP recovery curves of cytoplasmic condensates in U2OS cells expressing NLS-deficient GFP-tagged TDP-43 WT (D), ΔPAM (E), PAM 3R-E (F) and PAM 3R-A (G), 48 h after transfection. Every curve recorded for that construct is plotted (n = 20 condensates each). (H) Mobile fraction of the condensates in (D–G), taken as the fitted plateau of a one-phase association fit and expressed as a percentage of the pre-bleach intensity. Box plots are Tukey plots. n = 20 condensates per construct. Brown-Forsythe and Welch ANOVA with Dunnett’s T3, each construct versus wild type. ****P < 0.0001. (I) Recovery half-time of the condensates in (H), from the rate constant of the same fits, in seconds. Box plots and n as in (H). Brown-Forsythe and Welch ANOVA with Dunnett’s T3, each construct versus wild type. ns, not significant; ****P < 0.0001. (J) Mobile fraction against recovery half-time (logarithmic ordinate) for the individual condensates in (H) and (I). Small symbols, individual condensates: WT (green), ΔPAM (pink), PAM 3R-E (blue) and PAM 3R-A (light green). Large symbols, construct medians, with error bars showing the 95% confidence interval of the median in both dimensions. Shaded areas are convex hulls drawn as a visual guide, not confidence regions.

The two systems differ here. In yeast, wild type formed large mature puncta and deleting either cluster left puncta that were smaller and more numerous (Figure 1B); in U2OS cells, wild type formed only small puncta and the same deletions, together with most of the neutralization variants, removed them altogether. The two systems therefore agree that PA1 and PA2 are required for normal condensate formation, but differ in what their loss produces: immature puncta in yeast, and no puncta in U2OS cells.

Against this background ΔPAM was the exception. It formed a small number of cytoplasmic puncta larger and brighter than those of wild type, the only deletion in either system to produce assemblies at least as large as those of wild type (Figure 6A). The PAM point mutants behaved similarly: cells expressing K263E, 3R-E, K263E/3R-E, K263A, 3R-A or K263A/3R-A contained large puncta, whereas the reverse-charge control 2E-K resembled wild type (Figure 6B). The elongated, curved structures seen in yeast were not observed in U2OS cells within the 48 h examined here; all of the material was punctate.

We then stained wild type, ΔPAM, 3R-E and 3R-A with ProteoStat. Of these four, only 3R-E was positive; wild type, ΔPAM and 3R-A were all negative, ΔPAM despite forming the largest puncta of the four (Figure 6C; Figure S3C–S3F). The signal was not uniform across the 3R-E population. Within one field, condensates in some 3R-E cells were ProteoStat-positive while those in neighboring cells were not (Figure S4A, S4B), so a culture expressing 3R-E contains labeled and unlabeled puncta together rather than a uniformly converted population.

FRAP of individual cytoplasmic condensates 48 h after transfection separated the constructs by mobile fraction (Figure 6D–6G). Wild-type condensates were highly dynamic, recovering to 82.0 ± 9.9% of their pre-bleach intensity — far above the 36.1% measured for wild-type puncta in yeast, consistent with the small, labile assemblies formed here in the absence of stress. All three variants recovered less: 27.0 ± 11.1% for ΔPAM, 43.5 ± 30.4% for 3R-E and 23.0 ± 20.9% for 3R-A (mean ± SD; n = 20 condensates each; P < 0.0001 for each comparison with wild type; Figure 6H). Half-times separated them differently: ΔPAM (11.7 ± 5.1 s) did not differ from wild type (13.1 ± 3.8 s) and 3R-A recovered faster (4.3 ± 3.8 s, P < 0.0001; Figure 6I). The half-times of 3R-E did not differ from those of wild type by this test (8.1 ± 12.6 s), but their distribution was strongly right-skewed — a median of 2.7 s with a long tail to 43.4 s — so the mean does not describe the group well.

The 3R-E recovery curves were heterogeneous: some condensates recovered almost completely within seconds while others barely recovered over the same interval (Figure 6F; Figure S4C, S4D), and the dispersion lay between cells. Individual 3R-E condensates spanned mobile fractions from 6.7% to 90.1%, an interquartile range of 60.0 percentage points against 13.4 for wild type, 14.1 for ΔPAM and 10.8 for 3R-A (Figure 6H). The larger standard deviation of 3R-A (20.9) reflects a few high values rather than a broad distribution: its interquartile range was the narrowest of the four. Plotting mobile fraction against half-time made this explicit: wild type and ΔPAM each occupied a compact region, wild -type at high mobile fraction and ΔPAM at low, whereas 3R-E spread across the whole range, overlapping both and extending below them in half-time (Figure 6J). A 3R-E population at one time point is therefore not a set of equivalent condensates but one distributed between the dynamic and the immobile, mirroring the heterogeneity of exchange seen among 3R-E structures in yeast (Figure 5C).

Two properties of 3R-E were thus reproduced in human cells: its condensates acquired amyloid-like character, and they did so unevenly, differing from cell to cell in both ProteoStat reactivity and molecular exchange. Neither followed from a loss of exchange. ΔPAM formed the largest condensates of the four and 3R-A the smallest exchangeable pool, and both exchanged less than 3R-E, yet neither was ProteoStat-positive (Figure 6C, 6H). As in yeast, charge reversal within PAM rather than the material state of the condensate determined whether TDP-43 acquired amyloid-like character.

### The net charge of PAM, rather than the identity of the substituted residue, sets the rate of amyloid conversion

The full charge series could be examined only in yeast, the system that provides the readout: scoring conversion requires well-defined puncta that become unambiguous fibrils, and U2OS cells provide neither (Figure 6A, 6B). The substitutions examined so far altered PAM in several distinct ways — removing positive charge (3R-A, K263A), reversing it (3R-E, K263E), or adding positive charge (2E-K). We therefore asked whether they act through a single underlying variable: the net charge of the PAM segment itself.

To vary PAM charge independently of its basic residues, we introduced acidic substitutions at the two serines within the cluster, S266 and S273, alone and in combination (S266D, S266E, S273D, S273E, S266D/S273D, S266E/S273E), together with the charge-neutral control S266A/S273A. These substitutions place a single negative charge at each position, matching the sign if not the magnitude of the charge introduced by phosphorylation. We also combined these with 3R-E (3R-E_S266D/S273D, 3R-E_S266E/S273E) and with the corresponding alanine control (3R-E_S266A/S273A) (Figure 7A, 7B). Amyloid-like fibril formation was scored from ProteoStat-positive filamentous structures after 12, 24, 48 and 72 h of galactose induction across all 24 constructs (Figure 7C). This time course was performed as a single independent experiment covering every construct, so its absolute percentages differ from those of the shorter time course in Figure 2E — 3R-E reached 11.3% of cells at 24 h here against 18.2% there — while the ranking of the constructs is preserved. Comparisons are therefore made within each experiment.

**Figure 7.**
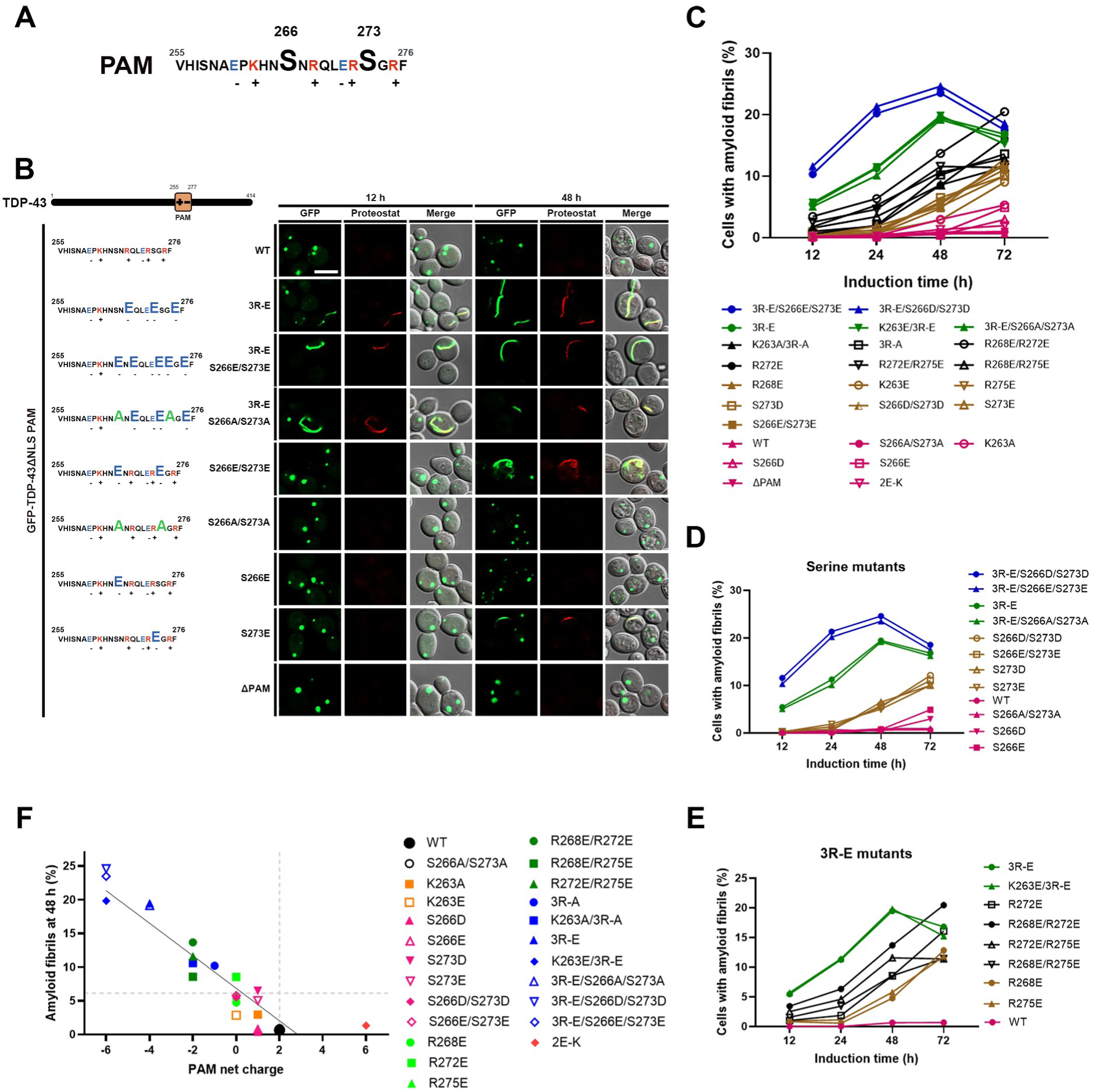
The net charge of PAM, rather than the identity of the substituted residue, sets the rate of amyloid conversion. (A) Amino acid sequence of the PAM cluster (residues 255–276) of TDP-43. Basic residues are shown in red and acidic residues in blue, with the sign of each charge indicated below; the two serines substituted in this figure, S266 and S273, are enlarged. (B) Schematic of TDP-43 showing the position of PAM (left, top) and the PAM sequences of the constructs used (left, bottom), with representative confocal images of W303a yeast cells expressing each GFP-TDP-43ΔNLS variant after 12 and 48 h of galactose induction (right). Cells were fixed and stained with ProteoStat; GFP, ProteoStat and merged images are shown for each time point. Scale bar: 5 μm. (C) Percentage of cells containing amyloid fibrils, scored from ProteoStat-positive filamentous structures, for all PAM variants after 12, 24, 48 and 72 h of galactose induction. More than 300 cells were scored per construct and time point; each point is a single pooled proportion, and no replicate-based statistics were applied. This time course is an independent experiment from the shorter one in Figure 2E and 2F, with which it overlaps at 12 and 24 h; absolute percentages therefore differ, and comparisons are made within each experiment. (D) The serine substitution series from (C) shown separately: the phosphomimetic single and double mutants (S266D, S266E, S273D, S273E, S266D/S273D, S266E/S273E), the charge-neutral control S266A/S273A, and their combinations with 3R-E (3R-E_S266D/S273D, 3R-E_S266E/S273E, 3R-E_S266A/S273A), with WT and PAM 3R-E for comparison. (E) The arginine substitution series from (C) shown separately: the single (R268E, R272E, R275E) and paired (R268E/R272E, R268E/R275E, R272E/R275E) substitutions, together with PAM 3R-E and K263E/3R-E, with WT for comparison. (F) Percentage of cells containing amyloid fibrils at 48 h plotted against the calculated net charge of PAM for all 24 constructs. Each symbol is one construct; the solid line is the least-squares linear fit (Y = −2.413 X + 6.852; slope −2.413 ± 0.217, 95% CI −2.864 to −1.963; R² = 0.85; F(1, 22) = 123.5, P = 1.7 × 10⁻¹⁰). The vertical dashed line marks the net charge of wild-type PAM (+2); the horizontal dashed line marks the median percentage of fibril-bearing cells across the 24 constructs (6.1%). ΔPAM, which has no PAM segment and therefore no defined net charge, is not included in this plot.

Adding negative charge at the serines conferred fibril-forming capacity on its own (Figure 7B, 7D). S266E/S273E and S266D/S273D, which leave all four basic residues intact, produced fibrils in 5.8% and 5.5% of cells at 48 h and in 11.1% and 12.1% by 72 h, against 0.9% for the charge-neutral control S266A/S273A and 0.6% for wild type, neither of which rose over the same period (Figure 7D). The single substitutions behaved in the same way but more slowly: S273D and S273E reached 10.0% by 72 h, and S266D and S266E, which were still at or below the wild-type level at 48 h, rose to 2.9% and 4.9% (Figure 7D). Conversely, combining the serine substitutions with 3R-E accelerated fibril formation from the earliest time point: 3R-E_S266D/S273D and 3R-E_S266E/S273E had already reached 11.6% and 10.3% at 12 h, against 5.4% for 3R-E, and peaked at 24.6% and 23.5% at 48 h, the highest of any construct tested, whereas the alanine counterpart 3R-E_S266A/S273A tracked 3R-E throughout (19.2% versus 19.5% at 48 h) (Figure 7D).

The arginine substitutions formed a dose series (Figure 7E). Reversing a single arginine lowered the net charge of PAM from +2 to 0 and produced fibrils in 4.8%, 8.5% and 5.8% of cells at 48 h for R268E, R272E and R275E. Reversing two lowered it to −2 and raised these values to 13.7% for R268E/R272E, 11.6% for R272E/R275E and 8.6% for R268E/R275E. Reversing all three (3R-E, net charge −4) raised them further to 19.5%, and adding K263E (net charge −6) to 19.8%. No single arginine accounted for the effect: each of the three contributed on its own, and every pair and the triple exceeded any single substitution.

Plotting the proportion of cells with amyloid fibrils at 48 h against the calculated net charge of PAM for all 24 constructs revealed a single linear relationship across the series (Figure 7F): Y = −2.413 X + 6.852, with a slope of −2.413 ± 0.217 (95% CI −2.864 to −1.963), R² = 0.85 and F(1, 22) = 123.5, P = 1.7 × 10⁻¹⁰.

Substitutions of lysine, arginine and serine fell on the same line, as did their combinations, indicating that the identity of the substituted residue is not what matters: each unit of negative charge added to PAM raised the proportion of cells bearing fibrils by approximately 2.4 percentage points.

One construct departed from this relationship. 2E-K, in which the two acidic residues of PAM are replaced by lysine to give the most positive net charge of the series (+6), still produced 1.3% fibril-bearing cells rather than none, and therefore lies above the fitted line, which crosses zero at a net charge of +2.8. Net charge thus accounts for the behavior of the series across the acidic range, while strongly basic PAM variants are not described by a simple linear extrapolation. Excluding 2E-K as a sensitivity analysis yields a steeper and tighter fit (slope −2.867, R² = 0.93, crossing zero at a net charge of +2.1); the 24-construct fit is the one reported throughout.

The strength of this relationship was itself time-dependent. Fitting the same regression at each time point returned slopes of −0.93, −1.85, −2.41 and −1.57 percentage points per charge unit at 12, 24, 48 and 72 h, with R² = 0.70, 0.75, 0.85 and 0.63 respectively. The relationship therefore tightened up to 48 h and then loosened, because the series converged: constructs with little negative charge continued to accumulate fibrils between 48 and 72 h — S266D/S273D rose from 5.5% to 12.1% and 3R-A from 10.2% to 13.6% — whereas the fastest converters had already reached their maximum by 48 h. Net charge thus sets the rate at which conversion proceeds rather than a fixed endpoint, and 48 h is the time at which the variants are maximally separated; this is the time point plotted in Figure 7F.

The ordering set by charge was not the ordering set by condensate dynamics. 3R-A formed puncta indistinguishable from those of wild type in both mobile fraction and half-time (Figure 4C, 4D) yet converted in 10.2% of cells at 48 h, whereas K263A, whose puncta were as mobile as those of 3R-E (50.4% versus 54.3%; Figure 4C), converted in only 3.0% (Figure 7C). Conversion followed the net charge of PAM rather than the material state of the condensate each variant formed. Because the mobile fractions of those puncta were themselves related to PAM net charge (Figure 4C), the two appear to be set by charge in parallel rather than the one acting through the other.

## DISCUSSION

Three findings define the contribution of this work. First, the net charge of a short polyampholyte motif outside the TDP-43 amyloid core determines whether condensates convert to amyloid in cells, scaling linearly across 24 variants and independently of the substituted residue. Second, this determinant operates independently of condensate material state: in yeast the variants that converted were among the most mobile of those tested, and in both systems the condensates with the smallest exchangeable pools remained ProteoStat-negative throughout. Third, phosphomimetic substitution of the motif’s serines was sufficient to trigger conversion, identifying a route by which this determinant could be set physiologically. Together these results place a defined sequence element upstream of amyloid conversion and separate that conversion from the solidification with which it has long been equated.

Our results run counter to that aging model (reviewed in Ahmad et al., 2022; Visser et al., 2024) at several points. ΔPAM, in which the motif is deleted entirely, formed among the least dynamic puncta of any construct tested, yet did not convert within the times examined (Figure 1C; Figure 2D), and the reverse-charge control 2E-K, equally static (Figure 4C), produced only 1.3% fibril-bearing cells against 0.6% for wild type (Figure 7F). The mutants that did convert formed puncta markedly more fluid than those of wild type beforehand (Figure 4), and that state did not become progressively less dynamic with time: for a given mutant, puncta at 12 h were indistinguishable from puncta at 4 h (Figure 4G, 4K, 4O). Loss of exchange was instead confined to individual structures as they converted. ProteoStat-positive and ProteoStat-negative puncta coexisted within one cell, including within a single chain of connected puncta, and exchange differed between regions of one structure (Figure 5G, 5H, 5I); the punctate portion of a structure that had already formed a fibril exchanged no less than free puncta, and the fall occurred at the fibrillar portion itself (Figure 5C). Loss of molecular exchange is therefore a local accompaniment of amyloid conversion rather than a uniform, time-dependent process that precedes and drives it.

What, then, determines whether conversion occurs? This may depend on where PAM lies relative to the amyloid fold. In cryo-EM structures of TDP-43 filaments extracted from patient brain, the ordered core comprises residues 282–360 in ALS with FTLD and residues 272–360 in type A FTLD-TDP, while the flanking sequences form an unstructured fuzzy coat that is not resolved (Arseni et al., 2022, 2023). The PAM (residues 255–276) lies almost entirely within that unresolved region: K263 and R268 fall outside the core of both folds, and R272 and R275 reach only its N-terminal edge in the type A fold. The charge that sets the rate of conversion therefore resides in a segment that adopts no defined structure in the mature filament, which places PAM upstream of the ordered core rather than within it: the electrostatics of a disordered flanking segment tune how readily the adjacent sequence enters the amyloid fold. This would explain why the identity of the substituted residue is immaterial while its charge is not. Nor is such charge dependence specific to PAM: the requirements of PA1 and PA2 also map to internal acidic patches rather than to the cluster as a whole (Figure S1). The same segment is poorly ordered in patient filaments, leaving exposed the W334–A341 hydrophobic surface proposed to mediate aberrant interactions (Arseni et al., 2023; Ulamec et al., 2020). On that reasoning, deleting the segment should favor conversion, which is the opposite of what we observe for ΔPAM; exposure of this surface is therefore not by itself sufficient to drive conversion in cells.

The same position that places PAM outside the amyloid fold places it immediately next to RRM2, which raises an alternative reading of the substitutions: that they act by weakening RNA binding rather than by changing charge. Three observations argue against this. Deleting the segment entirely (ΔPAM) abolishes conversion rather than enhancing it, the opposite of what loss of RNA binding would predict; the S→D and S→E substitutions add negative charge without removing any basic residue, yet confer fibril-forming capacity; and 3R-A and 3R-E remove the same arginines but convert at different rates, so the substituting residue rather than the loss of arginine determines the outcome. These observations do not exclude a contribution from RNA binding altogether — increasing the negative charge of a segment adjacent to RRM2 could itself weaken binding, and the two explanations need not be mutually exclusive. We therefore interpret net charge as the parameter that tracks conversion, without assigning it a single mechanism.

The same charge-reversal specificity was reproduced in an entirely different cellular environment. In U2OS cells, wild-type and charge-neutralized mutants remained ProteoStat-negative, whereas only the charge-reversed mutant produced ProteoStat-positive condensates (Figure 6). This indicates that the phenomenon does not depend on the chaperone complement or metabolic state of yeast — a relevant distinction given that HSPB1 modulates cytoplasmic TDP-43 phase separation and its liquid-to-gel transition (Lu et al., 2022). The comparison has a limit in one direction: U2OS cells express the construct in a minority of the population and cannot be followed through successive rounds of cell division, so the variants that in yeast convert only by 48 to 72 h could not be tested there, and their absence in U2OS is not evidence that they do not convert.

Two qualifications bound this account. First, the interface-nucleation work set out above used reconstituted single-component condensates or isolated low-complexity domains (Das et al., 2025; Linsenmeier et al., 2023; Shen et al., 2023); what set the rate of conversion here was not the geometry of the condensate but the net charge encoded in PAM, a determinant of a different kind rather than a contradiction of those models. Second, this is one route among several: cytoplasmic TDP-43 aggregation also proceeds independently of phase separation (Watanabe et al., 2020), and phase separation promotes spontaneous but delays seeded aggregation of the low-complexity domain (Pakravan et al., 2021), so condensate formation need not act in a single direction; and in human cells under oxidative stress, TDP-43 demixes within stress granules into a phase that later becomes ubiquitin- and phospho-positive aggregates (Yan et al., 2025).

Taken together, our observations suggest the following account of how a TDP-43 condensate becomes an amyloid-like fibril (Figure 8). Lowering the net charge of PAM increases the exchange of TDP-43 between the condensate and the surrounding cytoplasm and, separately, raises the rate at which that condensate converts. Conversion then proceeds structure by structure rather than across the population: within a single cell some condensates acquire amyloid while neighboring ones do not, and exchange is lost only in those that are converting. The same heterogeneity was seen in human cells, where only some 3R-E condensates were ProteoStat-positive and individual condensates differed widely in exchange (Figure 6). Because TDP-43 is hyperphosphorylated in disease and phosphorylation alters its polymerization and phase behavior (Gruijs da Silva et al., 2022; Kellett et al., 2025; Wang et al., 2018a), phosphorylation of S266 and S273 would lower PAM net charge in the same direction as the phosphomimetic substitutions did (Figure 7), offering a potential route by which conversion could be driven in disease without any change in sequence.

**Figure 8.**
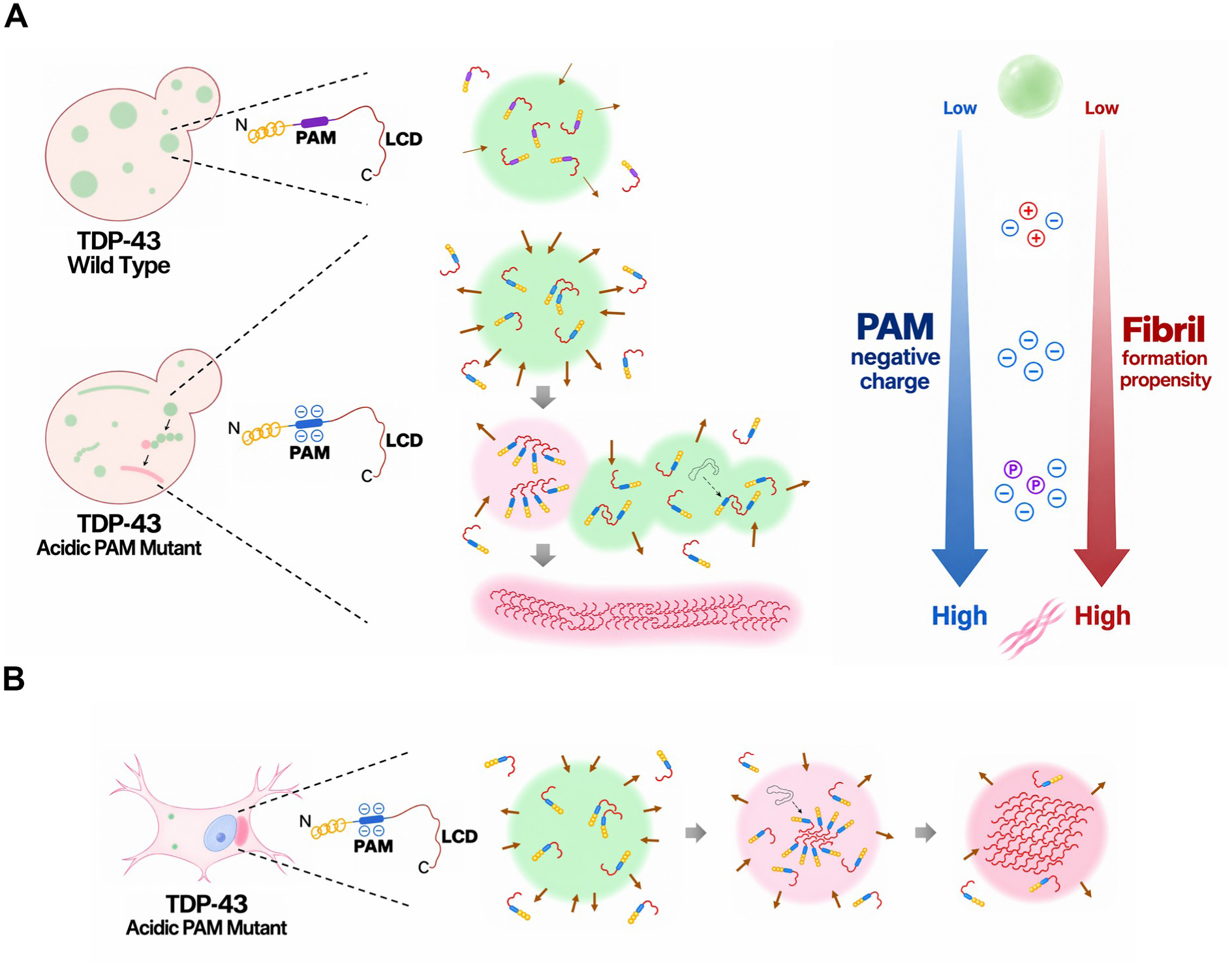
A model: PAM net charge as a tunable determinant for TDP-43 amyloid conversion. (A) In budding yeast, wild-type TDP-43 forms condensates that exchange little with the surrounding cytoplasm (top). Lowering the net charge of PAM increases that exchange (middle) and raises the rate at which individual condensates convert: conversion proceeds structure by structure, converting condensates (pink) losing exchange while neighboring condensates (green) remain dynamic, and ends in a fibril (bottom). The gradient on the right summarizes the relationship established in Figure 7: the more negative the net charge of PAM — whether reached by substitution or, potentially, by phosphorylation of S266 and S273 — the greater the propensity to form fibrils. (B) The same sequence of events in a mammalian cell: a dynamic condensate formed by an acidic PAM variant loses exchange as amyloid accumulates within it and converts in place.

An important next step will be to determine whether the fibrils formed by 3R-E adopt the fold seen in patient-derived TDP-43 fibrils, which will require solving their mutant structure by cryo-EM. This comparison is not straightforward: fibrils assembled in vitro from the TDP-43 low-complexity domain adopt folds distinct from the patient ones and are themselves polymorphic (Li et al., 2021; Shenoy et al., 2023). If the folds prove identical, the charge state of the disordered PAM segment would be shown to govern formation of the same amyloid core seen in patient pathology. A second direction is whether this charge state can be shifted without mutation — in particular, whether phosphorylation at S266 and S273 occurs in disease.

## METHODS

### Domain prediction and charge-cluster analysis

Intrinsically disordered regions (IDRs) of TDP-43 were predicted with four independent algorithms: IUPred2A, PrDOS, DISOPRED3 and PASTA 2.0. Aggregation-prone regions were predicted with AGGRESCAN and PASTA 2.0. Within the predicted IDRs, the distribution of charged residues was then analyzed using a sliding-window calculation of net charge along the sequence, which was used to designate the PA1, PA2 and PAM clusters.

### PCR-mediated cloning of plasmid DNA

The ΔNLS (K82A, R83A, K84A) mutant of TDP-43, in which three residues of the nuclear localization signal (NLS) were substituted with alanine residues to disrupt NLS function, was constructed by PCR-mediated cloning using primers flanking the substitution sites.

PCR-mediated cloning was also employed to generate DNA constructs of the ΔPA1, ΔPA2, ΔPAM, ΔPAMΔLCD and ΔPA1ΔPA2 mutants of TDP-43ΔNLS. For each deletion, primers complementary to 30 bp flanking the target domain were designed, with additional overlapping sequences incorporated at the 3′ ends of the primers to enable amplification. Following PCR, the resulting products were treated with DpnI for 1 h to eliminate the parental plasmid template. The digested products were subsequently transformed into E. coli, where homologous recombination facilitated the reconstitution of circular plasmids. Plasmid DNA was then extracted and verified by Sanger sequencing.

TDP-43ΔNLS PA1, PA2, and PAM residue substituted variants were also constructed by PCR-based site-directed mutagenesis using pMW20(U)-pGAL-GFP-TDP-43ΔNLS and pcDNA3.1-myc-HisA-GFP-TDP-43ΔNLS as templates. Primer sequences are provided in the Key Resources Table, and all mutant constructs were confirmed by DNA sequencing.

### Yeast cell culture, transformation, and protein induction

*Saccharomyces cerevisiae* yeast cells were cultured as mainly described in Choi and Song (2022). Yeast cells grown in YPAD medium until reaching the exponential growth phase (OD_600_ 0.5) were transformed with 100 ng plasmid DNA by heat-shocked at 42 °C for 30 min in the yeast transformation mixture consisting of 50% polyethylene glycol (Sigma-Aldrich, USA, P4338-500G), 1 M LiOAc (Sigma-Aldrich, USA, L6883-250G), 2 mg/mL salmon sperm DNA (Sigma, USA, D1626), and ddH_2_O, as described in Choi and Song (2022).

For protein overexpression, yeast cells carrying plasmids encoding TDP-43 and their respective variants were initially grown in SC-Ura medium supplemented with 2% glucose until reaching the exponential growth phase, and further cultured for 3 h in SC-Ura medium containing 2% raffinose and 0.1% glucose after 3 washes. Induction of gene expression was subsequently initiated by supplementing the culture with galactose to a final concentration of 2%, followed by incubation for the indicated time periods.

### 1,6-Hexanediol treatment of yeast cells

Yeast cells induced for protein overexpression in 2% galactose for indicated time periods were harvested by centrifugation at 4000 rpm for 5 minutes, and the resulting pellet was collected. The cell pellet was then resuspended and incubated in SC-Ura medium supplemented with 2% galactose, 2% raffinose, 0.1% glucose, and 10% 1,6-hexanediol (Daejung Chemical & Metals Co., LTD, South Korea, 629-11-8) for 1 h.

### Confocal microscopy of cells and image analysis

Except where stated otherwise, GFP-labeled proteins expressed in the cells were imaged using both a ZEISS LSM900 laser scanning confocal microscope (Carl Zeiss, Jena, Germany) equipped with high-sensitivity GaAsP-PMT detectors (2 standard and 1 additional GaAsP-PMT), and a ZEISS LSM980 confocal microscope with an Airyscan 2 detector (Carl Zeiss, Jena, Germany). For Airyscan super-resolution imaging with the LSM980, the Airyscan Joint Deconvolution mode was utilized, achieving lateral resolution down to ∼90 nm. Yeast cells were imaged with a 100×/1.40 NA oil-immersion objective and U2OS cells with a 20× objective, and all images were acquired at room temperature. Microscope settings, including laser power, detector gain, and pinhole size, were kept identical for all samples to ensure quantitative comparability. Image acquisition and processing were performed using ZEISS Zen software (version 3.9, Carl Zeiss).

For yeast cells, images were acquired on the LSM900 using the 488 nm laser at 0.6% power and a GaAsP-PMT detector. Because of the low GFP signal and the small size of yeast cells, the pinhole was opened to 3 Airy units (approximately 240 μm). Images were acquired at a frame size of 1024 × 1024 pixels, 8 bits per pixel, scan speed 9 (approximately 1 μs pixel dwell time), unidirectional scanning and 8- or 16-fold averaging. Identical culture conditions, exposure settings and acquisition parameters were used for all strains so that puncta intensities could be compared directly. Images were processed and analyzed with ZEN lite and Adobe Photoshop.

For the three-dimensional reconstruction shown in Figure 3A, z-stacks were acquired on a Leica STELLARIS 5 confocal microscope (Leica Microsystems, Wetzlar, Germany) fitted with a 63×/1.40 oil-immersion objective, with GFP excited at 488 nm and a z-step of 0.148 μm over 34–58 optical sections. Stacks were deconvolved with the LIGHTNING adaptive deconvolution module and rendered as three-dimensional volumes in Leica LAS X (version 4.9.2).

### FRAP in living yeast cells

FRAP was performed on puncta induced from the GAL promoter in yeast cells with the same condition for fluorescence imaging. Images were acquired at a frame size of 512 × 512 pixels, 8 bits per pixel, scan speed 9 (approximately 1 μs pixel dwell time), unidirectional scanning and without averaging. Each time series comprised 200 cycles acquired continuously, with no added delay between cycles, giving a frame interval of 0.738 s and a total observation window of 146.8 s. The entire punctum was selected as the bleach region of interest (ROI), so that recovery reports exchange between the structure and the surrounding cytoplasm; ROI size therefore varied with the size of each structure. Five pre-bleach images were acquired, after which the ROI was bleached with three consecutive iterations of the 488 nm laser at 5–15% power, and recovery was followed over the remaining 194 cycles. Approximately 20 foci were analyzed per strain. Structures that moved out of the focal plane during the 200-cycle series were discarded.

### U2OS cell culture and transfection

U2OS cells (ATCC, HTB-96, Manassas, VA, USA) were cultured in Dulbecco’s Modified Eagle Medium (DMEM with high glucose; Cytiva, USA SH30023.01) supplemented with 10% fetal bovine serum (FBS; Cytiva, USA, SH30088.03) and 1X antibiotic-antimycotic solution (Anti-Anti; Gibco, USA, 15240-062) at 37°C in a humidified incubator with 5% CO₂. Once the cells reached 70% confluence, they were transfected with pcDNA3.1-myc-hisA expression vectors encoding GFP-TDP-43 and its charge-cluster deletion, charge-neutralization and PAM substitution mutants using Lipofectamine 3000 (Invitrogen, USA, L300015), following the manufacturer’s protocol. After transfection, the cells were selected with 600 ng/μL Geneticin™ Selective Antibiotic (G418 Sulfate) (Gibco, USA, 10131035).

### Fixation and DAPI staining of U2OS cells

At 48 h post-transfection, U2OS cells grown on coverslips were fixed with 4% paraformaldehyde (PFA; T&I, South Korea, BPP-9004) for 15 min at room temperature, followed three washes with phosphate-buffered saline (PBS; WELGENE, South Korea, LB001-02). Cells were permeabilized with 0.2% Triton X-100 for 5 min at room temperature and subsequently incubated with DAPI (1 mg/mL stock solution) diluted 1:1,000 in PBS for 5 min. Coverslips were mounted in VECTASHIELD Mounting Medium for Fluorescence (Vector Laboratories, Burlingame, CA, USA; H-1000) and subjected to fluorescence imaging using a Zeiss LSM980 laser scanning confocal microscope equipped with a 20× objective (Carl Zeiss AG, Oberkochen, Germany).

### FRAP in living U2OS cells

FRAP was performed in living U2OS cells expressing GFP-tagged TDP-43ΔNLS constructs on a Zeiss LSM900 laser scanning confocal microscope (Carl Zeiss AG, Oberkochen, Germany) fitted with a 20× objective; no immersion oil was used. GFP was excited with the 488 nm laser and detected with a GaAsP-PMT detector, with the pinhole set to 3 Airy units. Images were acquired at a frame size of 512 × 512 pixels, 8 bits per pixel, scan speed 9 (approximately 1 μs pixel dwell time), unidirectional scanning and without frame averaging. Each time series comprised 200 cycles acquired continuously, with no added delay between cycles, giving a frame interval of 0.738 s and a total observation window of 146.8 s. The entire condensate was selected as the bleach ROI. Five pre-bleach images were acquired, after which the ROI was bleached with the 488 nm laser at 60–70% power using 20 bleaching iterations, and recovery was followed over the remainder of the series. Twenty condensates were analyzed per construct. Condensates that moved out of the focal plane during acquisition were excluded. Imaging was performed at room temperature; because all constructs were acquired under identical conditions, recovery parameters are reported and compared as relative values rather than as absolute measures of condensate dynamics.

### Image analysis for FRAP

The mean fluorescence intensity within the ROI at each cycle was extracted in Fiji using the Stowers plugin. Background was not subtracted, and curves were not corrected for photobleaching occurring during acquisition; all constructs were acquired and analyzed identically, so comparisons between them are unaffected. Each curve was normalized to the mean intensity of its own five pre-bleach frames, and the normalized curves were fitted in GraphPad Prism 11 (GraphPad Software, CA, USA) with the one-phase association model of Eq. 1.

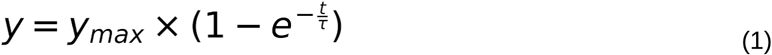

In Eq. 1, y_max_ is the value of y at infinite time (t ≈ ∞) and τ (tau) is the time at which y reaches 63.2% of y_max_. The mobile fraction was taken as the fitted plateau y_max_ and is expressed as a percentage of the pre-bleach intensity; the recovery half-time (t½) was calculated from the rate constant of the same fit as t½ = τ × ln2 and is expressed in seconds. Average curves were calculated and plotted in GraphPad Prism 11.

### Proteostat staining

Protein aggregates in yeast and U2OS cells were visualized using the PROTEOSTAT Aggresome Detection Kit. For yeast, cells fixed with 4% paraformaldehyde for 1 h at room temperature and washed three times with 1 mL PBS were subsequently permeabilized with 0.2% Triton X-100 for 20 min at room temperature and washed three times with PBS. PROTEOSTAT reagent diluted 1:2,000 was applied for 1 h at room temperature in the dark, followed by three PBS washes. Images were acquired using a Zeiss LSM900 laser-scanning confocal microscope (Carl Zeiss AG, Oberkochen, Germany).

U2OS cells, 48 h after transfection, were fixed with 4% paraformaldehyde for 30 min at room temperature. Following two PBS washes, cells were permeabilized with 500 μL of permeabilization solution on ice for 30 min with washes in PBS, and incubated with 750 μL of staining solution containing PROTEOSTAT reagent (diluted 1:5,000) and Hoechst (diluted 1:1,000) for 30 min at room temperature in the dark. Images were acquired using a Zeiss LSM980 laser-scanning confocal microscope (Carl Zeiss AG, Oberkochen, Germany).

### Transmission electron microscopy of yeast cells

Yeast cells were processed by conventional permanganate fixation, adapted from previously published protocols (Mayer et al., 2024; Seel et al., 2023). After 24 h of galactose induction, cells were harvested by centrifugation, resuspended in fixative containing 2% paraformaldehyde (EM grade, EMS) and 2.5% glutaraldehyde (EMS) in 0.1 M cacodylate buffer (pH 7.2), and fixed for 1 h at 4 °C. After three washes in 0.1 M cacodylate buffer (pH 7.2), an equal volume of molten 2% agar was added to the pellet and the sample was cooled on ice to solidify. The solidified sample was removed from the tube and cut into pieces of approximately 1–2 mm, which were incubated in 1 mL of freshly prepared 4% KMnO₄ for 1 h at room temperature. Blocks were washed four to six times in distilled water until the supernatant was almost colorless, incubated in freshly prepared 0.5% NaIO₄ for 15 min at room temperature, and washed three times for 5 min in distilled water. Blocks were then incubated overnight in 2% uranyl acetate, dehydrated through a graded acetone series (50, 60, 70, 80, 90 and 100%; 20 min at each step) and infiltrated with embedding medium (EM812, EMS). Sections of 60 nm were cut on an ultramicrotome (Leica Ultracut UCT; Leica Microsystems, Vienna, Austria), mounted on copper slot grids with a specimen support film, and double-stained with UranyLess (EMS, #22409) for 2 min and 3% lead citrate (EMS, #22410) for 1 min. Sections were imaged on a transmission electron microscope operated at 120 kV (Tecnai G2, ThermoFisher, USA).

### Western blotting analysis

Protein extraction from yeast cells was obtained using the alkaline lysis as described in Choi and Song (2022). U2OS cell extract was obtained as previously described in Goh et al. (2024). The western blot analyses were performed essentially as described in Choi and Song (2022) and Goh et al., (2024) with anti-GFP antibody (600-101-215 ROCKLAND, Limerick, PA, USA, 1:1000 dilution). The blots were stripped and re-probed with a mouse anti-tubulin antibody (Sigma-Aldrich, Cat# T5168, 1:10000 dilution) for loading controls. Bound antibodies were detected with horseradish peroxidase-conjugated secondary antibodies: anti-goat (705-035-003, Jackson ImmunoResearch, West Grove, PA, USA; 1:10,000 dilution) for GFP and anti-mouse (sc-2005, Santa Cruz Biotechnology, Dallas, TX, USA; 1:10,000 dilution) for Tub1.

### Quantification and Statistical Analysis

Box plots are Tukey plots: the box spans the interquartile range with the median shown as a line and the mean as a cross, the whiskers extend to the most extreme value lying within 1.5 × the interquartile range of the box, and values outside that range are plotted individually; n denotes the number of bleached structures and is stated in each figure legend. Statistical analyses were performed using GraphPad Prism. FRAP parameters were compared by Brown-Forsythe and Welch ANOVA, which does not assume equal variances across groups. Post hoc tests were selected according to the structure of the comparison: Dunnett’s T3 test where every group was compared with a single control, and Dunnett’s T3 restricted to comparisons planned a priori (Figure 5C, 5D). Because the condensate categories in Figure 5 form an ordered series, the overall trend in mobile fraction across categories 1, 2, 3-1, 3-2 and 4 was additionally assessed by the Jonckheere–Terpstra test for monotone trend (two-sided, normal approximation with tie correction), implemented in Python 3.10 with NumPy and SciPy. This test evaluates the ordered alternative directly and does not require adjacent categories to differ from one another pairwise. The same test was applied to the PA2 charge-neutralization series in Figure S1F, with the constructs ordered by the number of acidic residues neutralized (WT, PA2-2, PA2-1 and PA2); ΔPA2 was not included, because deletion removes the cluster rather than a defined number of charges. P < 0.05 was considered statistically significant.

### Data and code availability

Algorithms were implemented in Python 3.10 by the authors. The analysis pipeline included modules written with NumPy and SciPy and was used to process experimental data shown in Figure 1 and to perform the trend test reported for Figure 5 and Figure S1. The code used in this study was written by the authors and is available from the corresponding author upon reasonable request.

### KEY RESOURCES TABLE

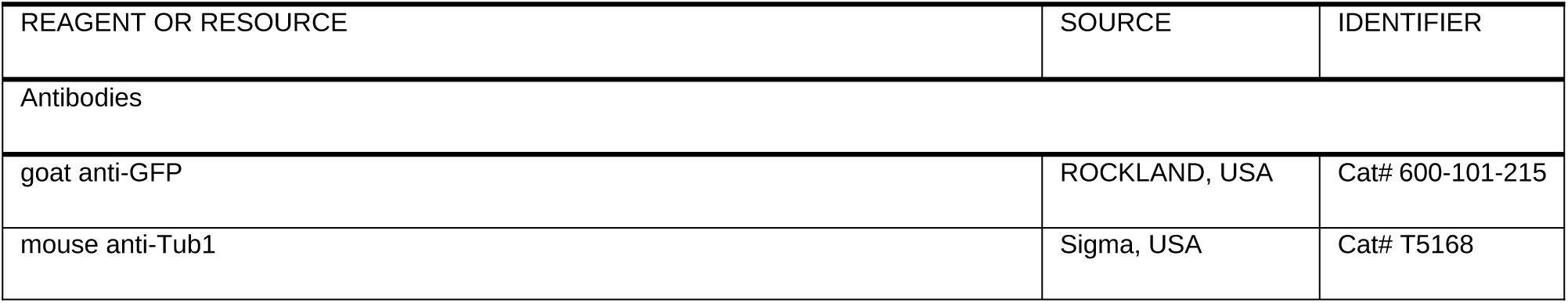
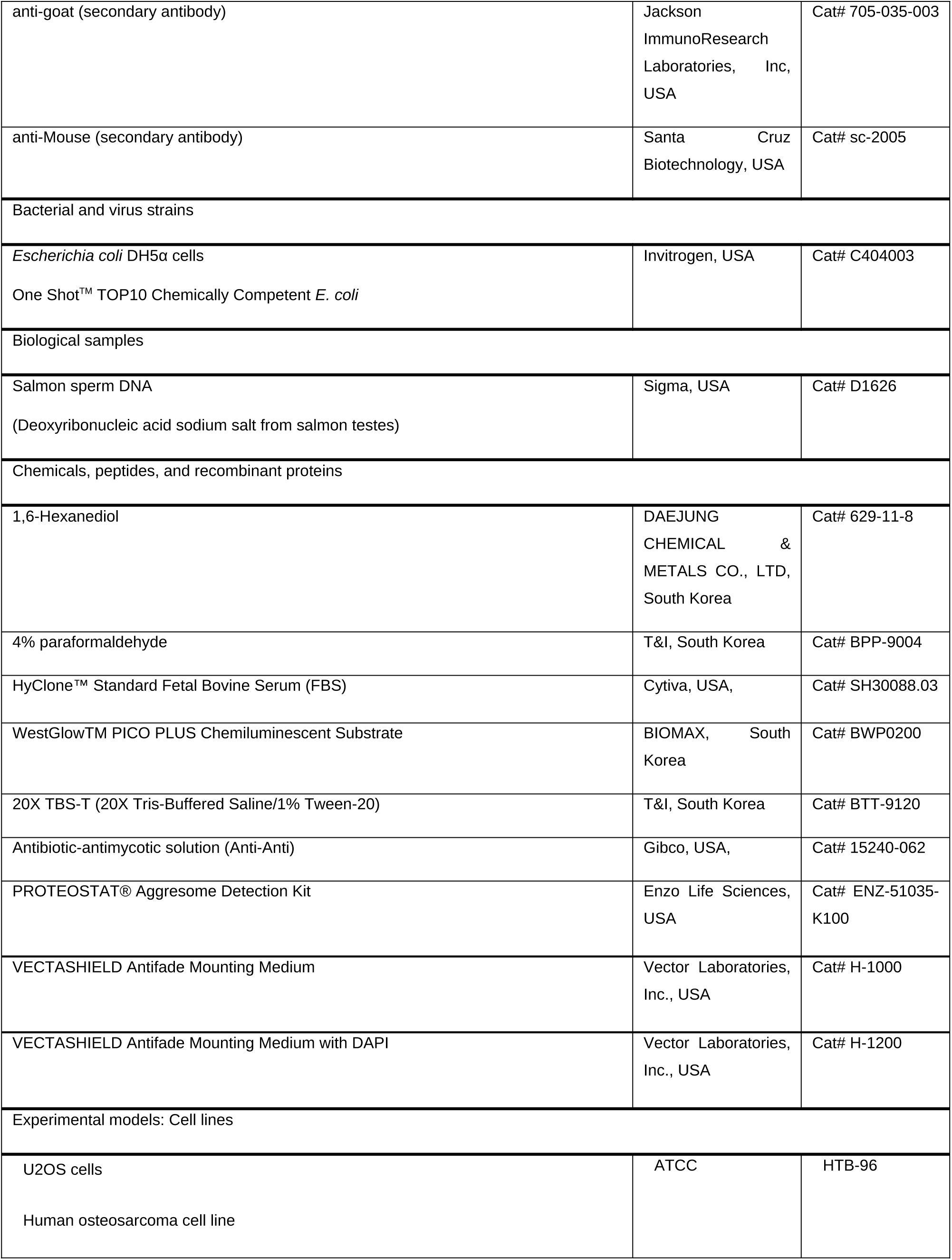
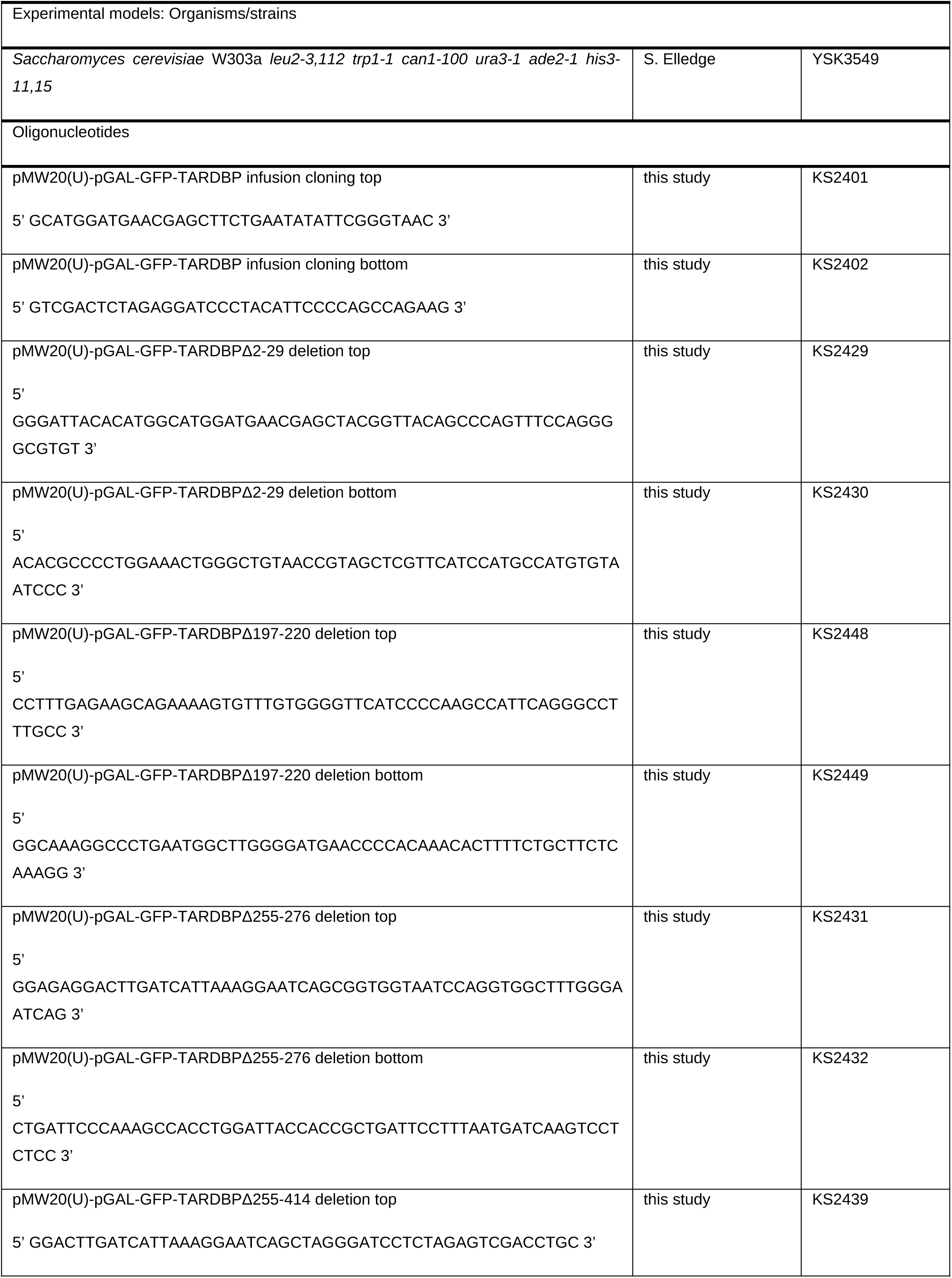
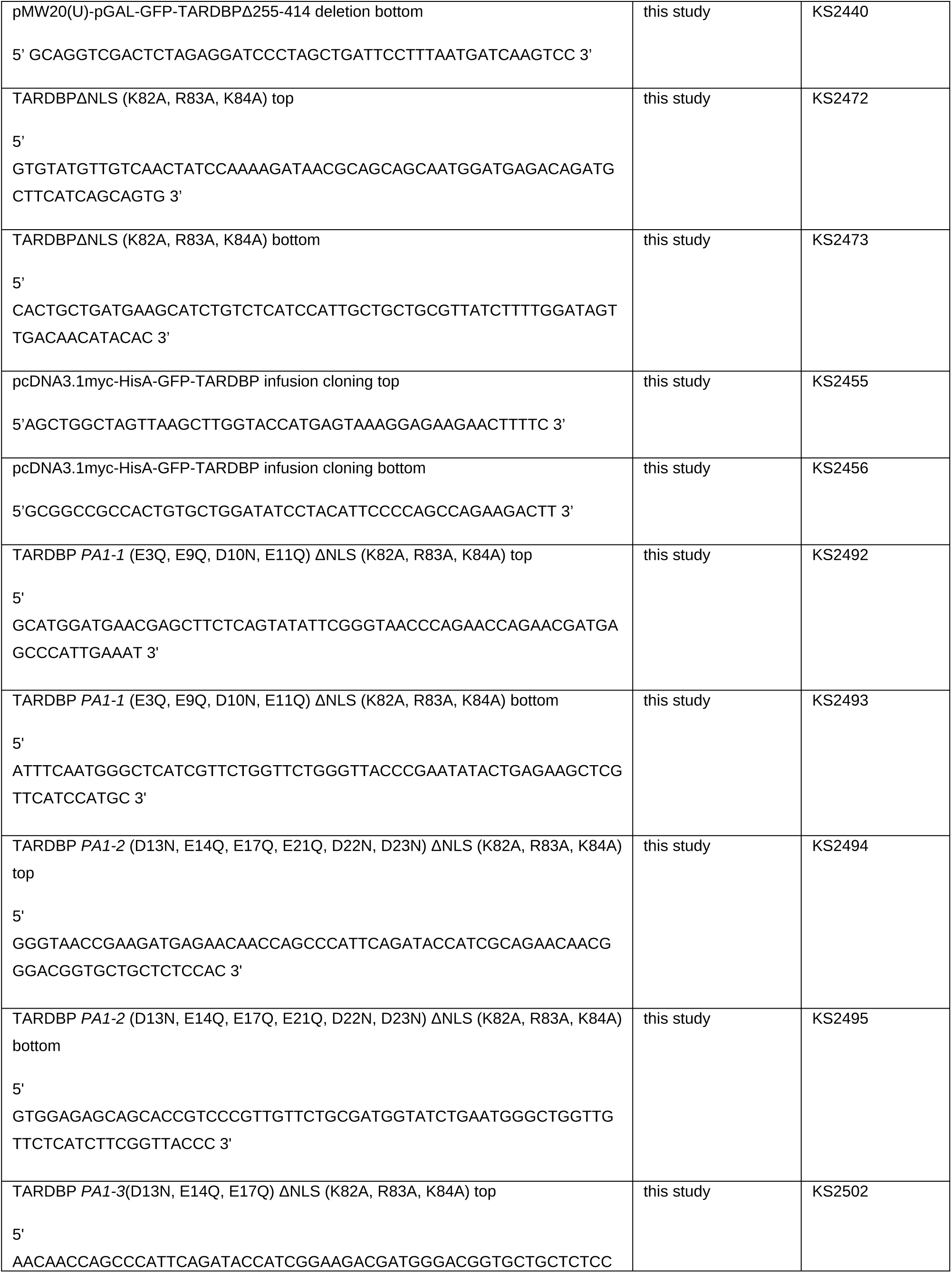
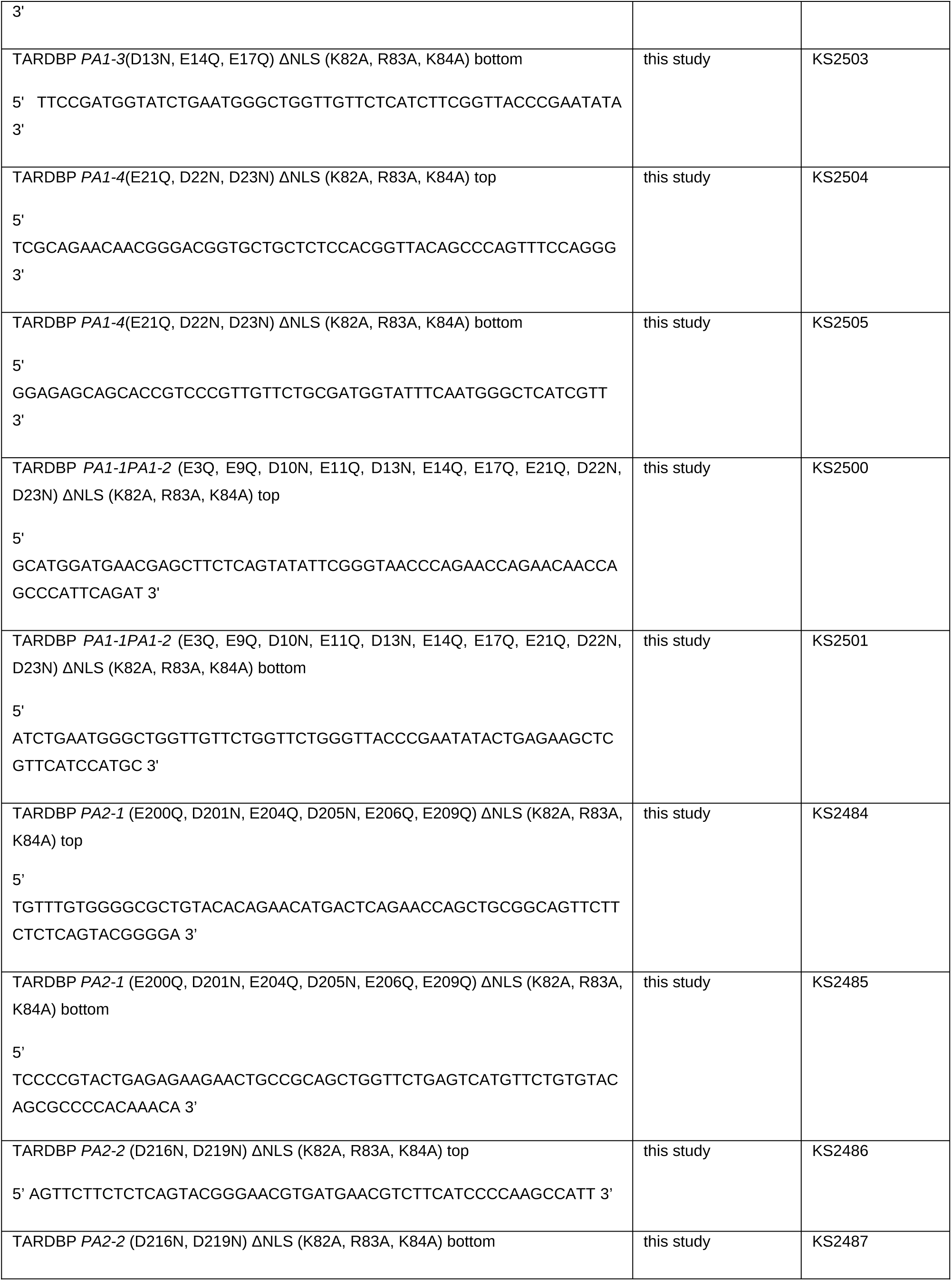
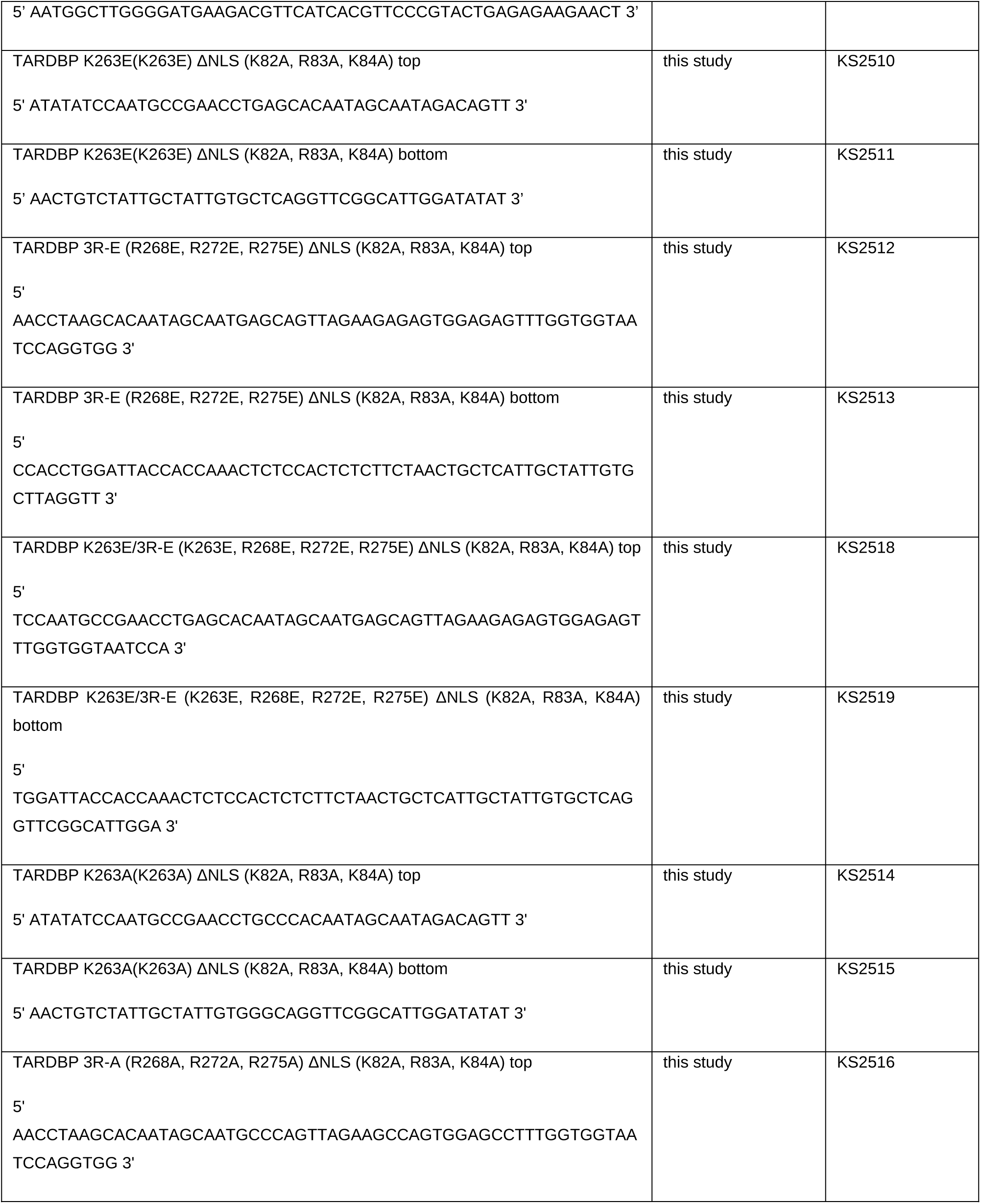
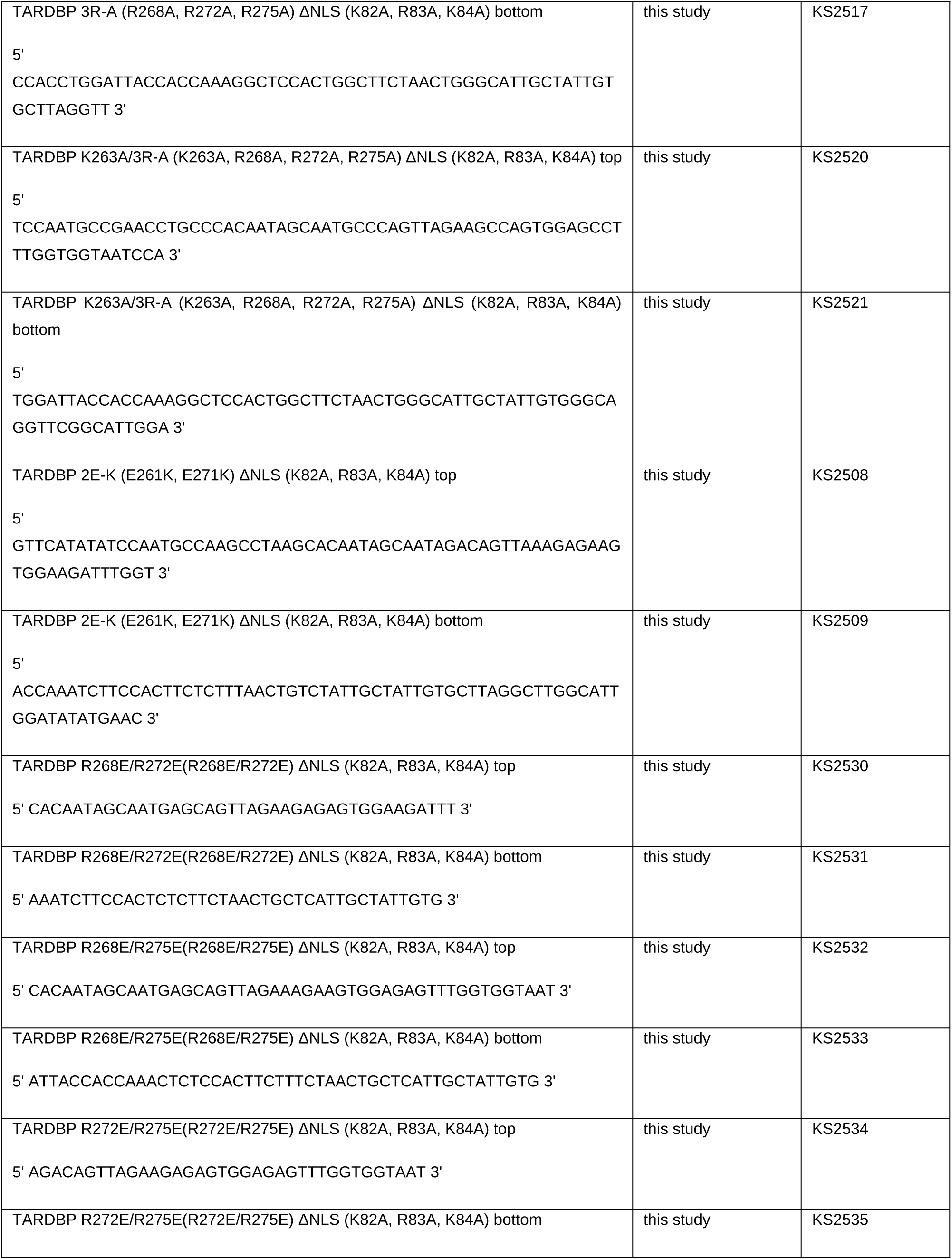
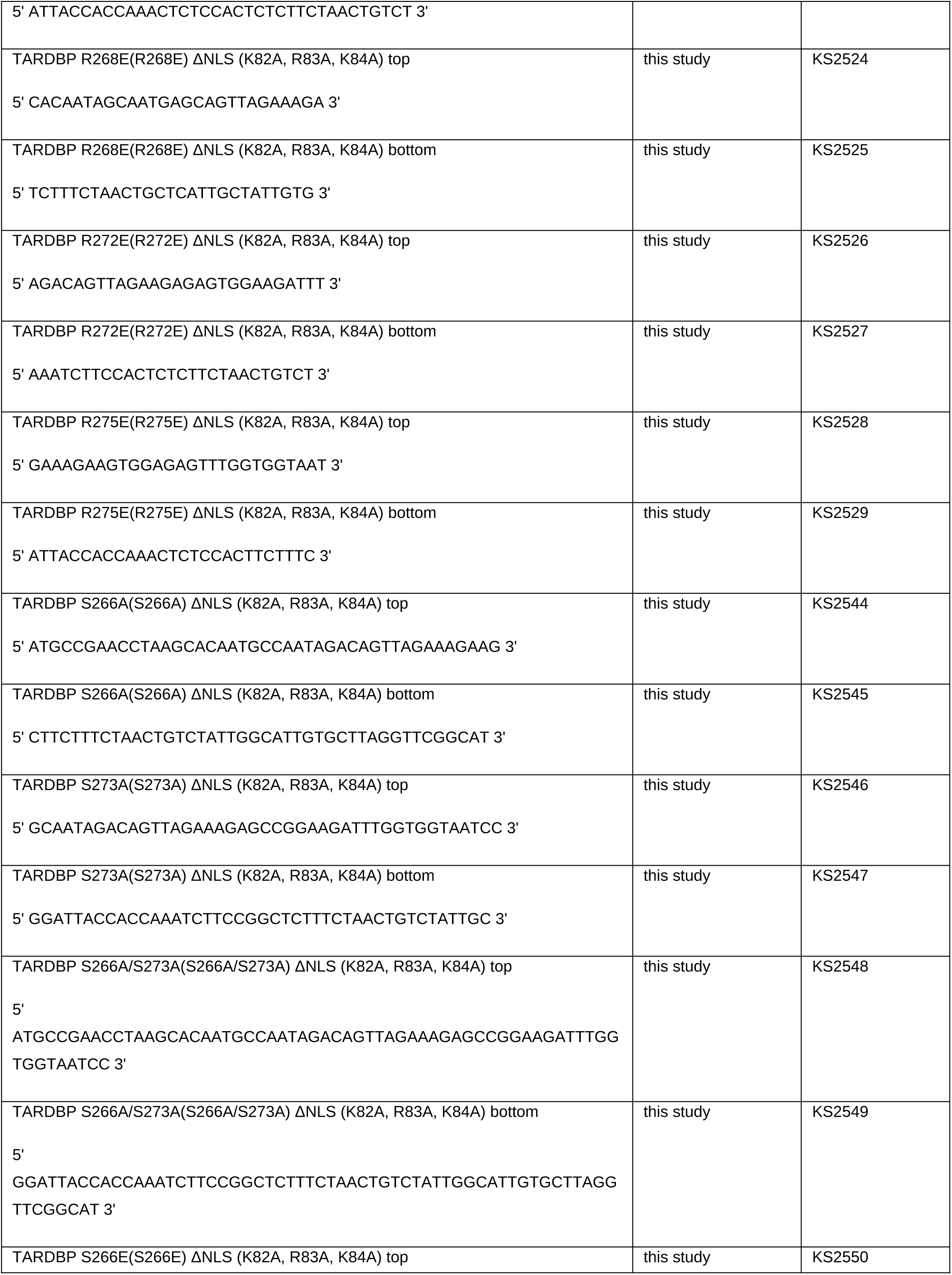
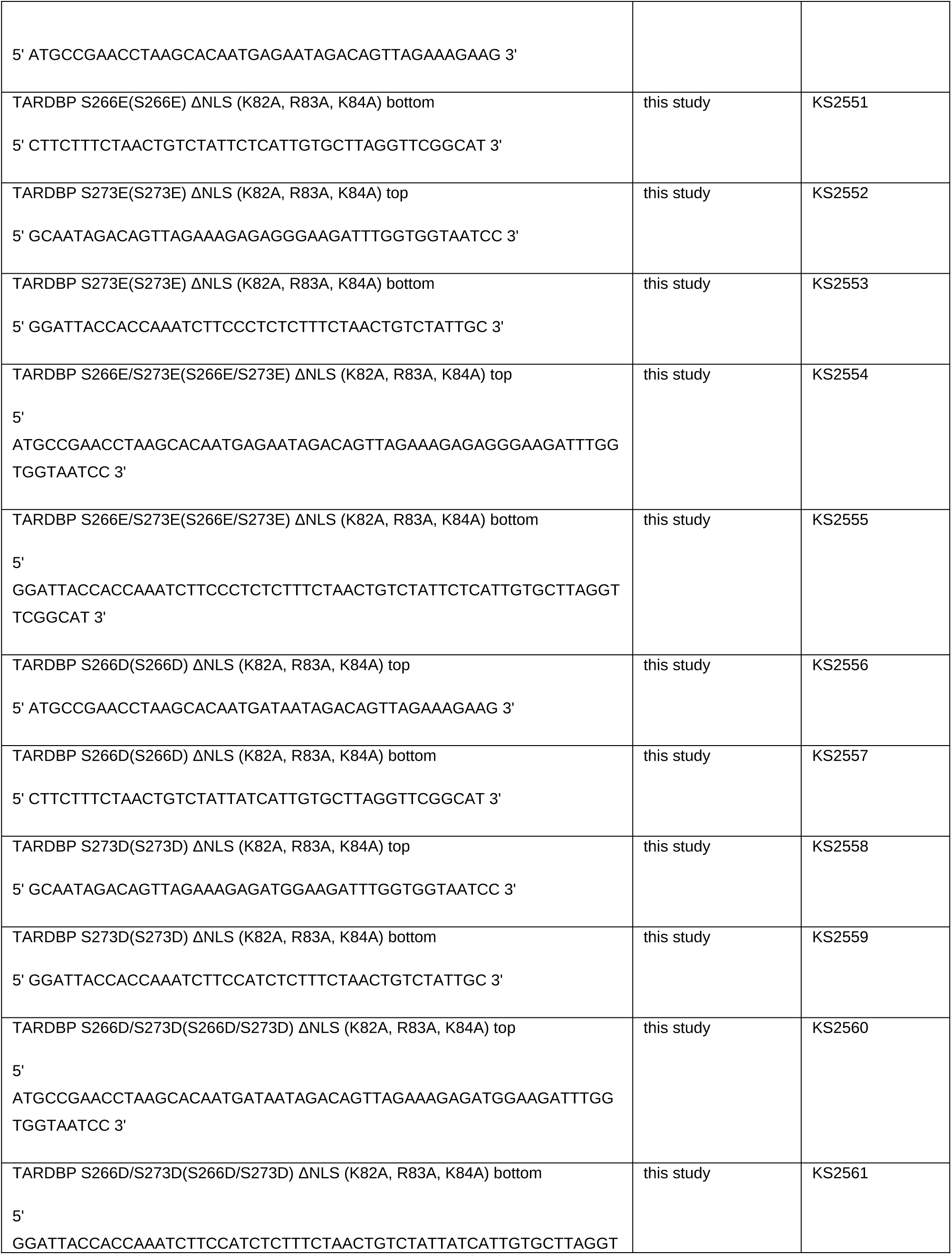
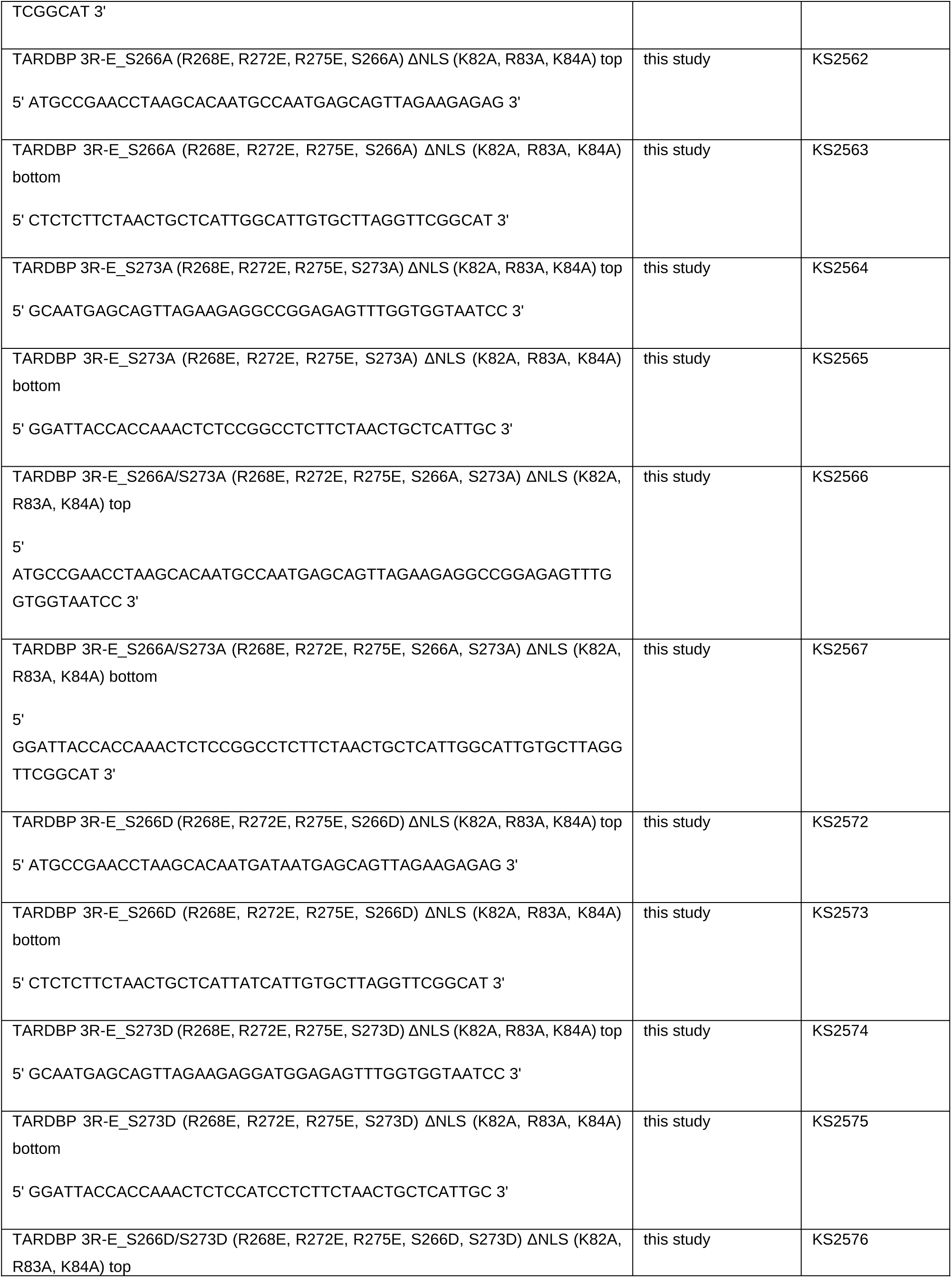
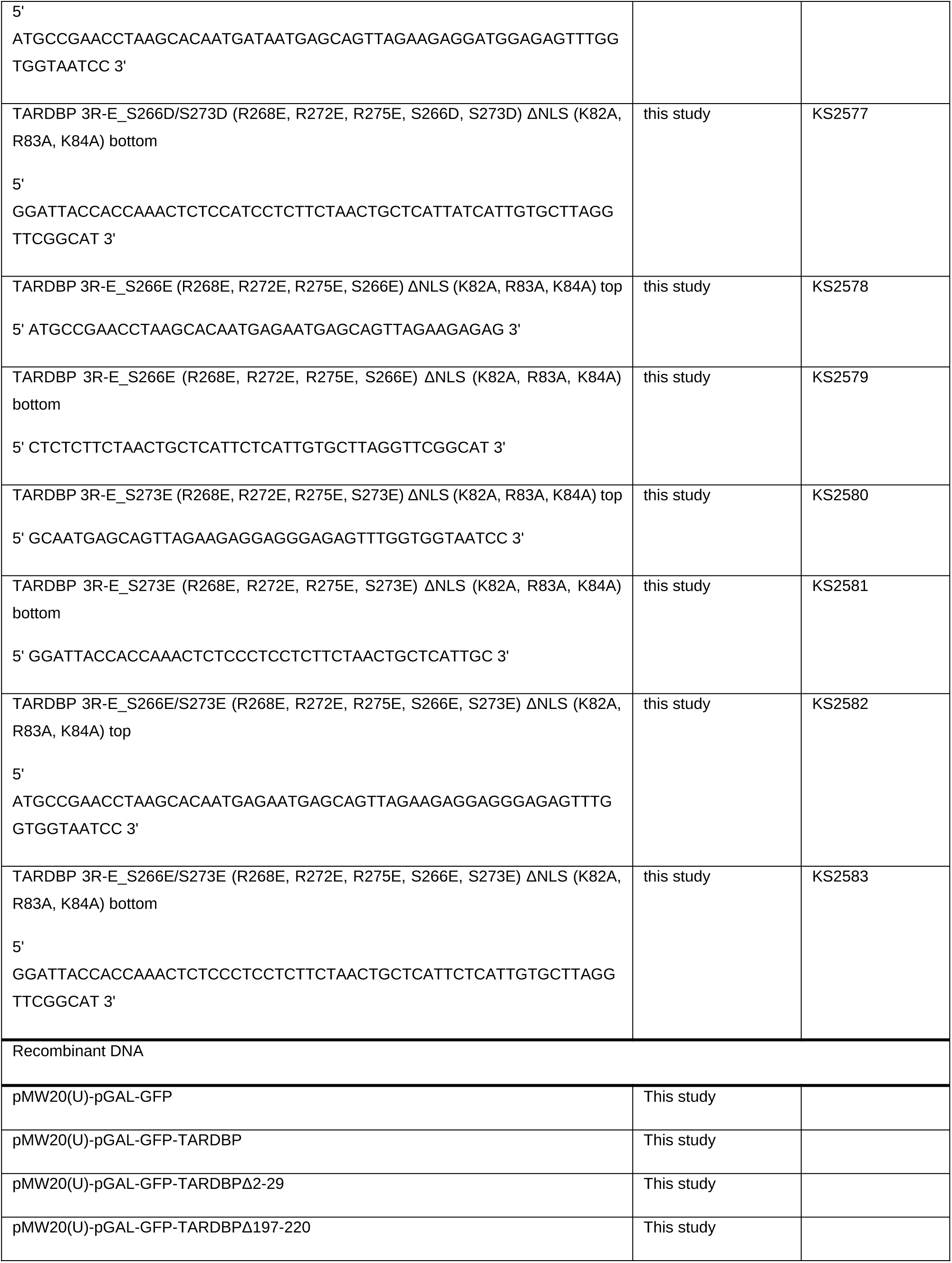
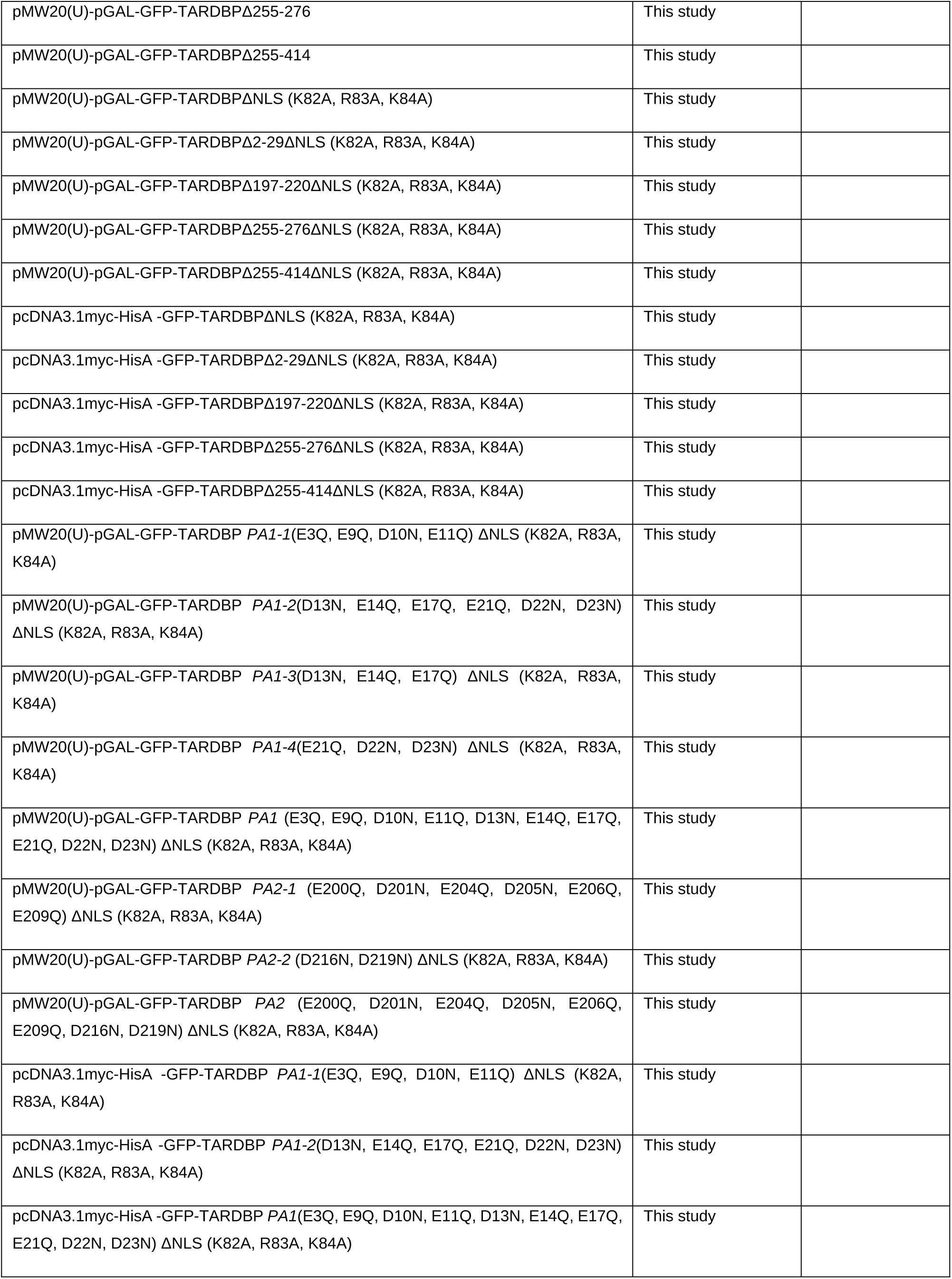
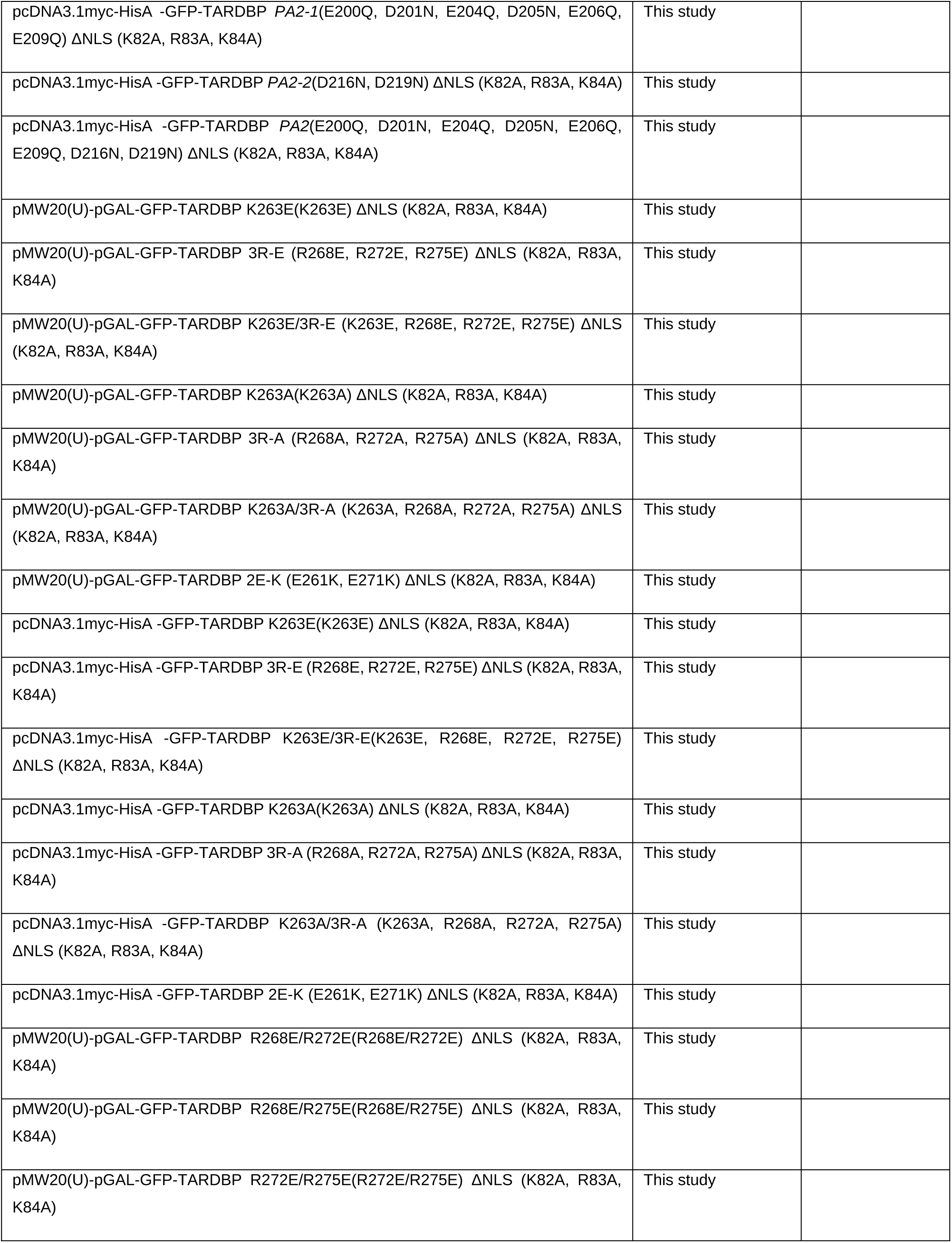
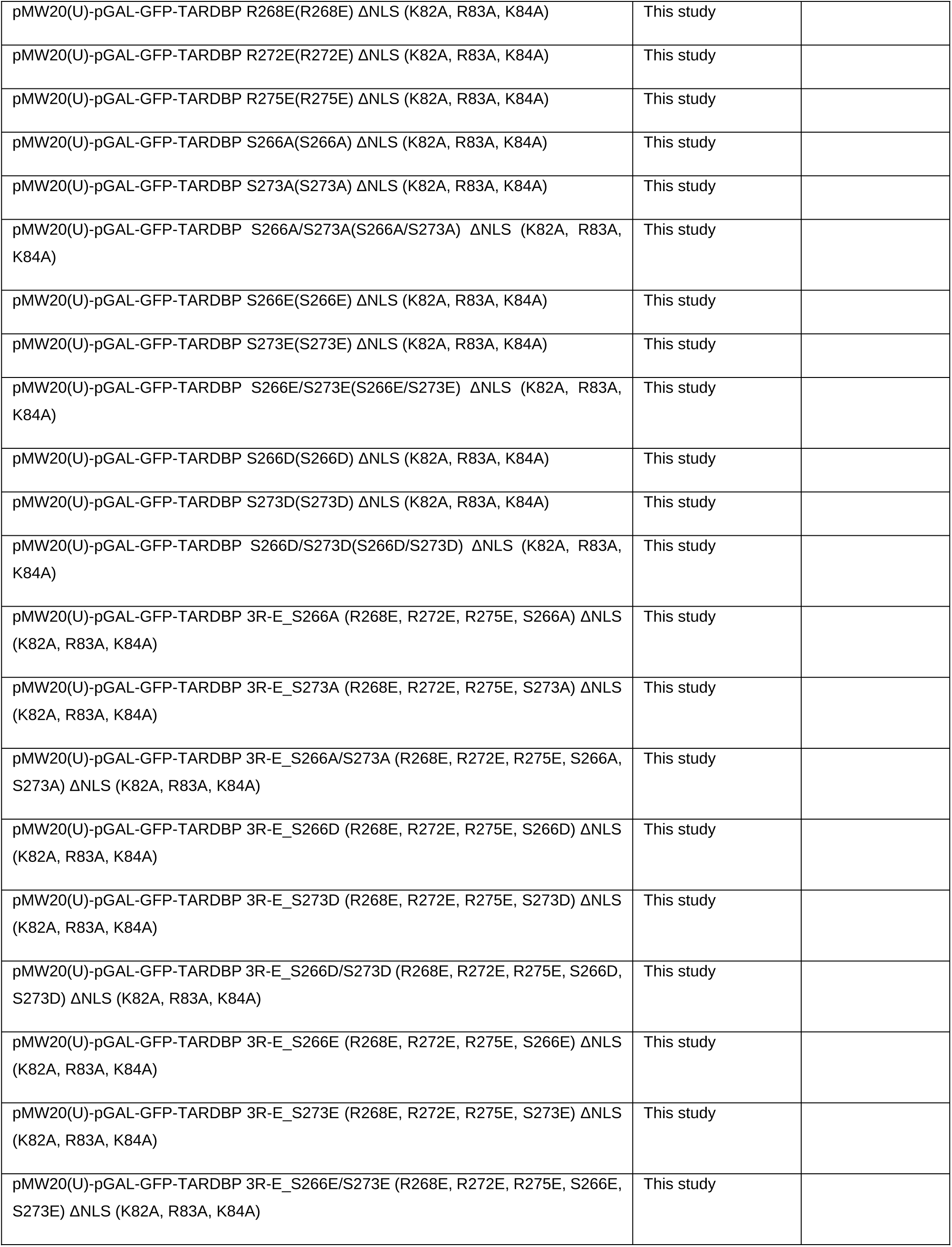
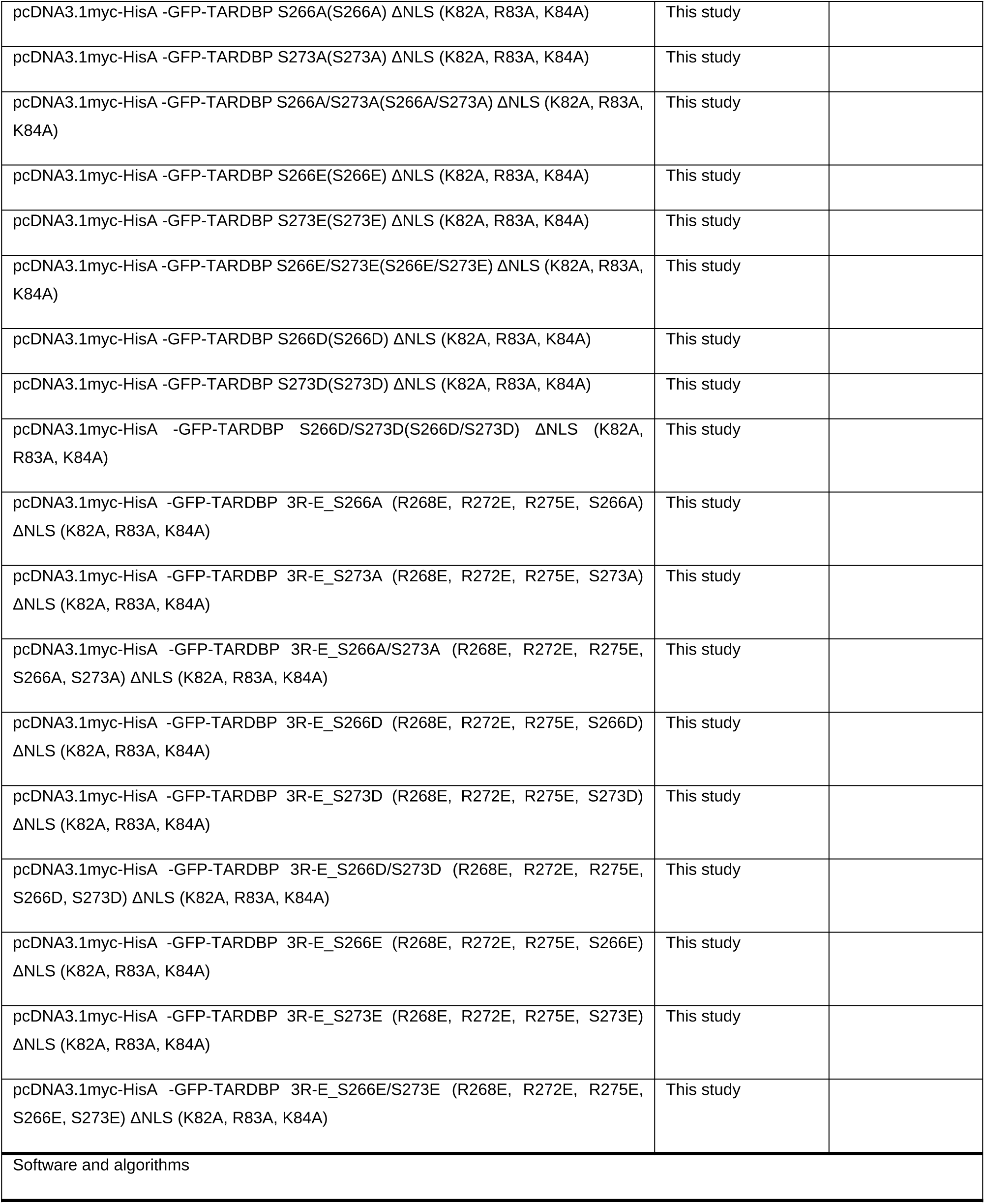
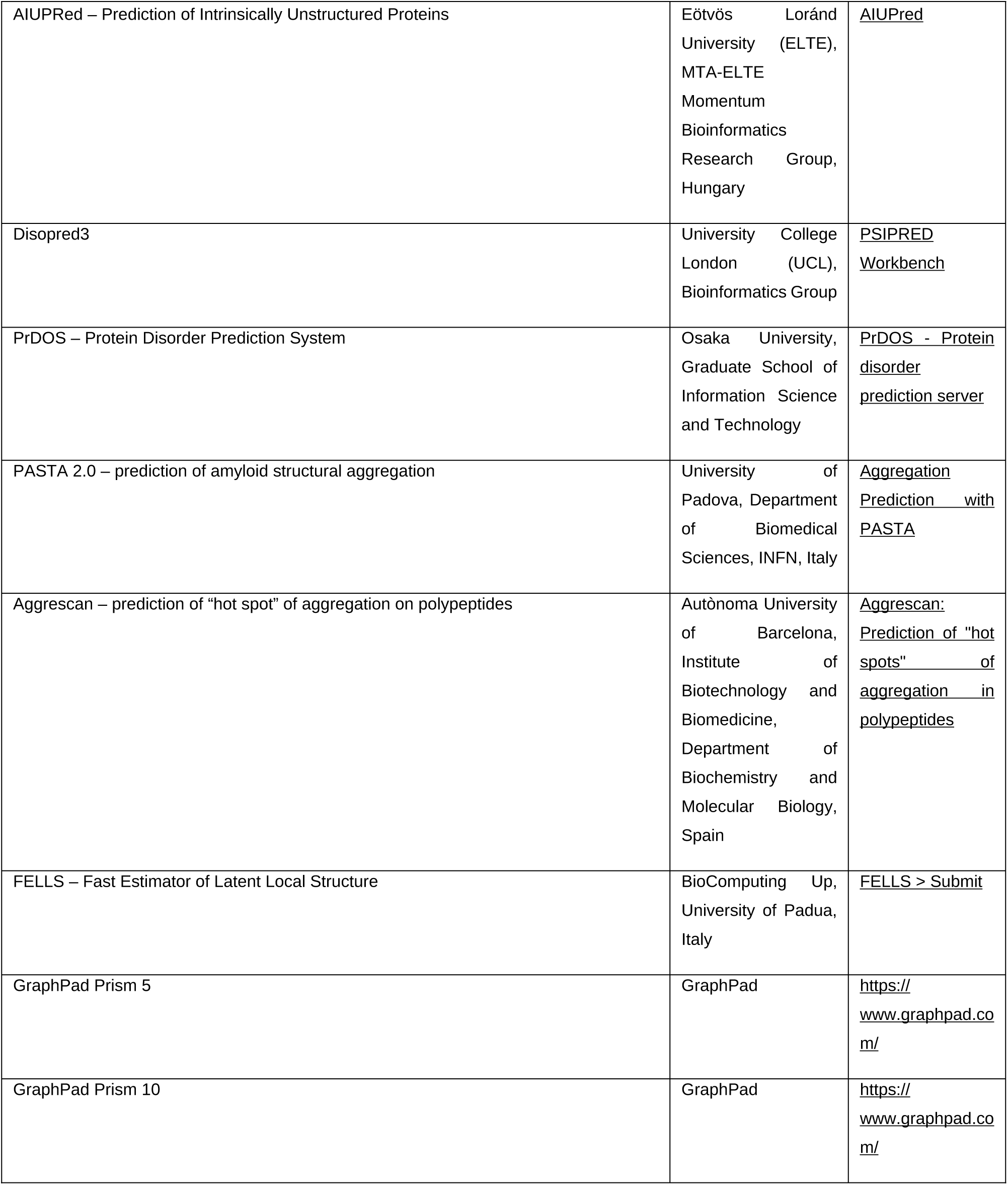
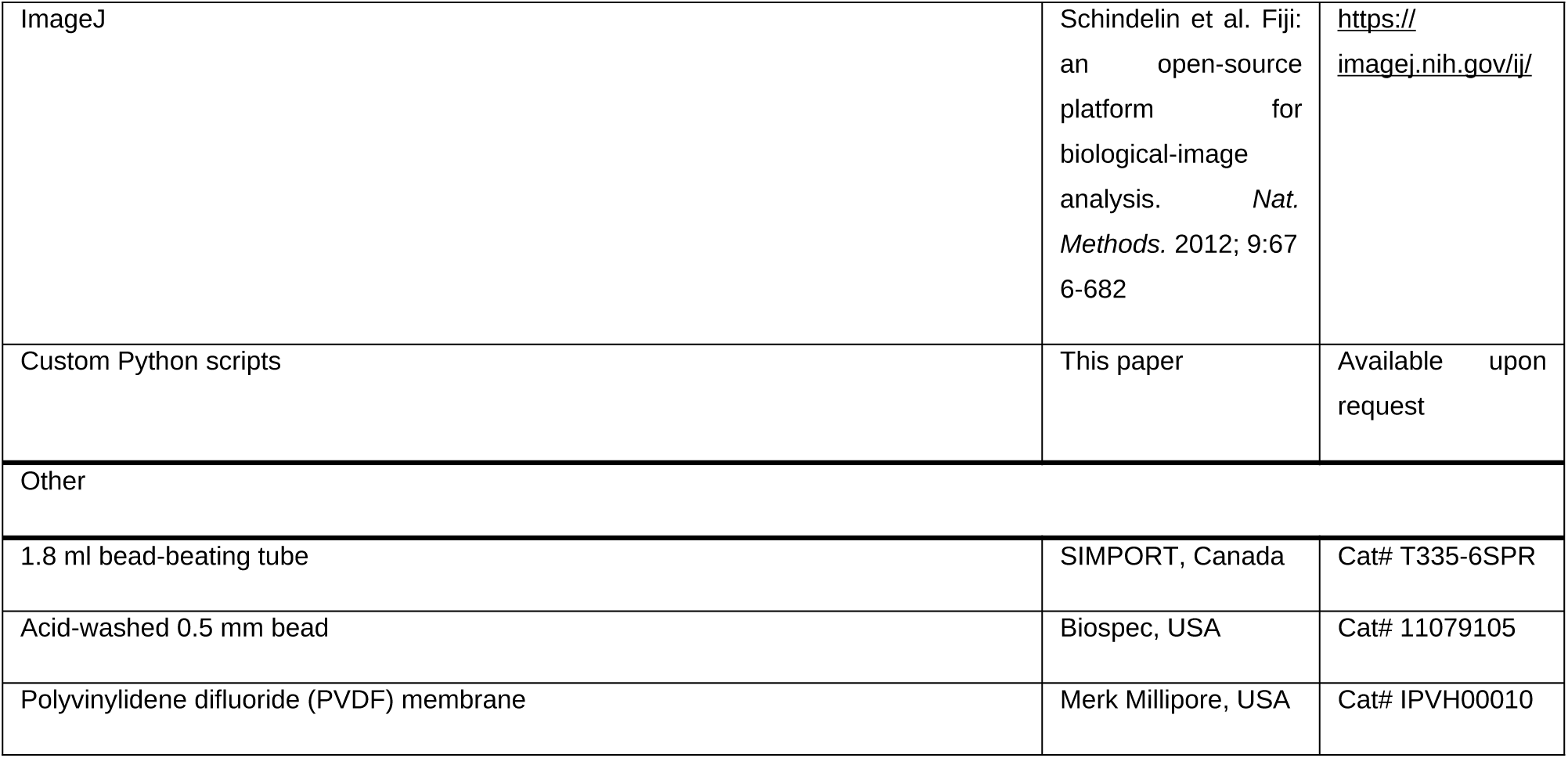

## ACKNOWLEDGMENTS

We thank the Cellular Imaging Core Facility at the ConveRgence mEDIcine research cenTer (CREDIT), Asan Medical Center, for technical support. The authors also thank Ewha Women’s University Fluorescence Core Imaging Center for valuable guidance in FRAP analysis. We specially appreciate the technical assistance provided by professor Dongmin Kang, the Director, and Research Technician Sinae Jang, of Ewha Fluorescence Core Imaging Center. TDP-43 gene clone was kindly provided by Dr. Hyung-Jun Kim in Korea Brain Research Institute. Y.J.C. was supported in part by the National Research Foundation of Korea (NRF) grant funded by the Korea government (MSIT) (RS-2024-00451507). Y.K. was supported in part by the Basic Science Research Program through the National Research Foundation of Korea (NRF) funded by the Ministry of Education (RS-2025-25408081). M.J. was supported by the National Research Foundation of Korea (NRF) grant funded by the Korea government (MSIT) **(RS-2025-16067602).**

**S.C. and S.J. were supported by the BK21 FOUR Program funded by the Ministry of Education (MOE, Korea) and the National Research Foundation of Korea (NRF).**

## AUTHOR CONTRIBUTIONS

Conceptualization, K.S. and Y.J.C.; methodology, K.S., Y.J.C., M.J., Y.J., and C.P.; Investigation, Y.J.C., Y.K., S.C., S.J., and M.J.; writing—original draft, K.S. and Y.J.C.; writing—review & editing, K.S., Y.J.C., C.P., and Y.J.; funding acquisition, K.S., M.J.; resources, K.S.; supervision, K.S. and Y.J.C.

## DECLARATION OF GENERATIVE AI AND AI-ASSISTED TECHNOLOGIES

During the preparation of this work the authors used Claude (Anthropic) in order to edit and condense the manuscript text and figure legends. After using this tool, the authors reviewed and edited the content as needed and take full responsibility for the content of the published article.

