## Supplemental figures for "TDP-43 amyloid conversion in cells tracks the net charge of a polyampholyte motif, not condensate solidification"

**Supplementary figures**

Figure S1

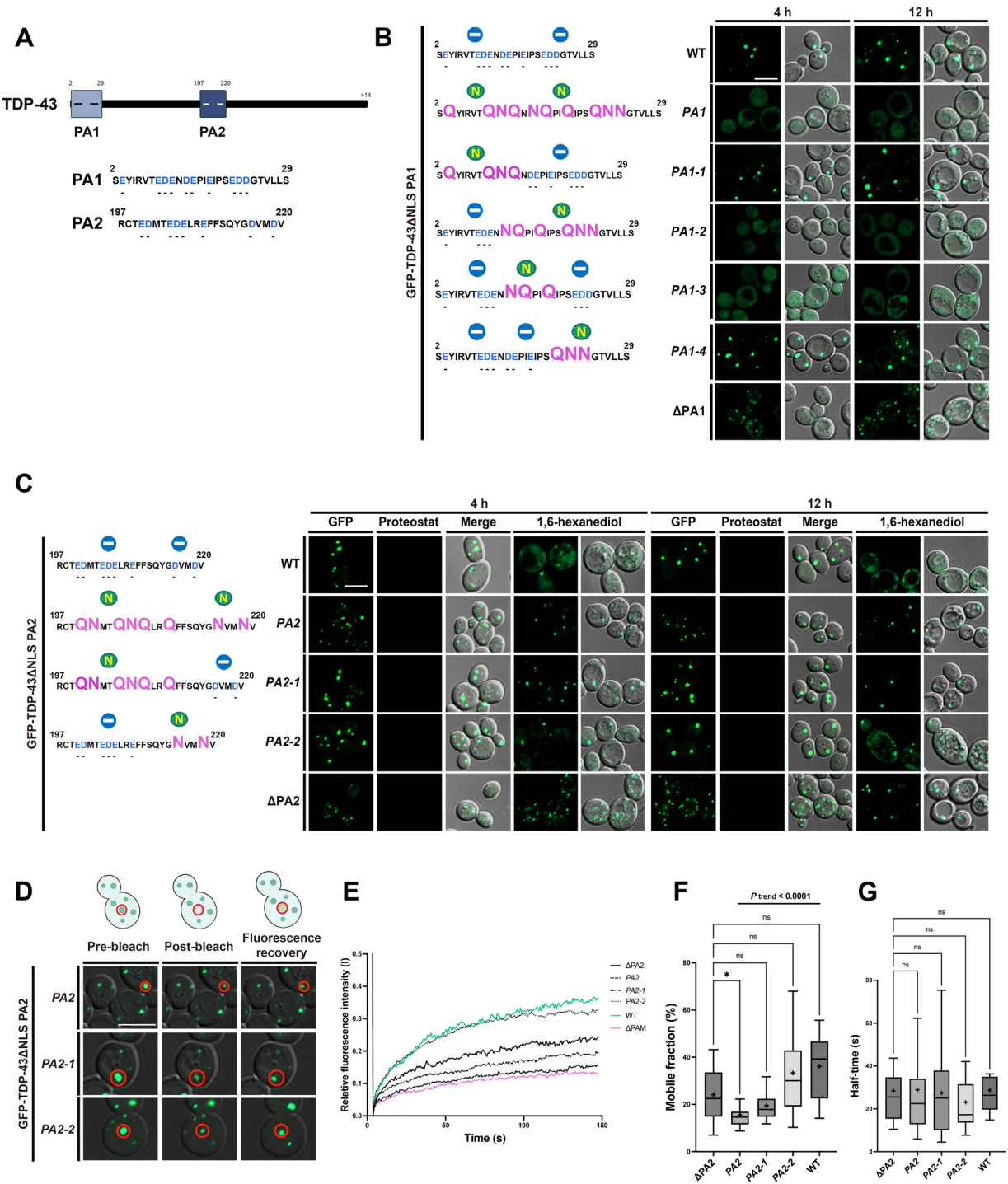

**Figure S1. The contribution of each anionic cluster of TDP-43 maps to its internal acidic patches,** **related to Figure 1**

(A) Schematic representation of the two polyanion clusters of TDP-43. PA1 (residues 2–29) and PA2 (residues 197–220), two negatively charged regions of TDP-43, are indicated, and their amino acid sequences are shown. Negatively charged amino acid residues are marked in blue.

(B) Representative confocal fluorescence microscopy images of budding yeast (W303a) cells expressing GFP-TDP-43ΔNLS and its PA1 charge-neutralization mutants *PA1*, *PA1-1*, *PA1-2*, *PA1-3* and *PA1-4*, together with ΔPA1, after 4 or 12 h of galactose induction. The corresponding amino acid sequences for WT and each *PA1* residue-substituted mutant are shown to the left of the respective mutant image labels. Representative images are shown from analyses of more than 100 cells for each construct at each time point. Scale bar: 5 μm.

(C) Representative confocal fluorescence microscopy images of budding yeast (W303a) cells expressing GFP-TDP-43ΔNLS and its PA2 charge-neutralization mutants *PA2*, *PA2-1* and *PA2-2*, together with ΔPA2, after 4 or 12 h of galactose induction. For each time point, GFP, ProteoStat and merged channels are shown, followed by cells treated with 1,6-hexanediol. The corresponding amino acid sequences for WT and each *PA2* residue-substituted mutant are shown to the left of the respective mutant image labels. Representative images are shown from analyses of more than 100 cells for each construct at each time point. Scale bar: 5 μm.

(D) Representative confocal images showing pre-bleach, post-bleach, and fluorescence recovery states of condensates formed by the TDP-43ΔNLS PA2 variants *PA2*, *PA2-1* and *PA2-2*. The imaging sequence comprised 200 frames, with the pre-bleach, post-bleach, and recovery images corresponding to frames 1, 6, and 200, respectively. The bleached region of interest (ROI) is marked with a red circle. Scale bar: 5 μm.

(E) Graph displaying the average FRAP recovery curves of the TDP-43ΔNLS PA2 variants together with WT and ΔPAM for reference. The WT, ΔPAM and ΔPA2 measurements are those shown in Figure 1E–1G; the WT and ΔPA2 values also appear in (F) and (G). The x-axis represents time in seconds (200 frames acquired at 0.738 s intervals, 0–150 s) and the y-axis the relative fluorescence intensity. Akima spline interpolation was applied to all FRAP curves, and plotting was performed using GraphPad Prism 11.

(F) Mobile fraction of the puncta shown in (E). For each bleached structure the recovery curve was normalized to its own pre-bleach intensity and fitted with a one-phase association model; the mobile fraction is the fitted plateau, expressed as a percentage of the pre-bleach intensity. $n = 18, 22, 18, 23$ and $14$ structures for $\Delta PA2$, $PA2$, $PA2-1$, $PA2-2$ and $WT$. Statistical comparisons were performed by Brown-Forsythe and Welch ANOVA with Dunnett's T3 post hoc test, with $\Delta PA2$ as the control group. The trend in mobile fraction across $WT$, $PA2-2$, $PA2-1$ and $PA2$, ordered by the number of acidic residues neutralized, was assessed by the Jonckheere–Terpstra test for monotone trend ($P_{\text{trend}} < 0.0001$). $P < 0.05$ was considered statistically significant. ns, not significant; $*P < 0.05$.

(G) Recovery half-time of the same structures shown in (F), taken from the rate constant of the one-phase association fit of each individual curve and plotted in seconds. Statistical comparisons were performed as in (F). ns, not significant.

Figure S2

A

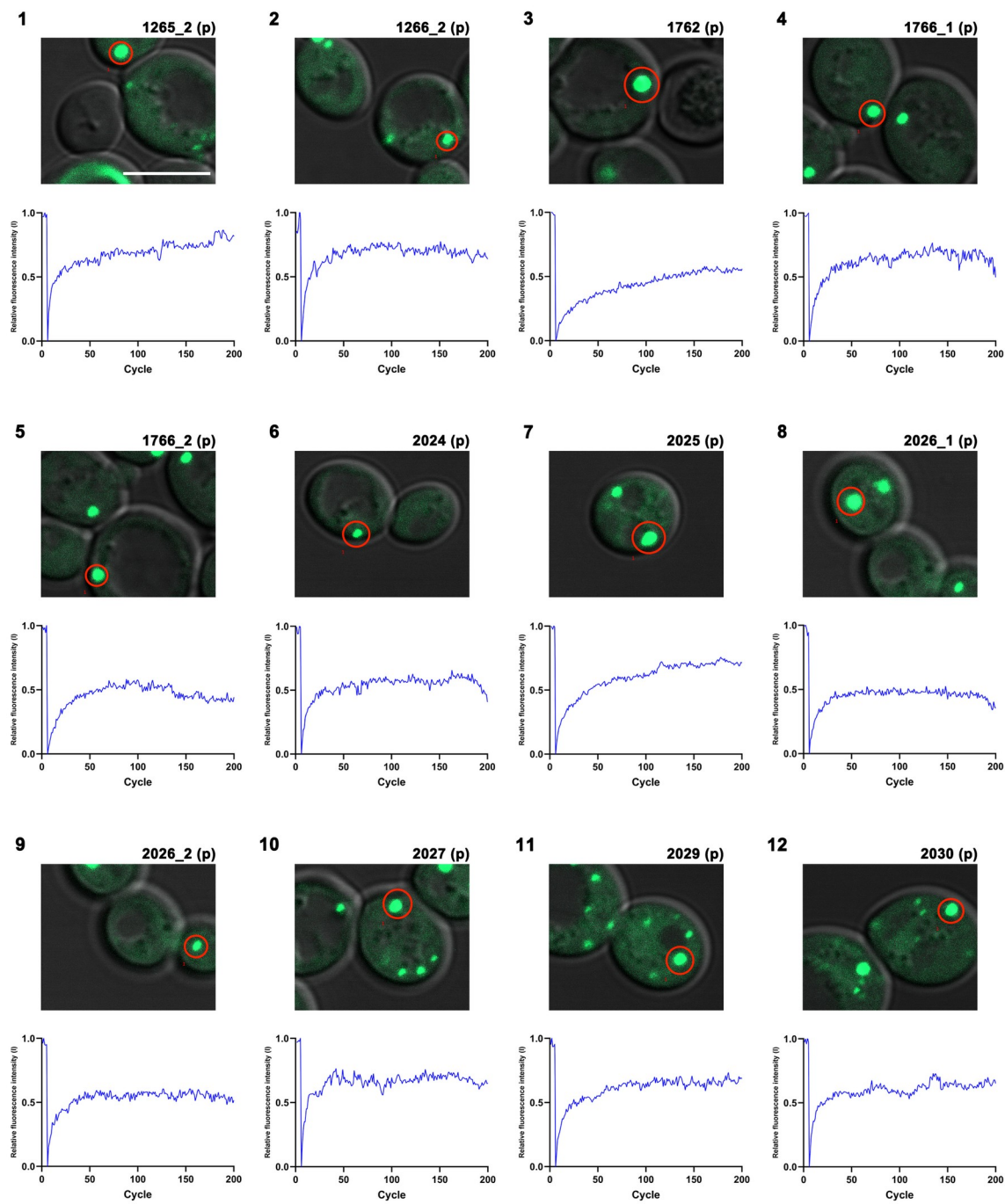

Figure S2

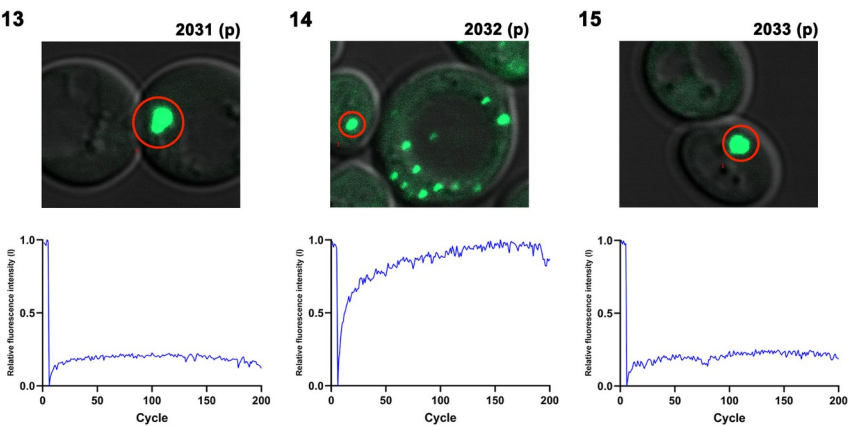

**Figure S2**

**B**

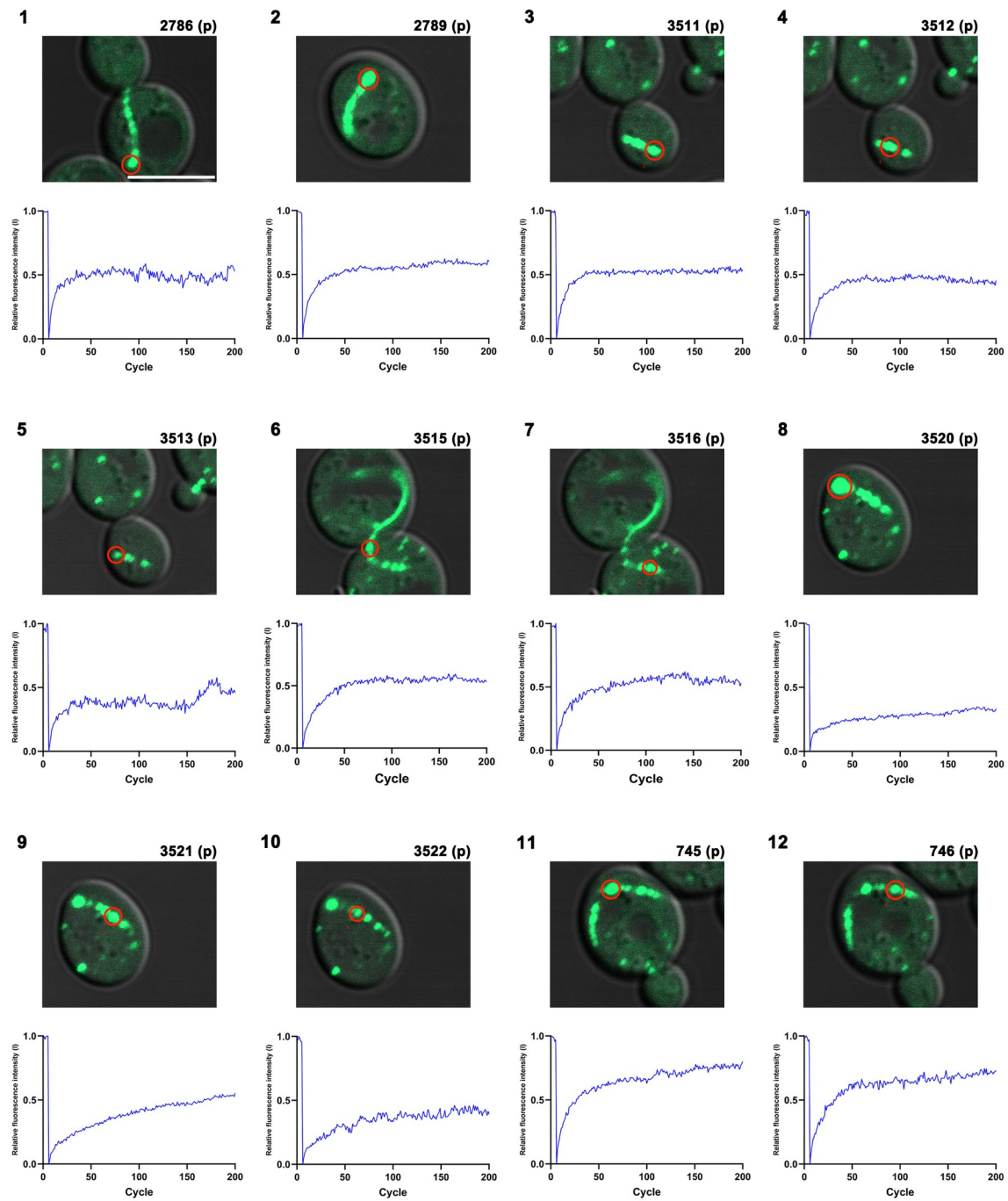

Figure S2

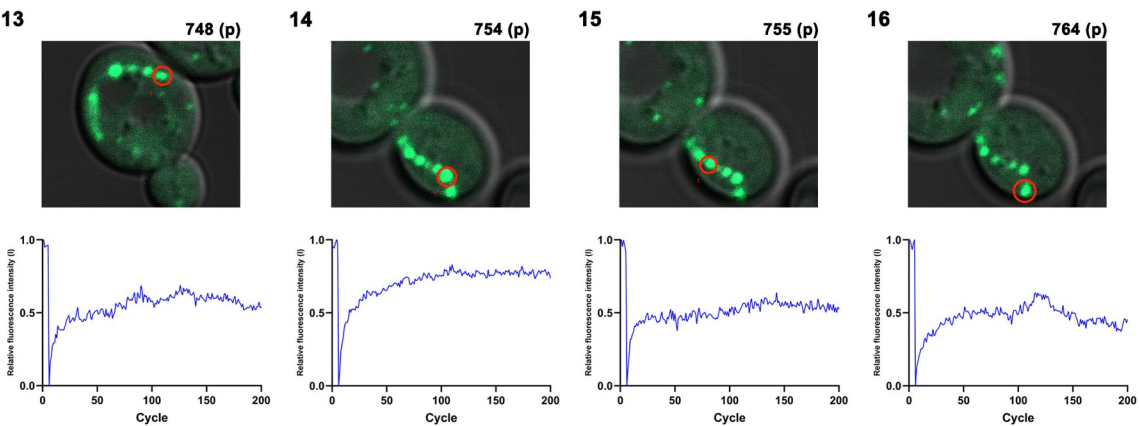

Figure S2

C

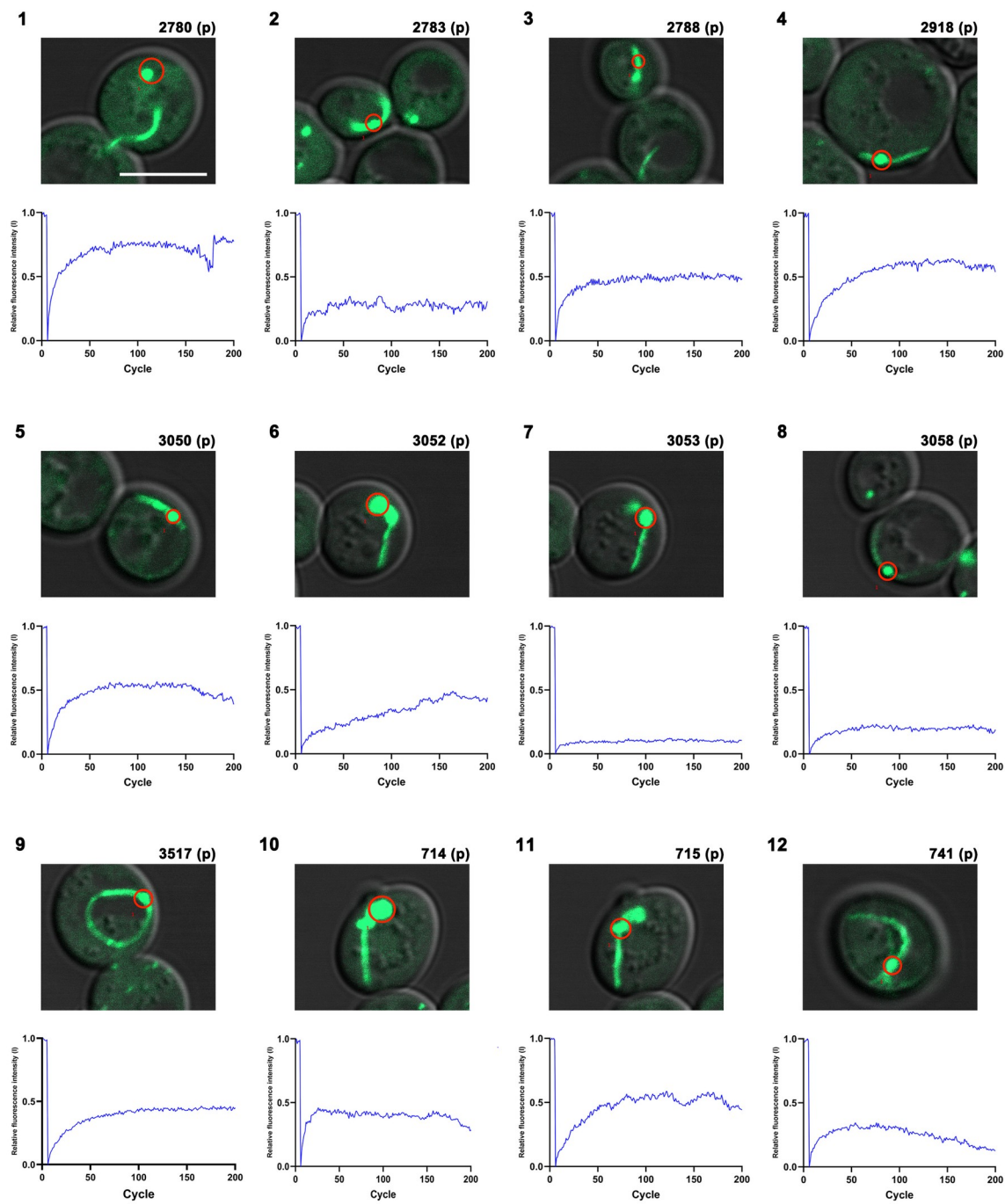

**Figure S2**

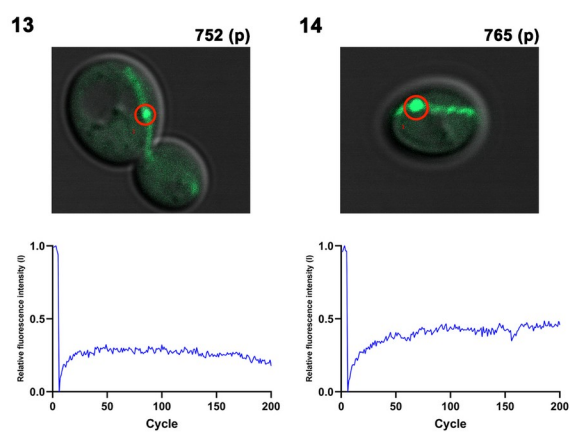

### Figure S2

D

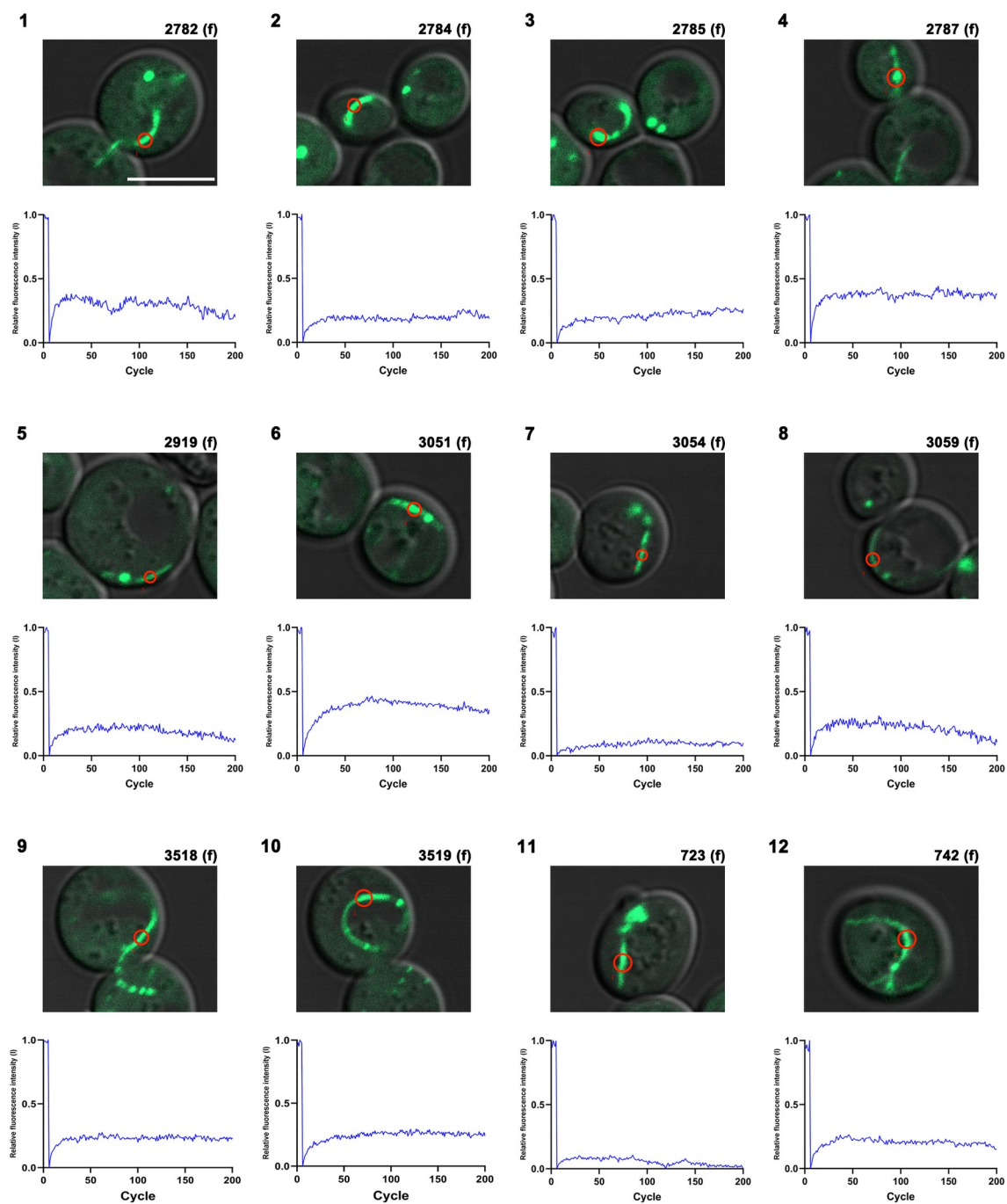

**Figure S2**

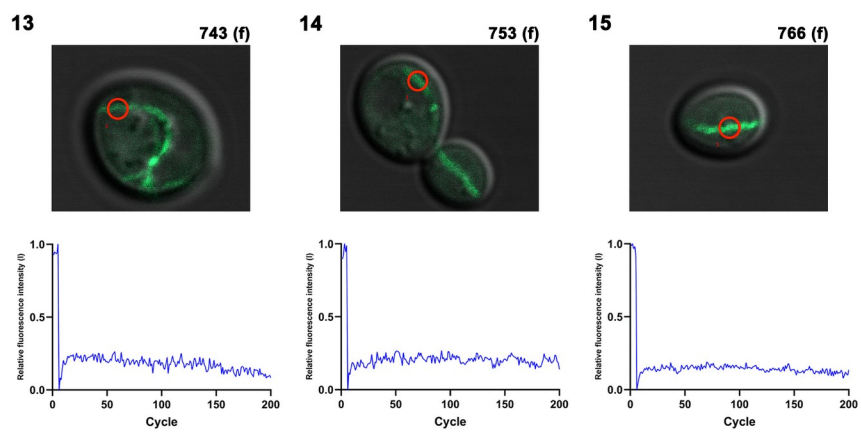

#### Figure S2

E

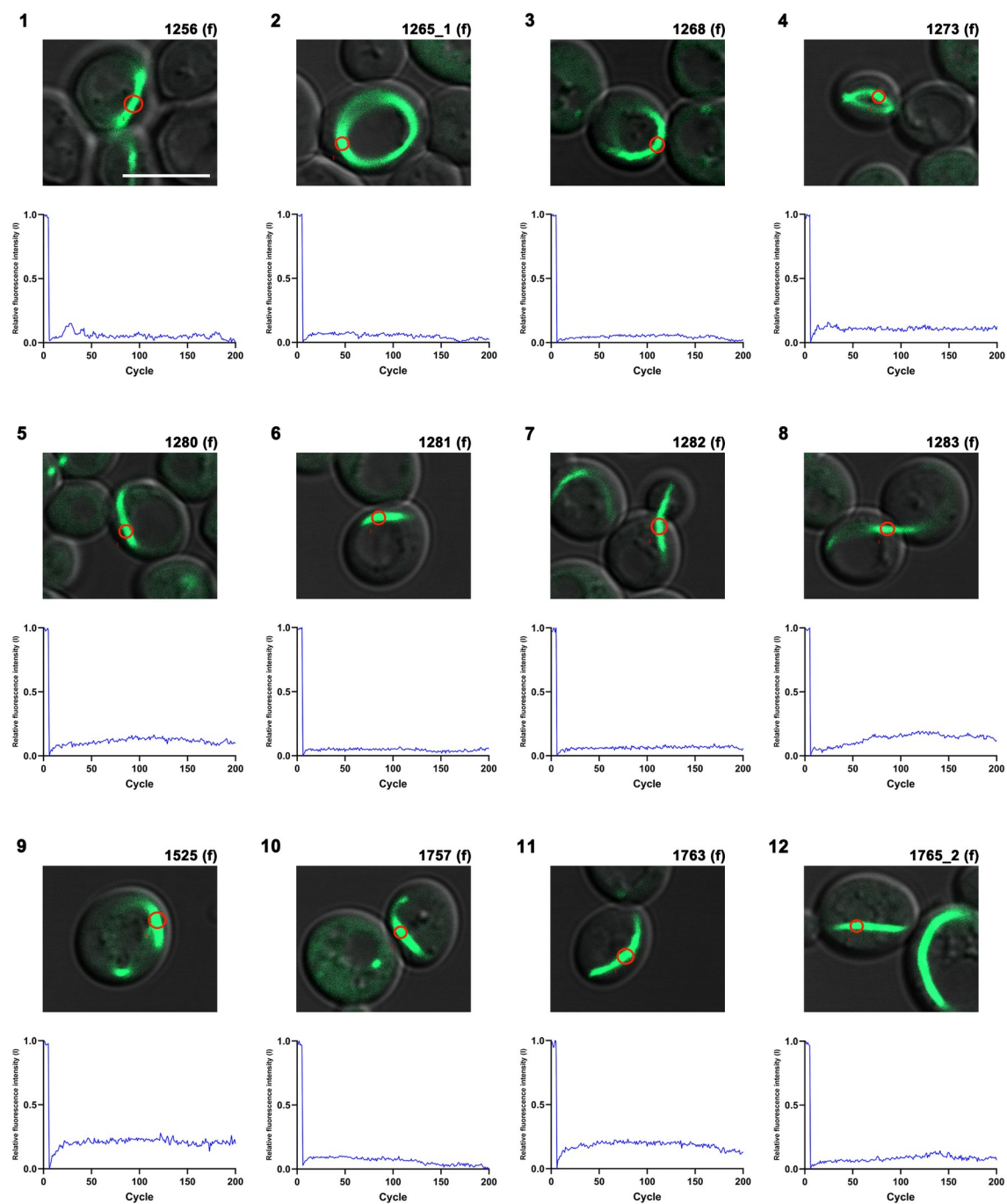

**Figure S2**

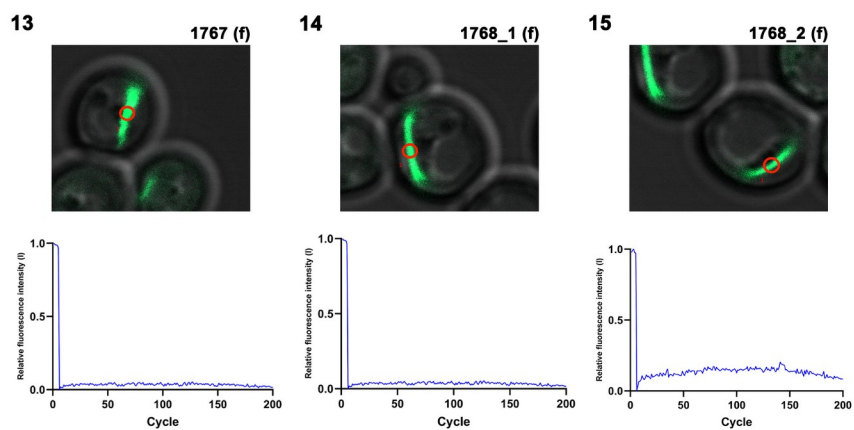

**Figure S2. Individual FRAP recovery curves for each morphological category of PAM 3R-E at 12 h, related to Figure 5**

Recovery curves of the individual structures that contribute to the averaged traces in Figure 5B, shown separately for each morphological category defined in Figure 5A: (1) puncta only, (2) connected puncta, (3-1) the punctate region and (3-2) the fibrillar region of a connected puncta–fibril structure, and (4) fibril only. Each panel plots every curve recorded for that category, with the mean overlaid. Yeast cells expressing GFP-TDP-43ΔNLS PAM 3R-E were induced with galactose for 12 h. n = 15, 16, 14, 15 and 15 structures for categories 1, 2, 3-1, 3-2 and 4. The x-axis is time (s) and the y-axis relative fluorescence intensity normalized to the pre-bleach value.

Figure S3

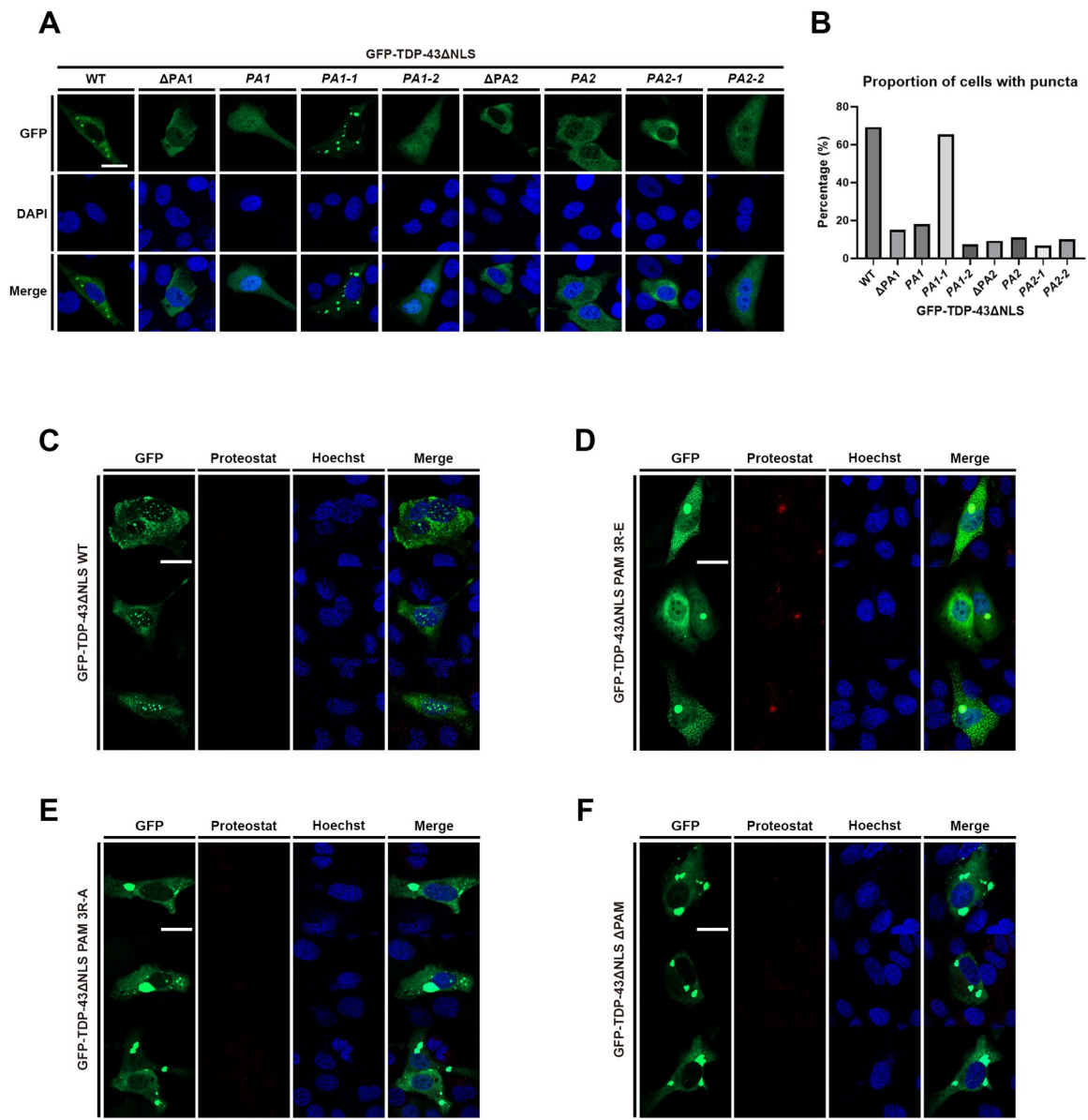

**Figure S3. Charged-cluster variants of TDP-43 in U2OS cells, related to Figure 6**

(A) Representative confocal images of U2OS cells expressing NLS-deficient GFP-tagged TDP-43 WT, ΔPA1, PA1, PA1-1, PA1-2, ΔPA2, PA2, PA2-1 and PA2-2, 48 h after transfection. GFP, DAPI and merged images are shown. Scale bar: 20 μm.

(B) Proportion of U2OS cells containing TDP-43 puncta for the constructs in (A). At least 50 cells were analyzed for each construct, from four independent experiments (n = 4).

(C–F) Representative confocal images of U2OS cells expressing (C) WT, (D) PAM 3R-E, (E) PAM 3R-A and (F) ΔPAM, 48 h after transfection, stained with ProteoStat and Hoechst. GFP, ProteoStat, Hoechst and merged images are shown. Scale bar: 20 μm.

**Figure S4**

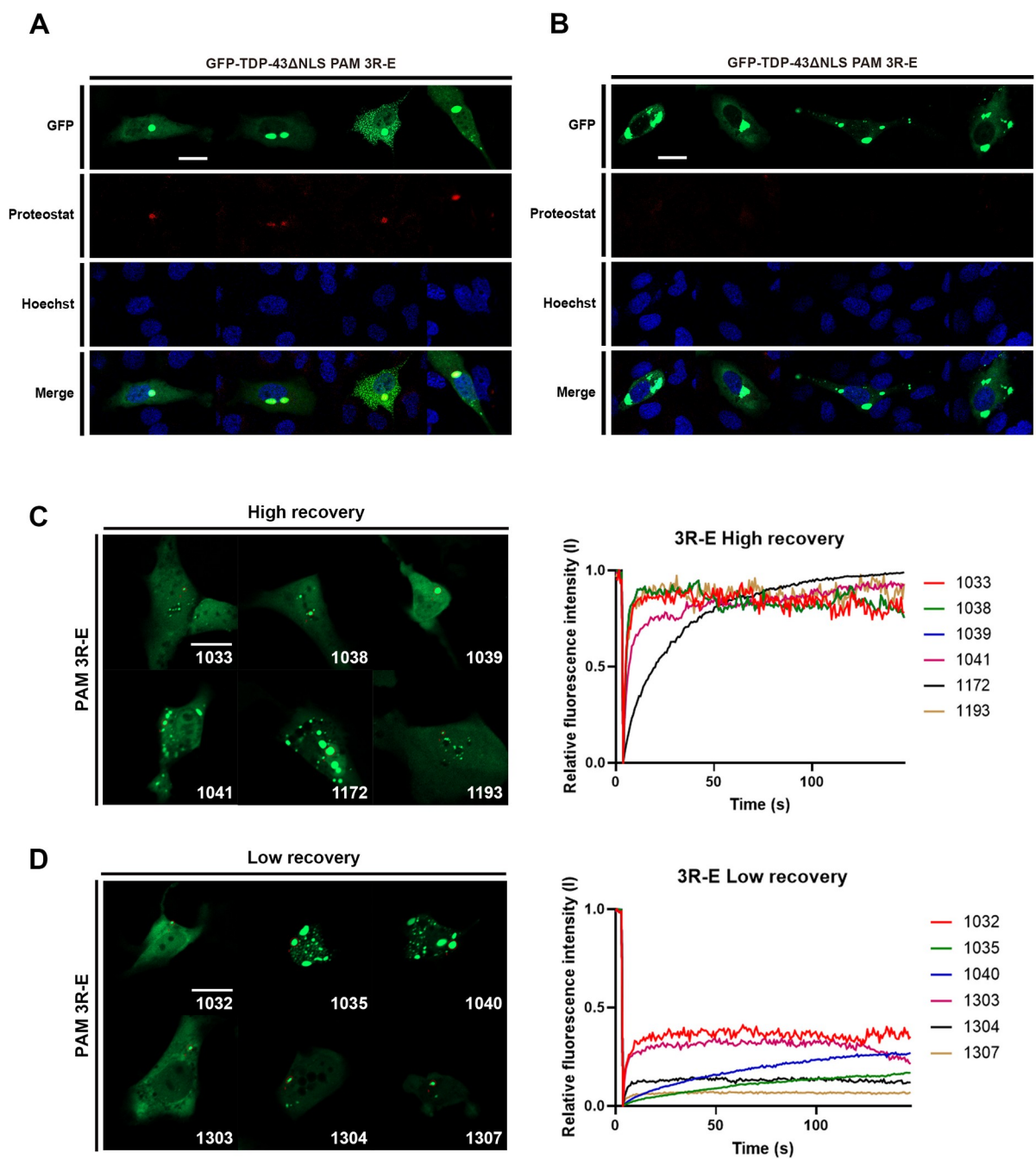

**Figure S4. ProteoStat reactivity and molecular exchange differ between individual condensates in U2OS cells expressing PAM 3R-E, related to Figure 6**

(A, B) Two representative fields of U2OS cells expressing NLS-deficient GFP-tagged TDP-43 PAM 3R-E, 48 h after transfection, stained with ProteoStat and Hoechst. GFP, ProteoStat, Hoechst and merged images are shown. Within the same field, condensates in some cells are ProteoStat-positive while those in neighboring cells are not. Scale bar: 20 μm.

(C, D) Representative fluorescence microscopy images and individual FRAP recovery curves of condensates in U2OS cells expressing NLS-deficient GFP-tagged TDP-43 PAM 3R-E, 48 h after transfection, showing (C) high and (D) low fluorescence recovery after photobleaching. Each curve corresponds to the individual condensate indicated in the image. The x-axis represents time in seconds and the y-axis the relative fluorescence intensity (I). Scale bar: 20 μm.

Figure S5

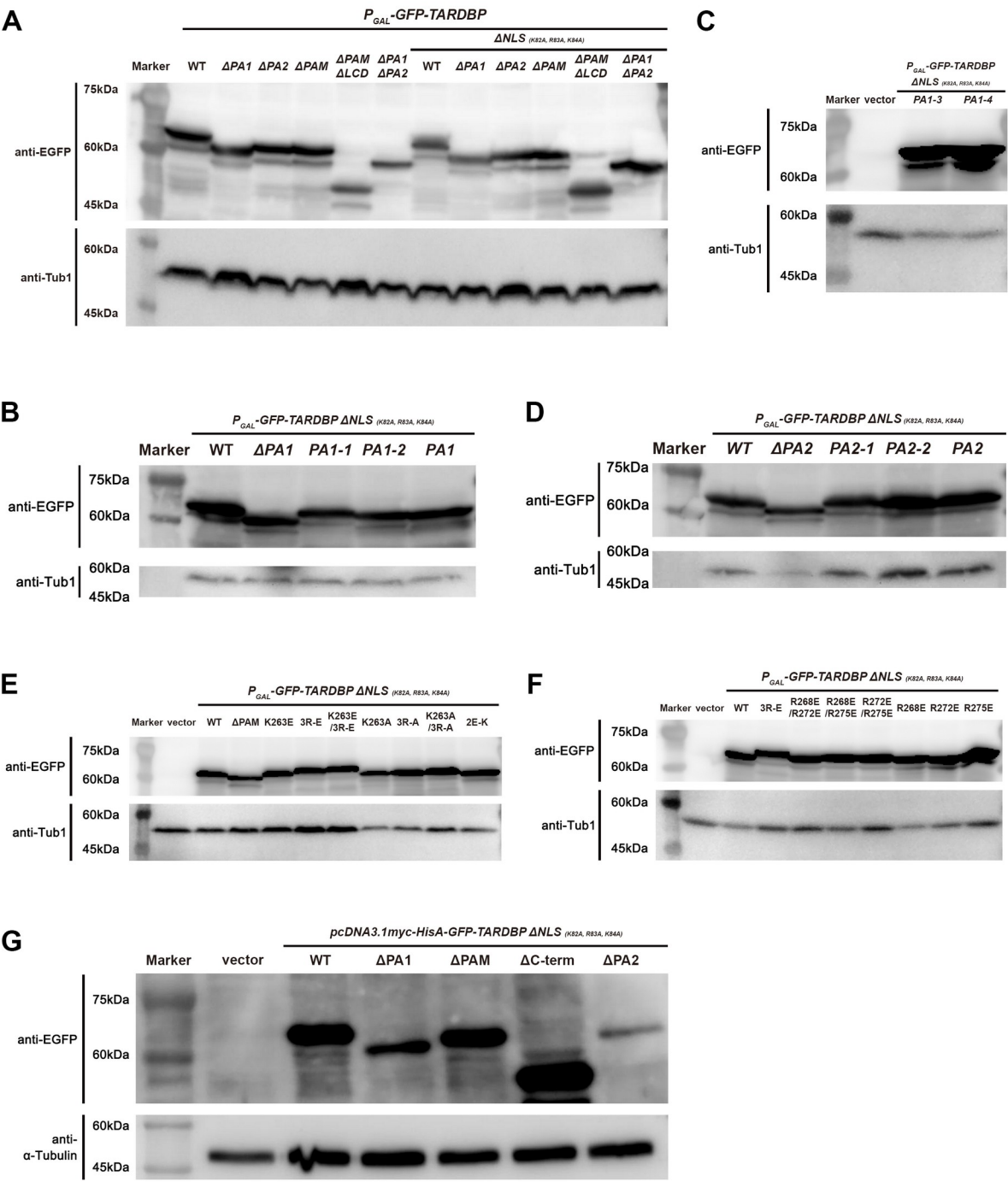

**Figure S5. Expression of GFP-tagged TDP-43 constructs, related to Figures 1, 2, 6 and 7**

Western blots confirming expression of the constructs used in this study, detected with anti-GFP. Yeast samples (A–F) were collected after 4 h of galactose induction and probed for Tub1 as a loading control. (A) GFP-TDP-43 and GFP-TDP-43ΔNLS domain deletion mutants in budding yeast. (B) PA1 charge-neutralization mutants in yeast. (C) PA1-3 and PA1-4 in yeast. (D) PA2 charge-neutralization mutants in yeast. (E) PAM charge-substitution mutants (K263E, PAM 3R-E, K263E/3R-E, K263A, PAM 3R-A, K263A/3R-A, 2E-K) in yeast. (F) Single and paired arginine substitutions (R268E/R272E, R268E/R275E, R272E/R275E, R268E, R272E, R275E) in yeast. (G) pcDNA3.1-based GFP-TDP-43ΔNLS constructs (WT, ΔPA1, ΔPAM, ΔPAMΔLCD, ΔPA2) in U2OS cells 48 h after transfection, probed for α-tubulin as a loading control.

**Video S1.** Three-dimensional rotation of the confocal z-stack reconstruction of the first cell shown in Figure 3A, related to Figure 3. A W303a yeast cell expressing GFP-TDP-43ΔNLS PAM 3R-E after 24 h of galactose induction, rendered as a three-dimensional volume and rotated through 360° about the vertical axis. Elongated curved filaments and discrete round puncta are both present within the same cell. Playback 24 frames s<sup>-1</sup>.

**Video S2.** As in Video S1, for the second cell shown in Figure 3A. Related to Figure 3.

**Video S3.** As in Video S1, for the third cell shown in Figure 3A. Related to Figure 3.

**Video S4.** As in Video S1, for the fourth cell shown in Figure 3A. Related to Figure 3.
